# PAF15-PCNA assembly exhaustion governs lagging strand replication and replisome integrity

**DOI:** 10.1101/2025.03.15.641049

**Authors:** Gita Chhetri, Sugith Babu Badugu, Narcis-Adrian Petriman, Mikkel Bo Petersen, Ganesha Pandian Pitchai, Aylin Seren Güller, Jan Novotný, Barath Balarasa, Morten Frendø Ebbesen, Frederik Tibert Larsen, Tina Ravnsborg, Anoop Kumar Yadav, Anita Lunding, Hana Polasek-Sedlackova, Ole Nørregaard Jensen, Kim Ravnskjaer, Jonathan R. Brewer, Jesper Grud Skat Madsen, Jens S. Andersen, Kumar Somyajit

## Abstract

Genome replication in eukaryotic cells is surveyed by the S-phase checkpoint, which orchestrates sequential replication origin activation to avoid exhaustion of hitherto poorly defined rate-limiting replisome components. Here, we find that excessive activation of replication origins depletes chromatin-bound PCNA and lagging strand components, thereby limiting additional PCNA loading at new origins when checkpoint control is disrupted. PAF15 (PCNA-associated factor 15) emerges as a dosage-sensitive regulator of PCNA, delineating the dynamic range of global genome duplication and defining distinct roles for PCNA on the leading and lagging strands. Through its high-affinity PIP motif and interaction within the DNA encircling channel of PCNA, PAF15 stabilizes PCNA exclusively on the lagging strand, optimizing and rate-limiting lagging strand processing. On the other hand, misregulation of PAF15—whether by overexpression or mislocalization to the leading strand—impairs replication fork progression and leads to cell death. These defects are mitigated by TIMELESS and CLASPIN, which restrain PAF15-PCNA interactions beyond the lagging strand. E2F4-mediated repression orchestrates PAF15 expression in normal and cancer cells, maintaining its optimal dosage for lagging strand-specific interactions with PCNA. Thus, the S-phase checkpoint functions in concert to restrict origin activation when lagging strand PAF15-PCNA assembly is exhausted, linking a previously concealed strand-specific rate limitation to overall replication dynamics.

---

Genome duplication is an essential process that ensures precise genetic inheritance and protects against genomic instability—a key driver of oncogenic transformation and cancer progression(*1, 2*). In eukaryotic cells, such precision hinges on two fundamental aspects: the regulation of the number of active replication origins and the velocity of active replisomes(*3*). The DNA replication program commences during the G1-phase with the licensing of an excess pool of replication origins, marked by the chromatin loading of inactive Minichromosome Maintenance (MCM) 2–7 hexameric replicative helicases. However, during S-phase, only a small subset (around 10%) of these origins is activated under stringent control to form active replisomes, characterized by the assembly of the MCM 2–7 complex with CDC45 and GINS1-4 (forming the active CMG helicase) alongside the DNA polymerases and layers of additional regulatory proteins(*4, 5*). The S-phase checkpoint, governed by the Ataxia Telangiectasia and Rad3-related (ATR) pathway coordinates replication origin firing with ongoing DNA synthesis during normal S phase as well as under replication stress to prevent premature or excessive origin activation and untimely mitotic entry(*6-8*). While insufficient origin activation results in DNA under-replication(*9*), excessive origin firing exhausts replication resources and destabilizes the genome (*10-12*). Thus, we hypothesize that the S-phase checkpoint operates within boundaries set by as-yet unidentified replisome dynamics, which might establish a global range of replication capacity and prevent excessive replication beyond this threshold.

## Unscheduled origin activation exhausts PCNA-DNA lagging-strand replication activities

To test this hypothesis, we investigated whether rapid origin firing in an S-phase checkpoint-deficient system would deplete core replisome components and thereby reveal the key rate-limiting steps of global genome replication. Therefore, we applied brief ATR inhibition to induce rapid and aberrant CDK activation and origin firing in a cell line expressing a TagRFP-chromobody targeting endogenous Proliferating Cell Nuclear Antigen (PCNA), a well-established proxy for active replisomes, to monitor the local replication dynamics in a large number of cell nuclei (*13*) (**Fig. 1A**). Employing quantitative image-based cytometry (QIBC) (*12, 14-16*), we first confirmed that short-term ATR inhibition (40 minutes) effectively unleashed CDK activity during S-phase (**Fig. 1A; Fig. S1A**) without inducing a DNA damage response, as indicated by the absence of pan-nuclear phosphorylated histone H2AX (γ-H2AX) (**Fig. S1B**). Such short-term ATR inhibition dramatically slowed replication fork speed (**Fig. S1C**) attributed to increased origin activation frequency, as evidenced by the single-stranded DNA replication bubbles marked by chromatin-bound Replication Protein A (RPA) in S-phase, while origin licensing, marked by MCM7 chromatin loading, remained unaltered (**Fig. S1D**). To our surprise, we did not observe any increase in PCNA chromobody foci upon ATR inhibition, as assessed through QIBC of both fixed cell population and live-cell tracking (**Fig. 1A**; **Fig. S1E**), despite the presence of elevated CDK activity and clear evidence of new origin firing. To independently validate the PCNA chromobody-specific readout, we used both an anti-PCNA antibody and GFP-tagged PCNA, yielding consistent results (**Fig. 1B, C**; **Fig. S2A-D**). ATR inhibition led to the focal accumulation of early replisome components—including CDC45 (a key member of the CMG complex), the replicative DNA polymerases POLε and POLδ, and the replication progression complex (RPC) factors TIMELESS and CLASPIN—on replicating chromatin (**Fig. S2B**). However, chromatin-bound PCNA did not correlate with the assembly of new replisomes (**Fig. 1B, C**; **Fig. S2A**).

**Figure 1:**
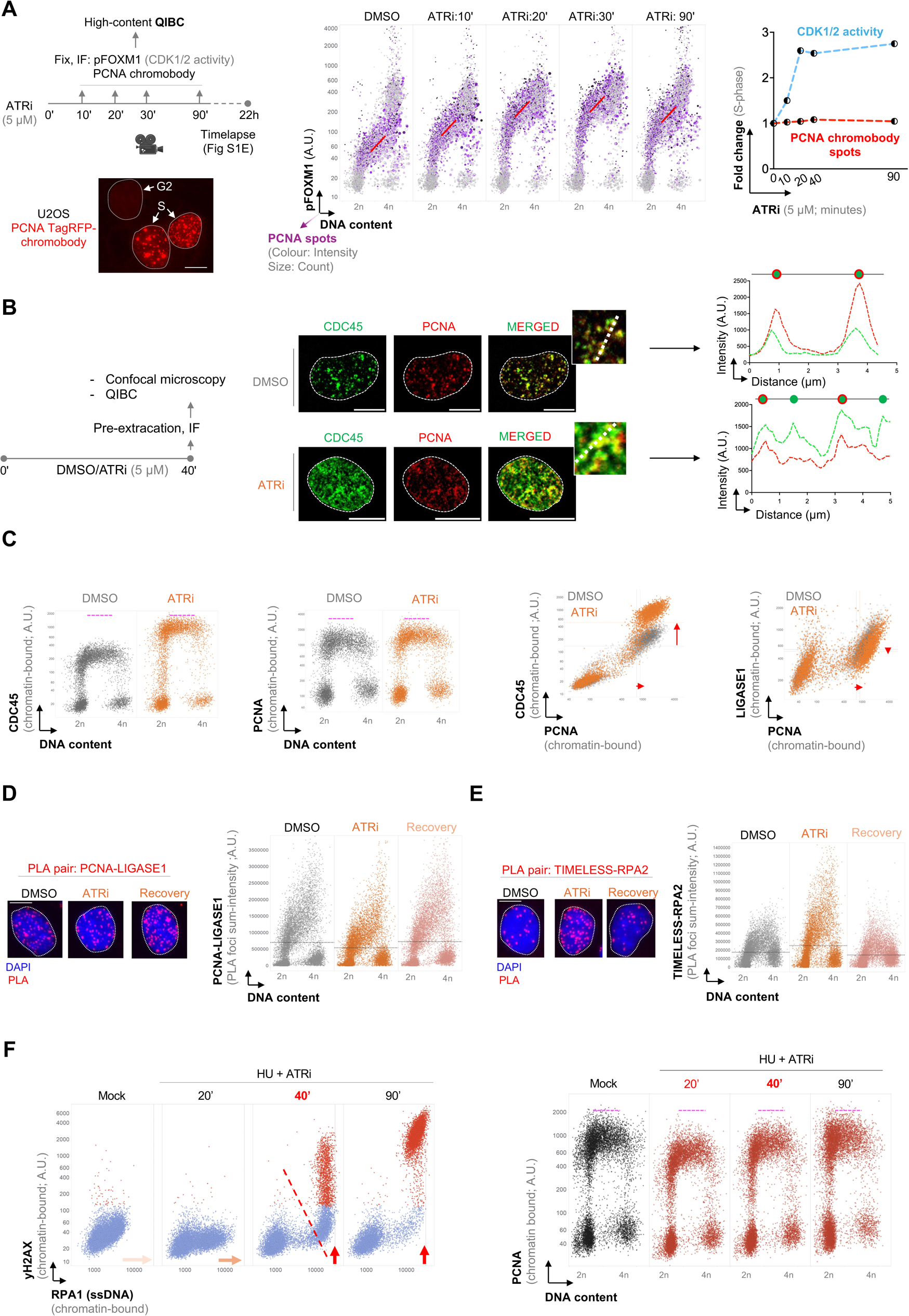
Excess replication origins drive PCNA-DNA lagging strand depletion. **(A)** Left: Experimental workflow employing U2OS cells stably expressing the TagRFP-PCNA chromobody, along with representative images of the PCNA profile during the S and G2 phases. Scale bars, 10 µm. ATRi: ATR inhibitor (AZD6738); pFOXM1: phospho-FoxM1 (threonine 600; a reliable proxy of CDK1/2 activities). Middle: QIBC of cells exposed to ATRi for the indicated time points, immunostained for pFOXM1. Nuclear DNA was counterstained with 4′,6-diamidino-2-phenylindole (DAPI). DNA content was derived from the total intensity of DAPI per cell (2n: G1, 4n: G2; > 8,000 cells per condition). The magenta color indicates the TagRFP-PCNA chromobody–positive population, and the size gradient indicates PCNA chromobody spots. All values are in arbitrary units (A.U.). Right: Fold change in CDK1/2 activities (as a function of mean intensity of pFOXM1) and PCNA chromobody spot count, derived from the QIBC data in the middle panel relative to the DMSO-treated condition. **(B)** Left: Experimental workflow. Middle: 3D confocal microscopy image of U2OS cells with endogenously tagged CDC45-GFP and immunostained for PCNA, exposed to ATRi as indicated. Scale bars: 10 µm. Right: Intensity line profiles showing the proximal localization of CDC45-GFP relative to PCNA within replication foci (from the insets in the middle panel). **(C)** QIBC of cells immunostained in (B) for the relative chromatin binding of CDC45, PCNA, and LIGASE1. n > 5,000 cells. Left: Representative images. Scale bars, 10 µm. Right: Quantification of the total PLA focus intensity per cell nucleus for the PLA pairs—PCNA and LIGASE1 **(D)** and TIMELESS and RPA2. The horizontal lines are medians. n > 10,000 cells. **(E)**. Cells were exposed to ATRi for 40 minutes, followed by 40 minutes of recovery, before being processed for PLA and QIBC. The horizontal line depicts the average values for 10,000 cells per condition. **(F)** U2OS cells were incubated with HU (2 mM) and ATRi (5 μM) for the indicated times and analyzed by QIBC to assess the chromatin binding of RPA1 and *γ*H2AX (left), and PCNA (right) from the same cells. 10,000 cells per condition. Left: Note that while RPA steadily accumulates (indicated by orange and red arrow), no DNA damage is detected before 40 minutes— damage becomes apparent when pan-nuclear, *γ*H2AX-positive cells (in red) emerge. The dotted line at the 40-minute time point indicates the onset of replication catastrophe. Right: PCNA does not accumulate on chromatin beyond the level observed in the mock condition in the same cells as in the left panel. Pink dotted lines in QIBC scatter plots indicate the approximate maximum levels of each protein.

PCNA, a highly abundant homotrimeric sliding clamp, acts as a structural scaffold that enables continuous leading strand synthesis by POLε(*17*) while coordinating the dynamic interplay among POLδ, FEN1, and LIGASE1 for lagging strand synthesis(*18*). Unlike the leading strand, lagging strand synthesis occurs in short Okazaki fragments (OkFs), necessitating a repetitive and time-intensive sequence of events(*19*). Strikingly, while the core and leading strand-specific factors were enriched (**Fig. S2B**), further analysis revealed that lagging strand processes dependent on PCNA—such as those involving FEN1 and LIGASE1—were specifically diminished during origin activation in checkpoint-compromised cells (**Fig. 1C**; **Fig. S2E-G**). This was consistently confirmed through biochemical chromatin fractionation (**Fig. S2F, G**), proximity ligation assay (PLA) coupled to QIBC (**Fig. 1D, E**; **Fig. S3A**), and QIBC across three independent cell lines (**Fig. S3B, C**), each with distinct replication dynamics(*14*). Moreover, these findings obtained by ATR inhibitor were fully recapitulated by both WEE1 inhibition and CLASPIN depletion, both of which enhance CDK activity and promote aberrant origin firing (**Fig. S3D-G**).

Unligated OkFs (as seen in FEN1-deficient cells) are processed via a non-canonical pathway, mediated by PARP1(*20, 21*). This is evidenced by the accumulation of nascent, S-phase-specific ADP-ribosylation chains following PARG inhibition (*20, 21*) (**Fig. S4A**). Aligned with our data showing that PCNA-lagging strand activities are inherently limited (**Fig. 1C, D**), we observed that even during normal DNA replication, PARG inhibition rapidly induces these chains specifically in S-phase (**Fig. S4B**). Moreover, acute ATR inhibition further amplifies this effect (**Fig. S4B**) by causing PCNA-LIGASE1 chromatin exhaustion. The critical importance of such non-canonical lagging strand maturation— particularly under conditions of excessive origin firing—is further underscored by the synthetic lethality observed when PARP1 inhibition is combined with ATR or WEE1 inhibition (**S4C, D**).

Premature origin firing and DNA replication stress have a catastrophic impact on the replicating genome, collectively defined as *replication catastrophe*(*11*). Exhaustion of the available pool of genome-protective RPA protein marks the onset of such terminal replication collapse, especially when checkpoint failure coincides with DNA replication stress induced by hydroxyurea (HU)-triggered nucleotide depletion (*12*). In light of our new findings, we compared the rate of RPA exhaustion with the chromatin paucity of PCNA. Intriguingly, PCNA and LIGASE1 are exhausted before RPA under HU treatment with ATR inhibition (**Fig. 1F**; **Fig. S4E**), suggesting that loss of core replisome activities possibly trigger replication catastrophe, which ultimately culminates when pathological fork DNA structures fail to be stabilized by the exhausted RPA pool.

Together, these results reinforce the notion that PCNA and lagging strand processes undergo natural depletion during unperturbed origin activation.

## Limited PAF15 pool and lagging-strand specific localization sustain PCNA stability and dynamics during unperturbed DNA replication

Despite its sheer abundance (*22*) and essential roles on both DNA strands, the potential exhaustion of PCNA—and its lagging strand activities—offers a unique window into the rate-limiting mechanisms that govern strand-specific replication dynamics.

To delve deeper, we combined mass spectrometry (MS) with TurboID-based biotinylation of PCNA-proximal proteins (**Fig. S5A-C**), identifying factors that may influence its rate-limiting mechanisms. We focused on PCNA loaders, including the canonical Replication Factor C1 (RFC1) and the CTF18-RFC variant—which encircle PCNA homo-trimers on the lagging and leading strands, respectively(*23, 24*)—and PCNA-associated factor 15 (PAF15, also known as PCLAF) (**Fig. S5C**). PAF15 contains a high-affinity PCNA-interacting peptide (PIP) motif and has been implicated in undergoing degradation during DNA damage, facilitating PCNA interaction with error-prone DNA polymerases(*25-28*), yet its role during unperturbed DNA replication remains unknown.

Mapping the dynamic range of origin activation following ATR inhibition, QIBC reveals that while CTF18 and RFC1 continues to accrue on replicating chromatin, PAF15 (like PCNA and LIGASE1) fail to accumulate compared to condition prior ATR inhibition, even as new origins continue to fire up to 60–90 minutes (**Fig. S5D**). This prompted us to investigate whether PCNA chromatin depletion is linked to a shortage of these factors in the soluble pool. Simultaneous quantitative, cell cycle-resolved measurements of the total and chromatin-bound fractions revealed that while PCNA, its effectors such as LIGASE1, DNA polymerases as well as CTF18 remained available, RFC1 exhibited a notable paucity in total levels (**Fig. 2A**; **Fig. S6A-E**). Strikingly, PAF15 was almost entirely sequestered on chromatin, with negligible detected excess protein during the exponential firing of replication origins (**Fig. 2B**). Such PAF15 paucity outside chromatin was confirmed across cell types and using antibodies targeting distinct endogenous protein epitopes, as well as through biochemical subcellular fractionation following G2/M synchronization and subsequent release into S-phase (**Fig. S6F-H**).

**Figure 2:**
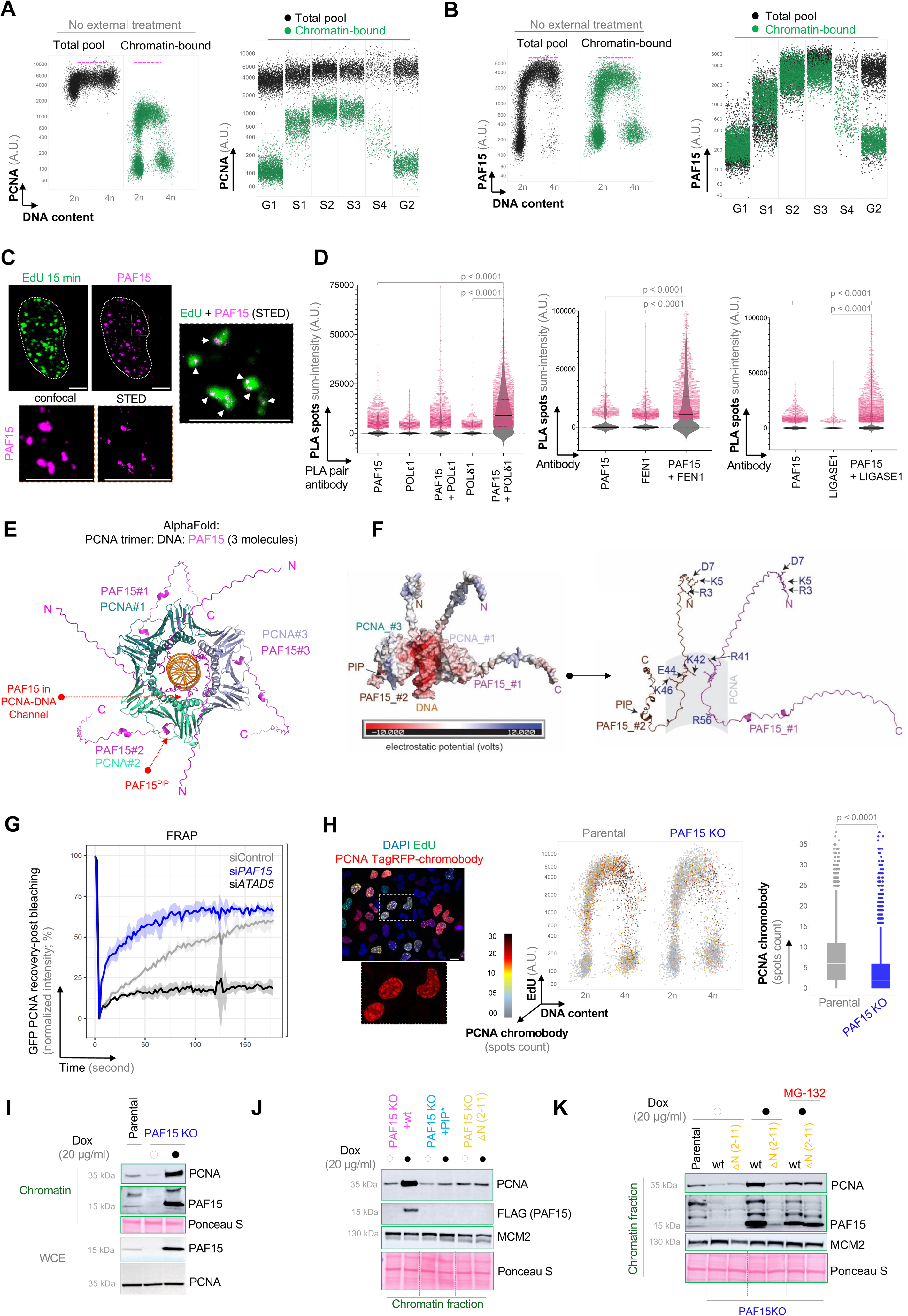
PAF15 is a dosage-limited, lagging strand-specific PCNA-stabilizing factor. QIBC-based side-by-side imaging and quantification of the total and chromatin-bound pool of PCNA **(A)** and PAF15 **(B)** in U2OS as well as their distribution in different phases of cell cycle. Cell cycle phases were stratified based on 5-ethynyl-2’-deoxyuridine (EdU) and DAPI. n > 10,000 cells. Pink dotted lines in QIBC scatter plots indicate the approximate maximum levels of each protein. **(C)** Confocal and STED microscopy images of cells stained for PAF15 and ongoing DNA replication with EdU. Scale bars, 5 µm. **(D)** Quantification of the sum of PLA focus intensity per cell nucleus obtained with indicated PLA antibody pair, PAF15-POLE1 and PAF15-POLD1 (left), PAF15-FEN1 (middle) and PAF15LIGASE1 (right). P values were determined by one-way ANOVA with Tukey’s test. **(E)** AlphaFold structural model of PCNA-PAF15-DNA complex. Red arrows indicate the PIP motif of PAF15 and traversal of the N-terminus of PAF15 to the DNA encircling channel of PCNA. **(F)** Adaptive Poisson-Boltzmann Solver (APBS) analysis of the modelled PCNA-PAF15-DNA complex formed by three PCNAs (one is removed for clarity), two PAF15 molecules and 19 nucleotides long DNA. Each PAF15 molecule contributes to the PCNA inner ring with four positive charges that interacts with DNA. The PCNA-PAF15-DNA complex is presented in side-view as cartoon representations (right) and surface representations of the electrostatic potential (left). To highlight the exposed charged surfaces inside of the PCNA ring, APBS is calculated for the full PCNA-PAF15-DNA complex as well as for models where individual components are systematically removed. The charged sidechains of PAF15 that contributes to the electrostatic potential within PCNA ring and at the N-terminus are indicated. **(G)** Fluorescence recovery after photobleaching (FRAP) of GFP tagged PCNA in U2OS with depleted PAF15 or known PCNA unloader, ATAD5. Cells were subjected to single bleach pulse and followed by the real-time recording of PCNA–GFP fluorescence recovery kinetics. Normalized PCNA–GFP FRAP curves reflect a gradual increase of the PCNA mobile fraction in PAF15 depleted cells. Each data point indicates mean ± s.d, n = 3 cells per condition. **(H)** Left: Representative image of TagRFP-PCNA chromobody with EdU and DAPI. Scale bars: 10 µm. Middle: QIBC of PCNA chromobody foci in U2OS parental and PAF15 KO cells. n > 10,000 cells. Right: Quantification of PCNA chromobody foci in EdU-positive S-phase cells from data in Middle. n > 4,000 cells. P values were determined by one-way ANOVA with Tukey’s test. **(I)** Protein levels of PCNA and PAF15 analyzed by western blotting (WB) in chromatin-bound and whole cell extracts (WCE) of U2OS parental or PAF15 KO with Doxycycline (Dox) inducible PAF15 wildtype expression. Cells were treated with Dox for 6 hours. **J)** WB analysis of PCNA in chromatin fraction of U2OS PAF15 KO with Dox inducible PAF15– either WT, mutated PIP motif or truncated N-terminal. **(K)** U2OS PAF15 KO cells with Dox inducible PAF15 were treated with Proteasome inhibitor MG-132 (10 µg) with Dox and chromatin fractions were analyzed by WB.

Of note, our findings on RFC1–PCNA exhaustion are consistent with a recent pre-print demonstrating the natural depletion of RFC1 under replication stress and checkpoint inhibition, conditions that impede fork restart (*29*). We focused on exploring PAF15 in greater depth, hypothesizing that its natural depletion might expose the rate-limiting mechanisms governing PCNA function after chromatin loading—particularly those involved in strand-specific DNA replication dynamics.

First, we confirmed that despite its low abundance, PAF15 is a *bona fide* component of the active replisome, as evidenced in large-scale iPOND (isolation of protein on nascent DNA)-MS (*14*)(**Fig. S7A**) and various imaging techniques (**Fig. 2C**; **Fig. S7B-D**). MS-based interactome analysis of chromatin-fractionated PAF15 suggested that PCNA is the primary partner responsible for recruiting PAF15 to the replisome (**Fig. S7E, F**). Consistently, the highly conserved PIP motif in PAF15 is fully essential for its localization to replicating chromatin (**Fig. S8A, B**). In contrast, compromised degradation of PAF15 during M/G1 by the anaphase-promoting complex/cyclosome (APC/C) ^Cdh1^—mediated via its KEN box (*25*)—does not lead to increased PAF15 loading (**Fig. S8B**), indicating that stabilization in G1/M does not affect its availability in S phase.

Because PCNA is essential for processive DNA replication on both strands, we next investigated which PCNA pool recruits PAF15 at the replisome. Using PLA to capture spatially proximal protein–protein interactions(*30*), we found that PAF15 specifically interacts with lagging strand components (POL δ, FEN1, LIGASE1), while being excluded from leading strand factors such as POL ε subunits (**Fig. 2D**; **Fig. S8C-E**). Although surprising, these results align with immunoblotting and MS-based analyses of chromatin fractions from various cell types, which consistently show higher levels of PCNA than PAF15 (**Fig. S8F, G).** These results suggest that PAF15 interacts with a specific subset of the PCNA pool, potentially directing PCNA to meet the distinct requirements of lagging strand synthesis, and that becomes exhausted during global replication.

PAF15 is an intrinsically disordered protein (see **Fig. S8A**). Crystallographic and NMR studies of its extended PIP-box suggest that upon binding to PCNA, PAF15 also accesses the DNA-encircling channel of PCNA (*28*)— although the functional significance of this engagement remains unclear. To this end, we employed AlphaFold modelling (*31*) to analyze full-length PAF15 in complex with the PCNA homotrimer on DNA, suggesting that PAF15 acts as a molecular wedge that completely traverses the PCNA ring, exhibiting interactions completely distinct from that of other PIP-box proteins (**Fig. 2E**; **Fig. S9A-C**).

An Adaptive Poisson-Boltzmann Solver (APBS) analysis(*32*) of the modelled PCNA– PAF15–DNA complex revealed that each PAF15 molecule contributes with positively charged residues to both the inner PCNA ring and its N-terminus, likely stabilizing its interaction with DNA (**Fig. 2F**). Given that purified human PCNA exhibits weak DNA affinity *in vitro* and high diffusion rates(*33*), we hypothesized that, owing to its unique interaction within the PCNA ring, PAF15 might enhance PCNA–DNA contacts. Consistent with this hypothesis, PAF15 depletion resulted in a remarkable increase in the PCNA exchange rate and reduced its chromatin residence time, as revealed by fluorescence recovery after photobleaching (FRAP) (**Fig. 2G**). Consistently, reduced PCNA on chromatin in the absence of PAF15 was confirmed via QIBC of the PCNA-TagRFP chromobody (**Fig. 2H**) and by immunoblotting of chromatin fractions across various cell types (**Fig. 2I**; **Fig. S10A, B**). To examine PCNA stability following rapid PAF15 loss, we tagged endogenous PAF15 at its C-terminus with an auxin-induced degron (AID) and GFP. However, tagged PAF15 failed to fully mimic the endogenous protein, exhibiting impaired chromatin loading (**Fig. S10C**). The GFP tag likely disrupts PAF15’s small structure and its dual interactions with both sides of the PCNA ring, thereby compromising its function (**Fig. 2E**). Nevertheless, this resulted in the generation of hypomorphic PAF15 cells with dramatically reduced PCNA chromatin stability, an effect further exacerbated by auxin-induced PAF15 degradation (**Fig. S10C**).

Finally, the role of PAF15 in promoting PCNA chromatin stability was directly validated by short-term inducible overexpression of PAF15 in KO cells, which rapidly stabilized PCNA on chromatin—a response absent when using the PIP mutant (**Fig. 2I, J**; **Fig. S10D**).

To gain further insights, we revisited the APBS analysis, which revealed that in addition to the positively charged residues near the PIP motif (potentially influencing the PIP peptide), PAF15 harbours additional positively charged residues at its Histone H3–like N-terminus (see **Fig. 2F**). These residues engage the backside of PCNA and, by closely interacting with DNA, may stabilize the entire PAF15–PCNA–DNA complex (**Fig. S11A**). To this end, we generated a series of PAF15 N-terminal truncation mutants. Notably, deleting the first 11 residues—encompassing the positively charged region—was sufficient to severely compromise PAF15 protein stability (**Fig. S11B, C**) and, consequently, PCNA retention on chromatin (**Fig. 2J**; **S11D**). However, restoring the stability of the truncated, yet PIP box– containing, form of PAF15 via proteasomal inhibition rescued PCNA stability on chromatin (**Fig. 2K**; **S11D**), altogether establishing PAF15 as a direct regulator of PCNA stability and dynamics on DNA.

Notably, these functions of PAF15 were independent of the ubiquitin ligase UHRF1, which has been co-crystallized with the PAF15 N-terminus and is known to mono-ubiquitinate PAF15 at K15 and K24 (PAF15Ub2) facilitating its degradation upon DNA damage(*34, 35*). Although UHRF1 depletion abolished PAF15 ubiquitination, it did not affect PAF15 loading onto replicating chromatin with PCNA (**Fig. S11E, F**). This observation aligns with the AlphaFold model, which predicts that UHRF1 binding does not interfere with the PIP-mediated interaction between PAF15 and PCNA (**Fig. S11G, H**).

## PAF15 safeguards Okazaki fragment processing and mitigates replication stress

We next examined whether the dramatic loss of chromatin-bound PCNA in the absence of PAF15 leads to alterations in replisome architecture. Surprisingly, MS analysis of PCNA TurboID revealed that PAF15 deletion selectively disrupts PCNA interactions with lagging strand factors, while other key replisome-associated PCNA interactions remain unaltered (**Fig. 3A**). Chromatin fractionation confirmed that although PAF15 loss destabilizes PCNA, the core replisome remains intact (**Fig. S12A**). Instead, the chromatin-bound fraction of lagging strand maturation protein—such as FEN1 and LIGASE1—were selectively reduced (**S12A-C**). This was further validated in thousands of S-phase cells using PLA-QIBC, which demonstrated significantly diminished interactions between PCNA and FEN1/LIGASE1 (**Fig. 3B**; **S12D, E**). Reconciling these, impaired OkF maturation in PAF15-deficient cells was directly evidenced by the accumulation of alkali-sensitive unligated Okazaki fragments (**Fig. 3C**; **S12F, G**).

**Figure 3:**
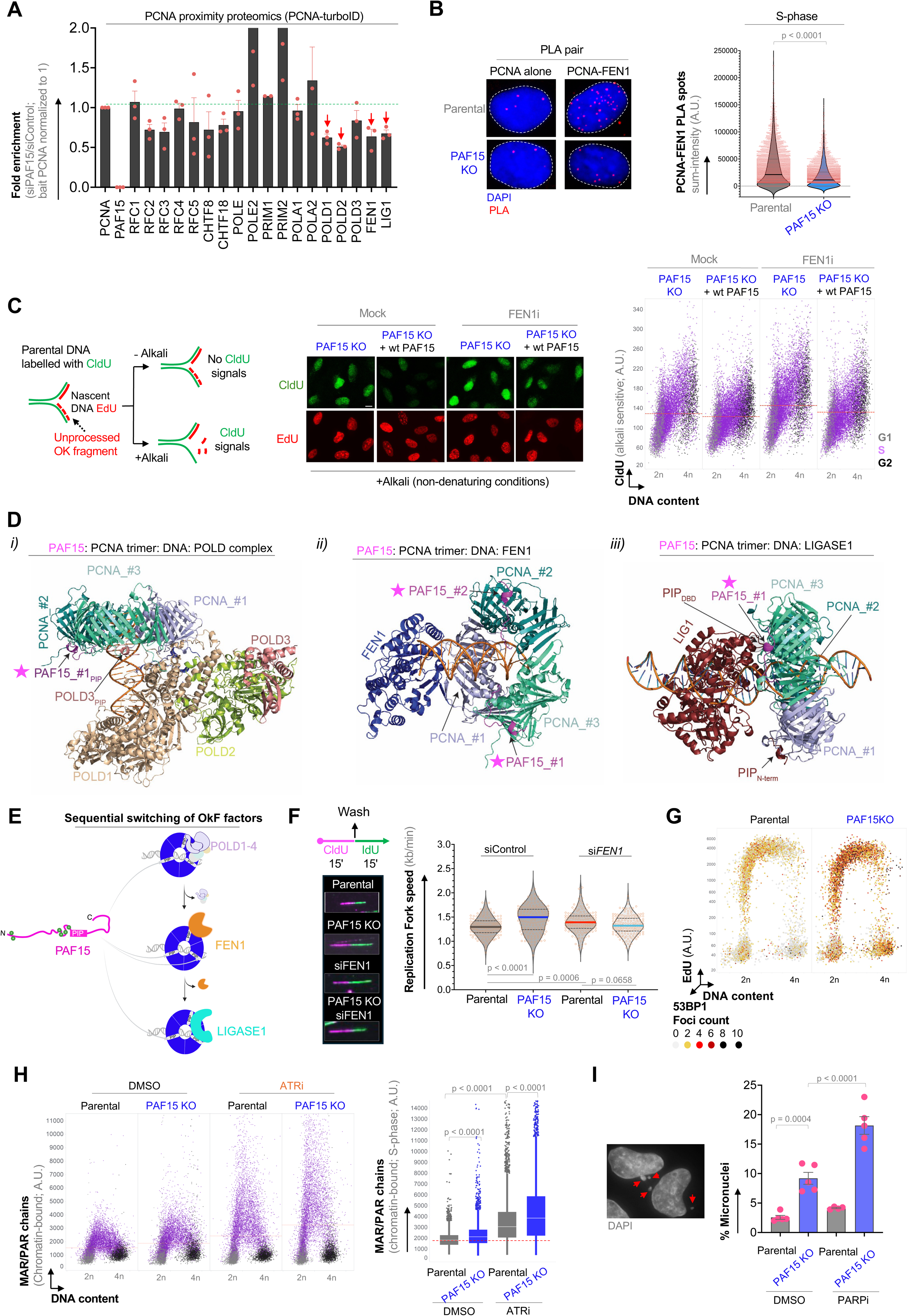
PAF15 shapes PCNA activity during Okazaki fragment maturation. **(A)** Proximity-based proteomic analysis turboID-PCNA in U2OS in PAF15 depleted cells. PCNA was normalized to 1 in both siControl and siPAF15 cells. **(B)** Left: Representative images. Scale bars, 10 µm. Right: Quantification of the sum of PLA focus intensity per cell nucleus obtained with PLA antibody pair: PCNA-FEN1 in U2OS parental and PAF15 KO cells. n > 5,000 S-phase cells per condition. P values were determined by one-way ANOVA with Tukey’s test. **(C)** left: Schematic visualization of the dissociation of Okazaki fragments from the nascent DNA (EdU) under Alkali treatment. Middle: Representative images. Right: QIBC of U2OS PAF15KO and PAF15 KO cells complemented with constitutive expression of wildtype PAF15 under FEN1 inhibition (FEN1i; 20 µM). n > 10,000 cells per condition. **(D)** AlphaFold visualization of the binding of i) PCNA-DNA-POLD1-4 complex, ii) PCNA-DNA-FEN1 and iii) PCNA-DNA-LIGASE1 complexes with PAF15. Alphafold folds a PCNA trimer-PAF15-POLD holocomplex (1-4)-DNA (left where PAF15 occupies one PCNA monomer whilst POLD1 and POLD3 saturates the other two. The structural model of a PCNA trimer -PAF15-FEN1-DNA shows that docking of FEN1 to one PCNA monomer allows PAF15 two occupy the other two. AlphaFold suggests a PCNA trimer-PAF15-LIG-DNA complex where LIGASE1 binds the PCNA trimer by its PIP_N-term_ and PIP_DBD_ domains allowing one PAF15 molecule to access the unoccupied PCNA. **(E)** Schematic visualization of the sequential binding of PCNA to POLD, FEN1 and LIGASE1 on lagging strand during maturation of Okazaki fragments based on AlphaFold in E. **(F)** Left: DNA fiber labelling protocol and representative images. CldU, 5-chloro-2′-deoxyuridine; IdU, 5-iodo-2′-deoxyuridine. Right: Replication speed of parental and PAF15KO with FEN1 depletion. n = 200 fibers per condition. P values were determined by one-way ANOVA with Tukey’s test. **(G)** QIBC of 53BP1 foci/nuclear bodies in indicated cells and labeled for EdU and DAPI to stratify cell cycle progression (n > 10,000 cells for each condition; colors indicate the number of 53BP1 nuclear bodies per nucleus. **(H)** QIBC analysis of the formation of mono-or poly-ADP ribosylated (MAR/PAR) chains in U2OS PAF15 KO under ATR inhibition. n > 10,000 cells per condition. Quantification in S-phase (5,000 cells per condition) shown on the right. P values were determined by one-way ANOVA with Tukey’s test. **(I)** Quantification of micronuclei formation in in indicated cells treated with PARP1 inhibitor Olaparib (500 nM, 24 h). 500 nuclei per condition. Values denote mean ± s.d. P values were determined by one-way ANOVA with Tukey’s test.

In addition to facilitating the processivity of replicative polymerases, PCNA also drives the activity of error-prone DNA polymerases—a process that requires PCNA mono-ubiquitination as a key signal for error-prone synthesis at sites of DNA damage(*36*). Surprisingly, the loss of PAF15, which is accompanied by compromised PCNA chromatin stability, failed to alter PCNA mono-ubiquitination upon exposure to the alkylating agent methyl methane sulfonate (MMS). This suggests that PAF15-mediated PCNA stability on replicating chromatin is independent of PCNA regulation in error-prone synthesis at DNA template lesions and is consistent with the fact that DNA damage triggers PAF15 degradation(*26*) (**S12H**).

PCNA, as a homo-trimer, orchestrates the activities of POL δ, FEN1, and LIGASE1 on DNA by acting as a dynamic toolbelt(*18, 24, 37*). However, the precise regulation of lagging strand substrate hand-off remains unclear, particularly how steric clashes are avoided when POLδ holocomplex (POLδ1-4), FEN1, and LIGASE1 compete for PCNA trimer in human cells. In fact, purified human PCNA–POL δ–FEN1 complexes on DNA—lacking PAF15—exhibit reduced stability and impaired substrate hand-off compared to yeast complexes, which naturally lack PAF15(*28, 38*). Therefore, we investigated whether the high-affinity interaction between PCNA and PAF15 is compatible with the PCNA-based switching of factors required for OkF maturation.

An AlphaFold model of the PCNA homotrimer–POLδ1–FEN1–LIGASE1–DNA complex (**Fig. S13A**) revealed an incompatible configuration with even a single PAF15 molecule (**Fig. S13B**). In this altered arrangement, PAF15 and LIGASE1 fold away from the PCNA ring, while LIGASE1’s PIP displaces FEN1, likely disrupting OkF processing (**Fig. S13B**). In sharp contrast, a model where POLδ1-4, FEN1, and LIGASE1 occupy PCNA subunits sequentially remains fully compatible with one or two PAF15 molecules (**Fig. 3D**; **Fig. S14A-C**). In such scenario, a constant association of PAF15 with PCNA monomers during OkF maturation not only would anchor PCNA on DNA but also likely enable a sequential exchange of lagging strand factors, preventing steric clashes among competing OkF proteins (**Fig. 3E**).

Indeed, supporting the notion that PAF15 operates in concert with OkF proteins, single-molecule DNA fiber analysis revealed that cells lacking PAF15 exhibit a significant increase in fork speed—independent of cell type or genetic ablation method (**Fig. 3F**; **Fig. S15A-D**). However, these cells also show replication fork asymmetry (**Fig. S15B, E**), a feature reminiscent of defective lagging strand maturation (*39*). PAF15 deletion combined with siRNA-mediated FEN1 depletion did not enhance fork acceleration and fork asymmetry beyond FEN1 ablated cells (**Fig. 3F**; **Fig. S15E**), confirming a cooperative role for PAF15 and FEN1 at the replisome. Failed OkF maturation in PAF15-deficient cells was accompanied 53BP1 nuclear body formation in G1 (**Fig. 3G**; **Fig. S16A-C**), micronuclei appearance (**Fig. S16D**), increased S-phase phosphorylation of RPA2 (pRPA2S33) (**Fig. S16E**), and reduced CDK activity (**Fig. S16F**)—each a hallmark of replication-associated DNA damage. Under these conditions, PAF15-deficient cells, both under normal replication and during checkpoint failure, resorted to a PARP1-dependent non-canonical OkF maturation pathway, as indicated by a significant increase in S-phase-specific ADP-ribosylation (**Fig. 3H**). Consequently, these cells accumulated lethal levels of replication stress when treated with PARP inhibitors (**Fig. 3I**; **Fig. S16G**).

Together, PAF15 stabilizes PCNA on chromatin to orchestrate canonical OkF maturation and suppress replication stress, while its depletion or natural exhaustion during excess origin firing disrupts lagging strand processing, enforcing a critical dependence on backup pathways.

## An excess of PAF15 or its binding to leading-strand PCNA is lethal

Given the strong impact of excess PAF15 on PCNA chromatin stability, we next investigated whether surplus PAF15 could overcome PCNA chromatin exhaustion during increased origin firing. Strikingly, excess PAF15 sequesters PCNA into PAF15–PCNA complexes, stabilizing PCNA while rendering it unavailable to support new replication origins upon replication checkpoint inhibition (**Fig. S17A**). Although this chromatin-bound PCNA initially appeared to rescue replication catastrophe, it ultimately resulted in a profound blockade of global DNA damage signaling (**Fig. S17B-D**). Consistently, over-stabilization of the PAF15–PCNA chromatin complex severely compromised DNA replication at multiple levels, culminating in complete cell death—a phenotype that was entirely rescued in variants harbouring either a mutated PIP motif or intrinsically unstable N-terminal truncated variants of PAF15 (**Fig. 4A-C**; **Fig. S17E-G, S18A**).

**Figure 4:**
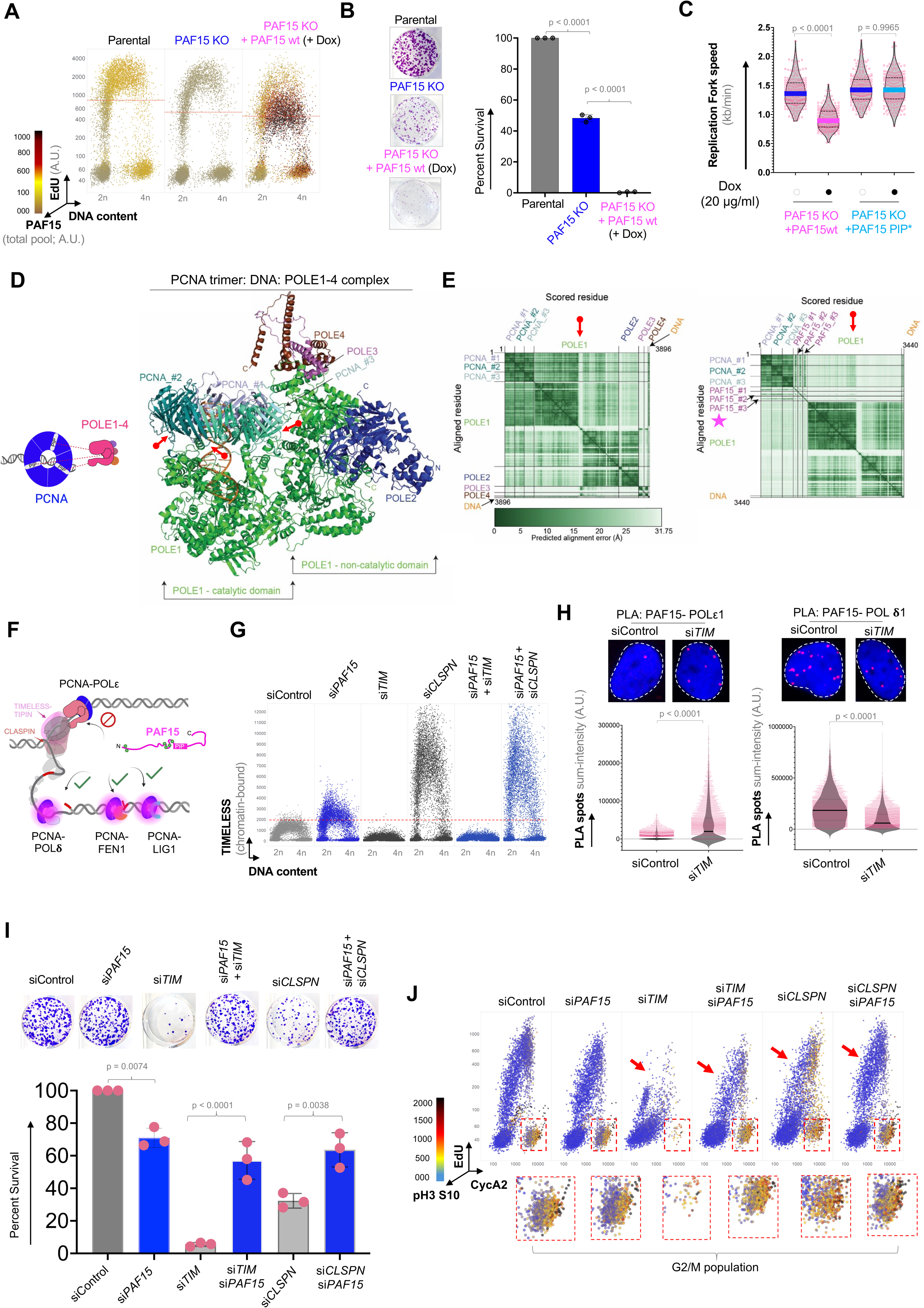
PAF15 access to leading strand PCNA is lethal. **(A)** QIBC analysis of the global DNA replication rate (EdU) and the total pool of PAF15 in U2OS parental, PAF15 KO and PAF15 KO cells treated with 20 µg/ml doxycycline to induce wildtype PAF15 for 24 hours. **(B)** Relative plating efficiency after 10 days of plating cells as treated in panel A. Values denote mean ± s.d. P values were determined by one-way ANOVA with Tukey’s test. **(C)** Replication fork speed in U2OS PAF15 KO treated with 20 µg/ml doxycycline to induce wildtype PAF15 or PIP motif mutated PAF15. n = 200 fibers per condition. P values were determined by one-way ANOVA with Tukey’s test. **(D)** Left: Hypothetical model. Right: The AlphaFold structural model of the octameric PCNA-POLE1-POLE2-POLE3-POLE4-DNA complex, showing shows that only POLE1 is accessing the PCNA ring. POLE2, POLE3 and POLE4 are docking onto the non-catalytical domain of POLE1 of the PCNA-DNA-POLE (1-4) complex. **(E)** AlphaFold modeling suggest mutual exclusive interaction of PAF15 and POLE1. The structural model of the PCNA-PAF15-POLE1-DNA complex shows that three molecules of PAF15 are displacing POLE1 from PCNA. This is evident when comparing the PAE scores (red arrow) of PCNA-POLE1 interaction in complexes containing PAF15 (right) with the PAE scores of complexes lacking PAF15 (left): The low PAE scores (<5Å) at the interaction interfaces of the model depicted in D) show high confidence in the structural model. **(F)** Model depicting PAF15 exclusion from leading strand. **(G)** QIBC analysis of chromatin-bound TIMELESS under the depletion of PAF15, TIMELESS (TIM) and CLASPIN (CLSPN), or their combination. A total of 5,000 cells were analyzed per condition. **(H)** Top: Representative images. Bottom: Quantification of the sum of PLA focus intensity per cell nucleus obtained with antibodies pairs, PAF15-POLE1 and PAF15-POLD1 under control or TIMELESS depletion. P values were determined by one-way ANOVA with Tukey’s test. **(I)** Relative plating efficiency of U2OS cells treated with indicated siRNAs after 10 days of plating cells. Values denote mean ± s.d. P values were determined by one-way ANOVA with Tukey’s test. **(J)** QIBC analysis of S-phase (EdU), CyclinA2 (CycA2) and mitotic (phosphorylated serine 10 on histone 3 – pH3 S10) fraction of U2OS cells treated with indicated siRNAs. A total of 10,000 cells were analyzed per condition. Red arrows indicate S-phase population revival in TIMELESS or CLASPIN depletion together with PAF15. Insets below depict G2/M cell population, which is restored in TIMELESS and PAF15 double depletion.

The drastic stabilization of the PAF15–PCNA chromatin complex raised the question of whether this effect resulted from doxycycline-induced PAF15 overexpression or whether it also reflected disruptions to the natural dynamics of PCNA on chromatin. Reassuringly, similar outcomes were observed following ATAD5 depletion—the natural PCNA unloader(*40*)—which immobilizes PCNA in complex with PAF15 (**Fig. S18B, C**). Eliminating PAF15 in ATAD5-depleted cells significantly reduced cell death (**Fig. S18D**), underscoring that only a limiting pool of PAF15 is required to maintain normal PCNA turnover, while any excess provokes a lethal replication response.

To underscore the lethal effect of aberrant PAF15, we examined the action of PCNA on the leading strand. In contrast to its role in mediating dynamic interactions with lagging strand factors, PCNA establishes a stable three-point contact on the leading strand, with the PIP box affinity site of each monomer directly engaging the POL ε1 subunit (**Fig. 4D**) (*41, 42*). Recent studies have established the equally important role of PCNA in leading strand progression within the human replisome(*17*)—a function previously thought to be confined to DNA damage tolerance(*43*). Strikingly, the AlphaFold model indicates that even a single molecule of PAF15 can disrupt the PCNA–POL ε interaction (**Fig. 4D, E**), explaining the massive impediment of the fork progression (see **Fig. 4C**; **Fig. S17E, F**).

To mechanistically address this further, we investigated whether, under normal conditions, factors at the leading strand protect POL ε-engaged PCNA from PAF15 while allowing PAF15 to interact with a more dynamic pool of PCNA on the lagging strand (**Fig. 4F**). In our search for these, we found that TIMELESS and CLASPIN—RPC components that couple several key replisome interactions, including linking POL ε to the CMG helicase(*44*) —play a crucial role in shielding leading strand-bound PCNA from PAF15.

First, loss of PAF15 enhanced the chromatin binding of the TIMELESS–CLASPIN complex—with TIMELESS being upstream of CLASPIN for chromatin loading **(Fig. 4G**; **Fig. S19A)**. Second, the increased chromatin binding of the TIMELESS–CLASPIN complex after PAF15 loss is driven by PCNA, as it was reversed by the PCNA inhibitor T2AA(*45*), which mirrors the PCNA binding properties of the PAF15 PIP motif **(Fig. S19B, C)**.

Consistent with data in mouse cells showing that CLASPIN contains a PIP motif and interacts with PCNA(*46*), human CLASPIN residues 304–341, which harbor the PIP motif, and are positioned closest to PCNA **(Fig. S19D, E)**. In this scenario, TIMELESS-CLASPIN may transiently shield PCNA during its loading onto the leading strand.

Supporting this, indeed removal of TIMELESS resulted in mislocalization of PAF15 to the leading strand (**Fig. 4H**; **Fig. S19F, G**) and triggered cell cycle arrest and cell death. Remarkably, these lethal phenotypes were robustly reversed when PAF15 was simultaneously removed across various cell types or ablation methods (**Fig. 4I, J**; **Fig. S20A-C**). The specificity of TIMELESS-CLASPIN-mediated protection of leading strand PCNA from PAF15 is further underscored by the observation that CLASPIN loss sensitizes CTF18-ablated cells—a sensitivity that is rescued by the simultaneous removal of CLASPIN and PAF15 (**Fig. S20D, E**).

Further structure-function studies involving strand-specific PCNA loaders(*23, 47*), as well as the TIMELESS-CLASPIN complex, are needed to elucidate the mechanism of PCNA delivery and functions at both replication strands.

Together, these results illuminate the role of TIMELESS-CLASPIN in shielding cells from the detrimental effects of misdirected PAF15 to leading strands and further highlight the critical need for strict regulation of the total PAF15 pool.

## E2F4-medited gene repression program rate-limits PAF15 dosage and genome integrity

PAF15 is an oncoprotein that is overexpressed in nearly every tumor type, as demonstrated by Pan-Cancer Atlas data (**Fig. 5A**). Moreover, elevated PAF15 levels correlate with reduced overall survival, exerting an even more pronounced impact than PCNA overexpression (**Fig. 5B**). Consistently, comparison of PAF15 levels in normal and cancer-derived cell lines revealed marked overexpression in the latter (**Fig. 5C**; **Fig. S21A**). When normalized for genomic content—reflecting the high origin frequency requirement—PAF15 levels were essentially equivalent across the tested cell types (**Fig. 5C**), underscoring a strict regulatory constraint that prevents the catastrophic consequences of PAF15 surplus. Indeed, analysis of single-cell RNA sequencing data from breast and renal tissues revealed that, although tumor cell types overexpress PAF15, the ratio of PAF15 to PCNA remains similarly low across different cell types (**Fig. S21B-E**). This observation aligns with the distribution of chromatin-bound PCNA and PAF15 in cells from early developmental stages and untransformed cells to cancer cell lines (see **Fig. S8F, G**).

**Figure 5:**
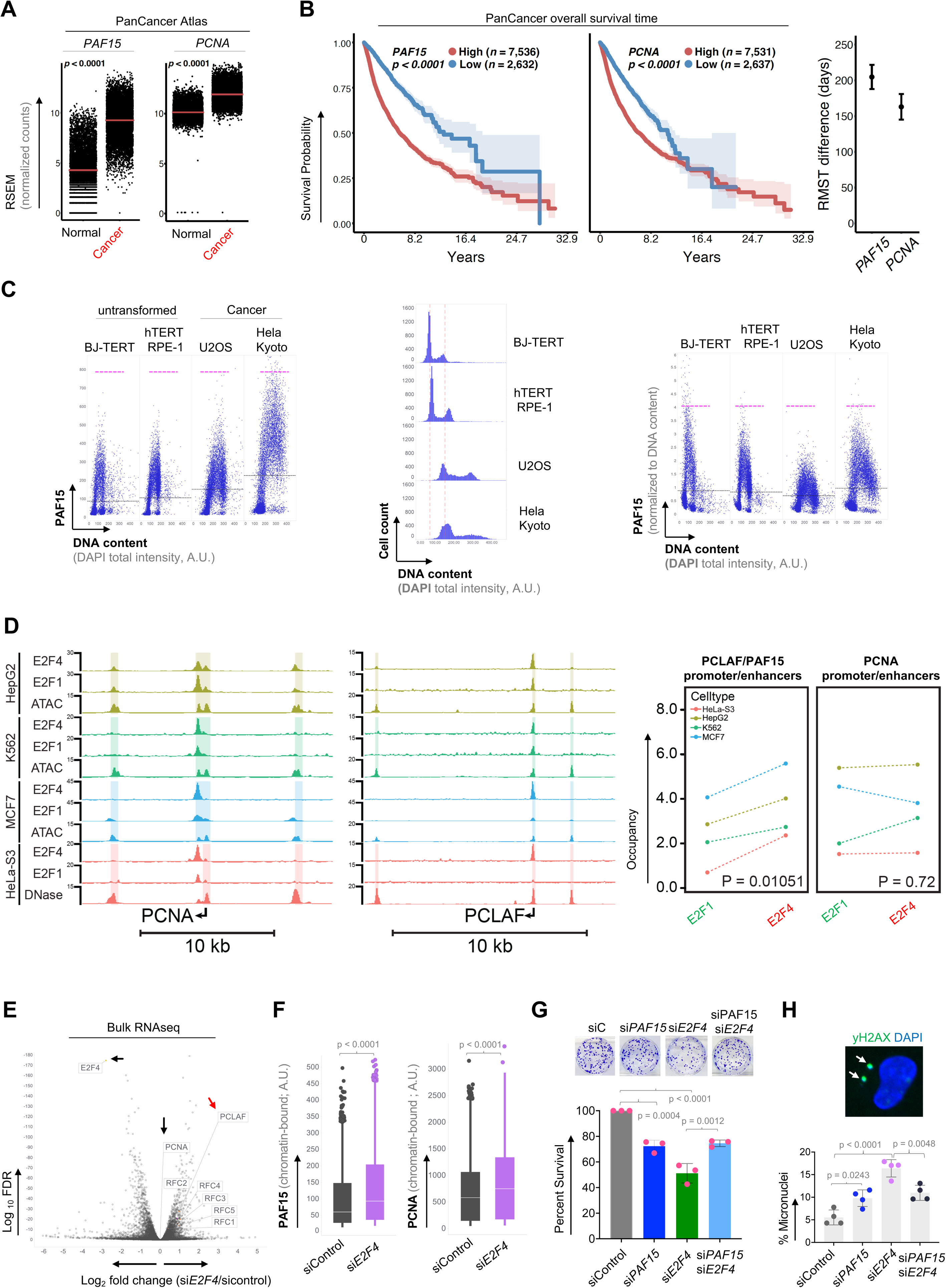
E2F4-mediated repression limits PAF15 dosage and PCNA integrity in genome replication. **(A)** Expression of PAF15 and PCNA in normal (n = 8,167) and tumor tissue (n = 9,471) from 33 cancer types. Data was extracted from the TCGA TARGET GTEx database. Expression is shown as a scatterplot with dots representing patients and red horizontal line showing population means. Mann whitney U test was employed to test difference in population means**. (B)** Left: Kaplan-Meier time-to-event analysis with log-rank p value for PanCancer ATLAS mortality. Patients were stratified by high and low expression of PAF15 and PCNA. Cut-off values were chosen based on the most significant values from Cox-regression on all values from lower to upper quartiles. Right: Difference in restricted mean survival time (RMST) between high and low expression of PAF15 and PCNA. **(C)** QIBC comparison of PAF15 levels in untransformed (BJ-TERT, hTERT RPE-1) and cancer (U2OS, HeLa Kyoto) cell lines before (left) and after (right) normalization to DNA content (middle). A total of 10,000 cells were analyzed per condition. In the scatter QIBC plots, the black horizontal line represents median values, while the dotted pink horizontal line indicates the arbitrarily defined maximum signal for each condition. **(D)** Left: screenshot of Detected peaks of ATAC-seq/DNase-seq and E2F 1/4 ChIP-Seq from indicated cells at the PCNA and PAF15 (PCLAF) locus (data are representative of two independent experiments, irreproducible discovery rate cutoff = 0.05). Highlighted regions indicate enhancers predicted to be causal using the Activity by Contact (ABC) model. Right: average occupancy of E2F4 or E2F1 at PCNA and PAF15 (PCLAF) enhancers derived from the ABC model. **(E)** Volcano plot depicting differentially expressed genes in cells treated with control or E2F4 siRNAs. Y-axis denotes − log10 P values while X-axis shows log2 fold change values showing enrichment of PAF15 with no change of PCNA expression upon E2F4 depletion. **(F)** QIBC analysis of chromatin-bound levels of PAF15 and PCNA under E2F4 depletion. A total of 5,000 cells were analyzed per condition. P values were determined by one-way ANOVA with Tukey’s test. Relative plating efficiency **(G)** and frequency of micronucleation (500 nuclei per condition) **(H)** in U2OS cells treated with indicated siRNAs. Values denote mean ± s.d. P values were determined by one-way ANOVA with Tukey’s test.

Mechanistically, inhibition of global protein degradation using two proteasome-specific inhibitors indicated that under unperturbed conditions, neither the APC/C^CDH1^—which targets PAF15 at the end of the cell cycle(*25*)—nor PAF15Ub2-mediated degradation following DNA damage (*26*) is fully responsible for enforcing PAF15 dosage limitation during S-phase (**Fig. S22A**).

Consequently, we examined whether the transcriptional program controlling PAF15 expression could be responsible. Indeed, transcription inhibition and blocking canonical E2F transcription factors by inhibiting CDK4/6, which prevents RB phosphorylation and maintains RB in an active, E2F-repressive state(*48*), resulted in markedly reduced PAF15 levels (**Fig. S22B-D**). Thus, we tested whether context-specific binding of E2F to regulatory elements—via activators (E2F1-3) and repressors (E2F4-5)(*49, 50*)—regulates PAF15 dosage to maintain a low PAF15-PCNA ratio as seen at the transcript number level (**Fig. S22D, E**). To dissect this, using the Activity-by-Contact (ABC) model, which integrates enhancer activity (via chromatin accessibility) with enhancer-promoter contact frequency(*51*), we identified the causal enhancers for PAF15 and PCNA (**Fig. 5D**). To further validate a causal association between these enhancers and PAF15 expression, we investigated if the enhancers harbor expression quantitative trait loci (eQTLs) for PAF15, those are naturally occurring single nucleotide polymorphisms (SNPs) at which the genotype is associated to PAF15 expression. We found that all three predicted causal enhancers harbor such loci (**Fig. S22F**), thereby further validating that these enhancers regulate PAF15 expression. Utilizing ENCODE-based chromatin immunoprecipitation sequencing (ChIP-seq) data, occupancy of E2F1 versus E2F4 revealed significant enrichment of repressive E2F4 on the PAF15 enhancer relative to E2F1, while PCNA enhancers exhibited balanced binding (**Fig. 5D**). Indeed, depletion of E2F4 led to a marked increase in PAF15 expression without affecting PCNA, as confirmed by RNA sequencing and QIBC across several cell types (**Fig. 5E; Fig. S23A, B**).

Finally, rise in PAF15 protein, following E2F4 loss, was accompanied by enhanced chromatin loading of PAF15 and improved PCNA stability, yet it triggered cell cycle arrest and compromised both cell survival and genome integrity across cell types—effects that were rescued by subsequent PAF15 loss (**Fig. 5F-H**; **Fig. S23C-E**).

Together, these findings demonstrate that deregulation of PAF15 expression due to disrupted transcriptional control results in a lethal cell cycle and replication response, mirroring the effects observed with PAF15 mislocalization to the leading strands caused by TIMELESS/CLASPIN loss.

## DISCUSSION

Collectively, our findings uncover a previously concealed, dosage-limited DNA strand– specific mechanism that orchestrates genome replication during an unperturbed cell cycle. We identify PAF15 as a critical low-dosage factor that binds PCNA via its high-affinity PIP motif—exclusively on the lagging strand—while it is effectively excluded from the leading strand by TIMELESS-CLASPIN. Whereas most PIP-containing proteins dock onto the outer surface of the PCNA ring, specifically along the interdomain connecting loop(*52*), PAF15 uniquely traverses the PCNA-DNA channel, thereby enhancing PCNA stability and slowing its dynamic exchange at the lagging strand. Loss of PAF15 compromises PCNA chromatin stability and OkF maturation, leading to moderate but persistent DNA replication stress and forcing cells to depend on non-canonical, PARP1-mediated processing for survival. Importantly, PAF15-PCNA interactions on the lagging strand become saturated during stochastic origin firing, so that the activation of new origins following checkpoint loss fails to provide adequate lagging strand processing capacity via canonical PCNA-dependent mechanisms (**Fig. S24A**). The striking convergence of PAF15-PCNA exhaustion with ATR-enforced origin firing restriction uncovers yet another fundamental role of the replication checkpoint. Without this safeguard, replication would proceed with uncoordinated leading and lagging strand synthesis, ultimately triggering genome instability and precipitating cell death or disease progression.

The late evolutionary emergence of PAF15 with its unique PCNA-binding mode in metazoans is intriguing (**Fig. S24B**). On one hand, it offers a finely tuned mechanism for OkF maturation; on the other, its overexpression or lack of regulation (especially without TIMELESS-CLASPIN) risks disrupting leading strand PCNA function or hindering lagging strand processing by competing out OkF proteins. PAF15 may have evolved in higher eukaryotes as a compensatory mechanism to enhance DNA binding by PCNA, thereby supporting efficient DNA replication despite the reduced inherent affinity of metazoan PCNA. Alternatively, this adaptation may necessitate significantly tighter regulation of OkFs, as the dramatically increased replication origins in metazoans amplify the risk of replication gaps and their mutagenic consequences (*16, 53, 54*) in the absence of such stringent control of lagging strand processing.

Since the foundational ‘replicon theory’ established that a replicon—a unit of DNA replication—requires both an initiator and a replicator(*55*), identifying the rate-limiting mechanisms in genome replication has been a focal point for understanding how genomes are faithfully maintained. Several such mechanisms have since been identified, all converging on the S-phase checkpoint(*5, 11, 56, 57*). Our finding that PCNA-lagging strand capacity becomes exhausted concurrently with stochastic origin activation adds an unexpected twist to the well-established fact that PCNA—the driving force behind lagging strand processing—is one of the most abundant replisome factors.

Rate-limiting regulation of lagging strand-specific PCNA by PAF15 illuminates several concealed replisome principles that enable DNA replication adaptation in physiology and pathophysiology (**Fig. S24A**).

First, limiting the pool of PAF15 enables cells to fine-tune PCNA dynamics specifically on the lagging strand, ensuring that even when PAF15 is fully exhausted, replisome progression remains unaffected. Indeed, cells defective in lagging strand mechanisms have been shown to proliferate, albeit with heightened replication stress responses(*20, 58*).

Second, the lagging strand naturally forms single-stranded DNA loops and presents checkpoint-activating substrates, such as 5′ RNA primers(*59*). Thus, under conditions of natural exhaustion of PAF15–PCNA–mediated lagging strand processing, unligated OkFs would efficiently trigger the S-phase checkpoint without necessitating replication fork stalling or DNA damage. These new findings further highlight that the coordinated—whether concerted or stochastically coupled—interaction between the S-phase checkpoint and PAF15-PCNA–mediated strand-specific exhaustion is essential not only for the timely activation of replication origins but also for safeguarding key replisome components, thereby protecting both the replisome and replication fork structure. Indeed, failure to restrain excessive origin firing has also been proposed to trigger excessive remodeling of replication forks by DNA translocases such as SMARCAL and HLTF(*29, 60*), irreversibly converting them into substrates for genome breakage and cell death.

Third, because PAF15 expression is tightly regulated by the dynamics of transcription factors E2F4 and E2F1, cells can adjust their replication machinery in response to varying demands—even as cancer cells expand their genomes and fine-tune the frequency of replication origins.

Finally, maintaining a surplus of PAF15 is more detrimental than exhausting PCNA-mediated lagging strand processing, as excess PAF15 beyond its physiological levels or its unrestrained access to the leading strand disrupts PCNA functions and compromises replisome integrity. Following this reasoning, we propose that augmenting PAF15 via its PCNA-DNA-targeted peptides will disrupt strand-specific PCNA dynamics in cancer cells that display a higher density of replication origins and excessive chromatin-bound PCNA compared to normal cells.

## Supporting information

Supplemental material combined with Sup Figures

## ACKNOWLEDGMENTS

This work was supported by the research funding from the Lundbeck Foundation Fellowship (R345-2020-1770), EU - Horizon Europe - ERC Starting Grant (META-SURVEILLANCE-101077859), The Danish Cancer Society (R325-A18913) to KS. Novo Nordisk Foundation (no. NNF21OC0068929) to JGSM. The Czech Science Foundation Junior Star (grant no. 22-20303M), the European Union’s Horizon 2022 Widera Talent program (ERA grant agreement no. 101090292) and EMBO Installation Grant (grant no. IG-5689-2024) to H.P.-S. The Danish Molecular Biomedical Imaging Center (DaMBIC, University of Southern Denmark) is supported by the Novo Nordisk Foundation (NNF) (NNF18SA0032928). We thank Jiri Lukas, Deo Prakash Pandey, Rasmus Siersbæk, and members of the Somyajit lab for stimulating discussions and critical reading.

