## Supplemental material combined with Sup Figures for "PAF15-PCNA assembly exhaustion governs lagging strand replication and replisome integrity"

**The PDF file includes:**

**Materials and Methods**
**Figs. S1 to S24**
References

### Materials and Methods

#### Cell culture:

The human U2OS osteosarcoma cell line (ATCC, HTB-96), HeLa Kyoto cervical carcinoma cell line (CVCL_1922), primary immortalized retinal epithelial cell line hTERT-RPE1 (ATCC, CRL-4000), primary immortalized foreskin fibroblast BJ cells (ATCC, CRL-2522) and their derivatives were grown in Dulbecco's modified Eagle's medium (DMEM, high glucose, Glutamax) containing 10% FBS and penicillin-streptomycin antibiotics (Thermo Fisher Scientific). E14 mouse embryonic stem cells (mESCs) were cultivated in 'ESC medium' supplemented with LIF (Leukemia inhibitory factor, 1,000 U $ml^{-1}$; Millipore) and differentiation was intiated by LIF removal. ESC medium: Glasgow Minimum Essential Medium (Sigma), 15% FBS (Gibco), 1 mM Sodium Pyruvate (Sigma), 1 × Non-Essential Amino Acids (Sigma), 1 × Penicillin-Streptomycin-L-Glutamine (Life Technologies), 0.1 mM β-Mercaptoethanol (Sigma). All cell lines and their derivatives were cultured under standard cell culture conditions (37 °C with 5% CO2, humidified atmosphere), and were routinely tested for mycoplasma contamination (MycoAlert, Lonza) and always found negative.

#### Chemical reagents:

HU (RNRi; Sigma-Aldrich, H8627; solubilized in $H_2O$), Ceralasertib-AZD6738 (ATRi; Selleckchem, S7693), Adavosertib- MK-1775 (WEE1i; Selleckchem, s1525), Palbociclib- PD-0332991 (CDK4/6i; Selleckchem, S1116), LNT 1 (FEN1i ; Tocris, 6510), MG132 (Proteasome inhibitor; Selleckchem, s2619), Bortezomib (Proteasome inhibitor; Selleckchem, S1013), PDD 00017273 (PARGi; TOCRIS, 5952), Docxycycline (Fisher scientific, BP2653-5), T2AA (PCNAi; TOCRIS, 4723), Triptolid (TOCRIS, 3253), Nocodozole (TOCRIS, 1228), CldU (Sigma-Aldrich, c6891), IdU (Sigma-Aldrich, I7125). The drugs were reconstituted in DMSO and were employed as indicated in figure legends.

#### Generation of knockout and complementation cell lines:

Knockout of the *KIAA0101*/*PAF15* gene in U2OS, Hela Kyoto, hTERT-RPE1 cells was generated using single gRNA(targeting exon 2, GGCTGCTCGAGCCCCCAGAA) cloned into pSpCas9(BB)-2A-Puro (PX459) V2.0 (Addgene Plasmid #62988**,** gift from Feng Zhang) via the BbsI restriction site followed by Lipofectamine LTX Plus transfection. After 2 days, transfected cells were selected in media containing 1 µg/ml puromycin (InvivoGen, ant-pr-1). After 30-40 h transfected cells were recovered in plain media and serially diluted into single-cell per well of 96 well plate to obtain single colonies, expanded, and tested for knockout efficiency by Immunofluorescence of PAF15 using high-content imaging (QIBC)-screen, western blotting and Sanger sequencing of gRNA targeting sites. Only cell lines that passed all validation steps were used. Clones with successful knockout of *PAF15* were selected for phenotypic validation. Similar approach by transfecting either p53 single gRNA (PX459-TP53-exon4, Addgene Plasmid # #217455**,** gift from John Diffley) or in combination with PX459-PAF15-exon2 (this paper) to generater *p53* knockout or *p53* + *PAF15* double knockout in hTERT-RPE1 cells. Knockout of *CTF18* gene in U2OS was generated using single gRNA (targeting exon 2, GGCCTCCTCGAACGTCCGCG) as describe above.

### Constitutive and T-REx inducible cell lines:

For complementation assays with constitutive expression, U2OS-*PAF15*-knockout cell lines were transfected with the following variants plasmids: 1X Myc-1X FLAG tag PAF15wt (Origene #RC200694), 1X FLAG-PAF15 PIP box mutant (F68F69 to AA mutation), 1X FLAG-PAF15 KEN box mutant (K78A mutation). Appropriate DNA constructs were transfected using Lipofectamine LTX Plus reagent in *PAF15*-knockout cells. Transfected cells were serially diluted serially diluted into single-cell per well of 96 well plate to obtain single colonies and selected with DMEM containing geneticin (Gibco, 10131-027) for 12 days. Individual colonies were expanded and tested by immunofloresence, employing quantitative-Imaged-Based-Cytometry (QIBC) for FLAG tag/ Myc-tag PAF15 (cellular localization) and expression level by western blotting, using antibodies against PAF15.

All the inducible cell line were generated utilizing tetracycline-regulated expression of gene of interest, using T-REx™ (Thermo Fisher Scientific, K102002). U2OS derivative (naïve U2OS, U2OS *PAF15*-knockout cell lines) were introduced with doxycycline inducible of each PAF15 wt, PAF15 PIP box mutant, PAF15 KEN box mutant, and ∆PAF15 variants (2-61 aminoacid deletion, 2-11 aminoacid deletion or 2-6 aminoacid deletion) were cloned in pcDNA™4/TO. These individual PAF15 variant constructs in pcDNA™4/TO were co-transfected with pcDNA™6/TR plasmid (expression vector for Tet repressor) by Lipofectamine LTX Plus reagent (Thermo Fisher Scientific, 15338-100). After 2 days, transfected cells were selected in DMEM containing Blasticidine and Zeocine (Thermo Fisher Scientific, Blasticidine: A1113902; Zeocine: R25001). Upon reaching 60-70% confluency, cells were serially diluted into single-cell per well of 96 well plate to obtain single colonies, expanded, and tested for PAF15 induction upon doxycycline by employing QIBC, western blotting and high-resolution microscopy.

### *KIAA0101*/*PAF15*-degron cell line:

A derivative of the U2OS cell line expressing C-terminally endogenously AID–GFP-tagged PAF15 was generated using CRISPR-Cas9 as previously described(*61*). Briefly, guide RNA targeting for C-terminal of PAF15 locus (guide: TAACGTCTCCTTGTTTACCC) was cloned into pX330-U6-Chimeric_BB-CBh-hSpCas9 (Addgene plasmid 42230, a gift from F. Zhang) via the BbsI restriction site. Cells were co-transfected by Lipofectamine LTX with Plus reagent (Thermo Fisher Scientific, 15338-100) with pX330 plasmid containing cloned gRNA, a donor plasmid containing the tag (AID-GFP) with flexible linker flanked by 900 bp homology arms complementary to the C-terminus of the PAF15 and pCMV6-A-Puro-Tir1-9xMyc plasmid conferring puromycin resistance. After 24 hours of transfection, the cells were selected with DMEM containing puromycin (1 μg/ml) for 3 days and then serially diluted onto 100 mm dishes. The cells were grown in DMEM to obtain single colonies, which were expanded for further characterization by junction PCR spanning the C-terminal of PAF15. Selected clones were functionally validated by immunofluorescence (sub-cellular localization), employing QIBC. Only cell lines that passed all validation steps were used.

### PCNA chromobody,  TurboID-PCNA and CDK2 repoter cell lines:

U2OS cells, U2OS *PAF15*-knockout cell line were transfected with plasmid (RFP–pCellCyc- leChromobody) containing RFP–PCNA chromobody (Chromotek, #ccr) encoding a single-chain antibody to endogenous PCNA. Single clones were selected by geneticin as described in the previous section. Using the same procedure, pBABE-NLS-HA-TurboID-PCNA (Addgene plasmid #215074 gift from Michele Pagano) was introduced into naïve U2OS cells. Single clones were selected by puromycin as above. Similarly, CDK2 reporter (Fluorescently tagged segment of DNA Helicase B that translocates from the nucleus to the cytoplasm in response to phosphorylation by CDK2) cell line was generated in naïve U2OS cell by introducing DHB-mVenus (Addgene plasmid # 136461 gift from Tobias Meyer and Sabrina Spencer).

### Gene silencing by siRNA:

Transfections of siRNA duplexes were achieved with Lipofectamine RNAiMAX (Thermo Fisher Scientific) at a final concentration of 1-20 nM for 48 h (please see figure legends for more details). For *TIMELESS*, *CLASPIN* knockdown, transfection was performed with 1-5 nM for 24-30h to prevent adverse cell cycle effects. The siRNAs were purchased from Thermo Fisher Scientific as Silencer Select reagents targeting the following genes. *KIAA0101*/*PAF15* (#A:s18863 #B:s18862), *TIMELESS* (s17054), *CLASPIN* (s34330), UHRF1 #A(s26553), #B(s26555), *FEN1* (s5105), *RAD9* (s11720), *ATAD5* (s36632), *E2F4* (#1:114193, #2:114194,

#3:s4414, #4:s4415). Non-targeting siRNA from Thermo Fisher Scientific (Ambion negative control #1 (4390844)) was used as control siRNA in all experiments.

### Antibodies:

Antibodies to the following proteins were used: KIAA0101/PAF15 (rabbit, Abcam, ab226255, 1:1000 for IF, 1:2000 for WB), KIAA0101/PAF15 (rabbit, Cell Signaling, 81533S 1:1000 for IF; 1:2000 for WB), KIAA0101/PAF15 (mouse, Santa Cruz Biotechnology, sc-390515, 1:1000 for IF), TIMELESS (rabbit, Abcam, ab109512, 1:1000 for IF with preextraction before fixation, 1:1000 for WB), CLASPIN (mouse, Santa Cruz Biotechnology, sc-376773, 1:100 for IF with preextraction before fixation; 1:200 for WB), PCNA (mouse, Santa Cruz Biotechnology, sc-56, 1:2000 for WB), PCNA (human, immunoconcepts, 1:500 for IF preextraction before fixation), RPA2 (mouse, Abcam, ab2175, 1:500 for IF with preextraction before fixation), Phospho RPA2 (S33) (rabbit, Bethyl, A300-246A, 1:1000 for IF), MCM2 (mouse, Novus, H00004171-M01, 1:1000 for WB; 1:1000 for IF with preextraction before fixation), MCM7 (mouse, Santa Cruz Biotechnology, sc- 9966, 1:2000 for WB; 1:500 for IF with preextraction before fixation), CDC45 (rabbit, Cell Signaling, 11881S, 1:1000 for WB), POLE1 (rabbit, Abcam, ab226848, 1:1000 for IF with preextraction before fixation, 1:2000 for WB), POLD1 (rabbit, ab186407, 1:1000 for IF with preextraction before fixation and 1:2000 for WB), histone H2B (rabbit, Abcam, ab1791, 1:2000 for WB), FLAG M2 (Sigma, F1804, 1:1000 for WB, 1:500 for IF), 53BP1 (rabbit, Novusbio, NB100-305, 1:2000 for IF), 53BP1 (mouse, Millipore, Mab3802 1:1000 for IF), α-tubulin (mouse, Santa Cruz Biotechnology, sc5286, 1:500 for WB), CyclinA2 (rabbit, Abcam, ab181591, 1:5000 for IF), GFP (rabbit, Chromotek, PABG1, 1:2000 for IF with preextraction before fixation, 1:5000 for WB), H2A.X Phospho Ser139 (mouse, Biolegend, 613402, 1:1000 for IF), H2A.X Phospho Ser139 (rabbit, Cell Signaling, 14655S, 1:2000 for IF), Poly/Mono-ADP Ribose (rabbit, Cell Signaling, 89190S, 1:1000 for IF with preextraction before fixation), FEN1 (rabbit, Invitrogen, MA5-33136, 1:1000 for IF with preextraction before fixation), LIGASE-1 (rabbit, Abcam, ab177946, 1:1000 for IF with preextraction before fixation), RFC-1 (rabbit, Sigma Atlas, HPA069306, 1:100 for WB), Myc-tag (rabbit, Abcam, ab9106, 1:1000 for IF, 1:2000 for WB), CTF18 (mouse, Santa Cruz Biotechnology, SC374632, 1:1000 for WB), HA-tag (mouse, Santa Cruz Biotechnology, SC7392), phospho CHK1 (S345) (rabbit, Cell Signaling, 2348S, 1:1000 for WB), phospho CHK2 (T68) (rabbit, Cell Signaling, 2197S, 1:1000 for WB). Validation of all commercially available primary antibodies is provided on the manufacturers' websites. Secondary-antibody conjugates for IF included goat-anti mouse and goat-anti rabbit Alexa Fluor 488 (A11029; A11034), Alexa Fluor 568 (A11031; A11036), Alexa

Fluor 647 (A21236; A21245) reagents (highly cross-absorbed) each diluted 1:1000, matched to the appropriate filter set for fluorescence microscopy (Thermo Fisher Scientific).

**DNA fiber analysis:**

DNA fiber speards were performed as described(*14*). Briefly, cells ($10^5$) were pulse labeled with 25 µM CldU for the indicated time (Please see Figure panels for respective labelling protocols), washed three times with DMEM medium, and pulse-labeled with 250 µM IdU with or without indicated treatment for the indicated time. Labeled cells were harvested on ice-cold PBS and mixed with unlabeled cells (1:3). Subsequently, 2 µl of the cell suspension was placed on SuperFrostTM slides (AB00008032E01MNZ20) and mixed with 8 µl of lysis buffer (0.5% SDS, 200 mM Tris pH 7.5, 50 mM EDTA) followed by vigorous pipetting for *in situ* lysis. After 2 minutes of incubation, slides were tilted to allow the lysate to flow along the slide slowly until to the end of slide. Next, slides were fixed in methanol:acetic acid (3:1) for 12-15 min, washed four times in PBS, and transferred to 2.5 M HCl for DNA denaturation for 80 min. Afterwards, slides were neutralized by washing four times in PBS and blocked in blocking buffer (1x PBS, 0.1% TritonX, 1% BSA) for 5 minutes. CldU was stained by incubating slides with rat anti-BrdU antibody (Abcam ab6326, 1:100 in blocking buffer) for 90 min. Afterwards, slides were washed once with PBS containing 0.1% Tween, followed by three wash steps with PBS, fixed with 4 % formaldehyde for 12 min and incubated with AlexaFluor 594–conjugated goat anti–rat IgG (1:100; Thermo Fisher Scientific; A-11077) for 60 min. Slides were washed four times with PBS, and IdU was stained using mouse anti-BrdU antibody (1:100; Becton Dickinson, 347580) overnight at 4 °C followed by AlexaFluor 488–conjugated goat anti–mouse IgG (1:100; Thermo Fisher Scientific; A11029) for 90 min. Fibers were acquired using Olympus BX53 Upright Fluorescence Microscope with a 40x air objective. For quantification of replication structures, at least 200 DNA fibers were counted per experiment. The lengths of red (CldU) or green (IdU) labeled patches were measured using the Fiji ImageJ software (National Institutes of Health). Fork speed in kb/min was calculated by multiplying the measured length in µm with a conversion factor of 2.59 kb/µm and dividing by the duration of the labeling pulse.

**Immunofluorescence:**

Cells were grown on round 12-mm diameter, 1.5-mm-thick glass coverslips (cleaned in 96 % ethanol, dried, and autoclaved). Unless stated chromatin-bound, cells were washed with ice-cold PBS and fixed in 4% buffered formaldehyde for 12 min at room temperature before permeabilization with PBS containing 0.2 % Triton X-100 for 5 min. For assessing chromatin-bound proteins, cells were first pre-extracted with ice-cold PBS containing

0.2 % Triton X-100 for 2 min on ice before fixation in 4% buffered formaldehyde for 10 min at room temperature. When Click-iT EdU staining was performed, cells were incubated for 30 min in 10 µM EdU before fixation or pre-extraction. EdU staining was performed according to the manufacturer's instructions (Thermo Fisher Scientific) prior to incubation with primary antibodies. All antibodies were diluted in Dulbecco's modified Eagle's medium (DMEM, high glucose, Glutamax) containing 10 % FBS. Primary-antibody incubations were performed at room temperature for 1 h. Coverslips were washed three times with PBS containing 0.2 % Tween (Sigma-Aldrich). Secondary-antibody incubations were performed at room temperature for 30 min and were supplemented with 4 ,6-diamidino-2-phenylindole dihydrochloride (DAPI, 0.5 mg/ml; Sigma-Aldrich, D8417) to counter-stain DNA.

For accessing unligated Okazaki fragments (OkFs), the relevant cells (please also see figure legends) were incubated with 10 µM CldU for 48 hours to label the parental template DNA. After washing, the cells were exposed to 10 µM EdU for the last 60 minutes to label nascent DNA—both within the replisome and in postreplication regions—to mark S-phase cells. The cells were then fixed and permeabilized as detailed above and incubated with an alkaline buffer (1X: 50 mM NaOH and 1 mM EDTA in PBS) for 30 minutes before proceeding with EdU Click-iT staining, immunostaining of CldU (using a rat anti-BrdU antibody, Abcam ab6326, at 1:500), and counterstaining of DNA with DAPI as described above.

After three washes in PBS, coverslips were washed twice in distilled water, dried on 3mm paper, and mounted in 4.5 µl Mowiol-based mounting medium (containing Mowiol 488 (Calbiochem)/glycerol/Tris-HCL, pH 8.5). For all the confocal and STED imaging, slides were mounted with Prolong Diamond Antifade Mountant (Thermo Fisher Scientific, #P36961).

### Quantitative image-based microscopy (QIBC):

QIBC was performed as previously described(*14*). Briefly, images were acquired with a ScanR inverted microscope high-content screening station (Olympus) equipped with wide-field optics, a 203, 0.75-NA (UPLSAPO 203) air objective, fast excitation and emission filter-wheel devices for DAPI, FITC, Cy3, and Cy5 wavelengths, an MT20 illumination system, and a digital mono- chrome Hamamatsu ORCA-R2 CCD camera (yielding a spatial resolution of 320 nm per pixel at 203 and binning of 1). Images were acquired in an automated fashion with the ScanR acquisition software (Olympus, 3.4). Depending on cell confluency, 100 images were acquired containing more than 5,000 cells per condition. Acquisition times for the different channels were adjusted for nonsaturated conditions in 12-bit dynamic range, and identical settings were applied to all the samples within one experiment. Images were processed and analyzed with ScanR analysis software.

First, a dynamic background correction was applied to all images. The DAPI signal was then used for the generation of an intensity-threshold-based mask to identify individual nuclei as main objects. This mask was then applied to analyze pixel intensities in different channels for each individual nucleus. After segmentation of nuclei, foci were segmented as above, and the desired parameters for the different nuclei or foci were quantified, with single parameters (mean and total intensities, foci count, and foci intensities) as well as calculated parameters (sum of foci intensity per nucleus). These values were then exported and analyzed with TIBCO Software, version 12.4. This software was used to quantify absolute, median, and average values in cell populations and to generate all color-coded scatter plots. Within one experiment, similar cell numbers were compared for the different conditions (at least 5,000-10,000 cells), and for visualization low x-axis jittering was applied (random displacement of objects along the x axis) to make overlapping markers visible.

### Confocal microscopy:

Confocal images were acquired with a Nikon A1 confocal Ti-2 microscope integrated with a 100x 1.45 NA oil, Plan Apochromat λ, objective and NIS-Elements AR software (version 5.20.02). A resonant scanner equipped with A1-DUG hybrid 4-channel detector was used for image acquisition at 512 x 512 pixels. Laser power, detector gain and exposure time were appropriately adjusted with identical settings applied within a series of experiments. Microscope performance and channel alignment were regularly checked by imaging of 200-nm multicolor fluorescent beads.

### Stimulated Emission Depletion (STED):

For combined stimulated emission depletion (STED) and confocal microscopy, EdU labelled preextracted samples were imaged using an Abberior Facility Line STED microscope (equipped with a 100x NA 1.4 oil objective (UPLSAPO100X Olympus). Click-iT EdU coupled to AlexaFlour 488 was imaged for confocal microscopy but PAF15 (using STAR RED goat-anti-rabbit (1:250, #STRED-1007, Abberior, Göttingen, Germany) was imaged for both confocal and STED microscopy. $5\times 5\ \mu m^2$ regions of interest showing immobilized dye signal were imaged for the indicated number of frames using standard confocal and STED imaging conditions, 10 µs dwell time, 50 nm pixel size and repetition frequency of 40 MHz. To collect emission spectra, 488 and 561 nm laser lines were used with a 10 µW power as measured at the sample plane. All the data were acquired using the iMSPECTOR version 16.3.13787 acquisition software. Mean filter with a radius of one pixel was applied to remove background noise.

### Fluresence recovery after photobleaching (FRAP):

U2OS cells expressing GFP tagged–PCNA were seeded in imaging dishes (Nunc, Lab-Tek, 155361) and before imaging transferred to $CO_2$-independent medium. Fluorescence recovery after photobleaching (FRAP) data were acquired using Nikon A1 Ti2 microscope with 60x 1.2 NA Plan Apo Water Immersion objective and NIS-Elements (Ver. 5.30.02) software, under stable temperature conditions of 37 °C. After 10 pre-bleaching frames (pre), a single bleach pulse (488-nm argon laser set to 100% power) was delivered in a defined region followed by time-lapse imaging for 3 minutes at maximum scanning speed (6 frames per second) with the laser transmission attenuated to 2.5%. Image analysis was performed by first extracting the mean GFP-associated fluorescence intensity for each time point in the following regions: Bleaching region (Ifrap(t)), background outside the nucleus (Iback(t)), signal within the nucleus in which bleaching was performed (Iref(t)). After background correction, double normalization was applied, which corrects for differences in the starting intensity in the Ifrap region and for loss in total nuclear fluorescence in the Iref region owing to the bleaching pulse and to acquisition bleaching.

### Cell synchronization:

U2OS cells were synchronized at the G2/M phase by the addition of nocodazole. Exponentially growing U2OS cells were incubated with 200 ng/ml nocodazole for 20 h. For enrichment of cells into different cell cycle phase, cells were washed and cultured in fresh DMEM media. The cells were collected at 0, 5, 8, 10, 12, 15 and 24 h and fixed with 4% buffered formaldehyde for 10 min at room temperature and permeabilized with PBS containing 0.2 % Triton X-100 for 5 min. QIBC based PAF15 total pool and cell cycle analysis was performed with Click-iT EdU staining, assessing sequentiall cell cycle phase transition. Complementarily, western blot was performed to assess the PAF15 in soluble (S) and chromatin-bound (CB) subcellular fractions.

### RNA isolation for RNA sequencing and quantitative real-time PCR:

$2 \times 10^6$ U2OS, Hela Kyoto, hTERT-RPE1, BJ, cells were subjected to either no treatment or treatment with 10 µM CDK4/6 inhibitor for 24 h. Total RNA was isolated using the RNeasy® Mini Kit (Qiagen), following the manufacturer’s protocol. RNA concentration was measured using a NanoDrop spectrophotometer. cDNA was prepared, according to the manufacturer’s instructions, using the AMPIGENE® cDNA Synthesis Kit (Enzo Life Sciences). The qPCR reactions were perfomed in triplicates using iTaq™ Universal SYBR® Green Supermix (Bio-Rad). Relative expression levels were calculated using the $2^{-\Delta\Delta CT}$ method.

The following primer pairs were used: PAF15 Forward (5'-GGCGGGATAGTTTCGGGTC-3') and Reverse (5'-CGAGCAGCCACCACTTTTCT-3'); PCNA_Vr2 Forward (5' GCAGATGTACCCCTTGTTGT-3') and PCNA_Vr2 Reverse (5'- ATCCTCGATCTTGGGAGCCA-3'); GAPDH Forwards (5'-CACCATCTTCCAGGAGCGAG-3') and Reverse (5'- TGATGACCCTTTTGGCTCCC-3').

For DMSO and CDK4/6 inhibitor three biological repeats in hTERT-RPE1 cells, RNA-seq was performed at BGI Genomics using DNBSEQ stranded mRNA libraries generated on the DNBseq™ NGS platform after quality control. The quality of the raw sequencing data was assessed using FastQC (v0.11.9) and MultiQC (v1.10.1). The raw bulk RNA-seq data were aligned to the human genome assembly (GCF_000001405.39_GRCh38.p13), and differential gene expression was analyzed using the DEF analysis plan and a Poisson distribution model at BGI Genomics.
For control and E2F4 depleted three biological repeats in hTERT-RPE1 cells, RNA-seq was performed according to the manufacturer's instructions (TruSeq2, Illumina) using 500 ng of RNA for preparation of cDNA libraries. Sequencing reads were mapped to the human genome (hg38) using STAR55, and tag counts were summarized at the gene level using HOMER56, allowing only one read per position per length. TiCoNE25 was used to cluster differentially expressed genes as determined by DESeq257.

### Clonogenic survival assay:

Naïve cells, gene knockout cells and cells stably expressing PAF15 variant constructs were transfected with control and other siRNAs for 48 h; these cells were  seeded in six well plates in triplicates (200/ 500/ 1000 cells per well). After 24 h, genotoxic treatments were proceeded as indicated in figure legeds. Cells were incubated for 10 days and fixed with 4% formaldehyde and stained with crystal violet. Individual colonies were counted manually and the percentage survival was calculated as values for indicated siRNAs divided by values for control siRNA, after correcting for the respective plating efficiency.

### Proximity Ligation Assay:

PLA was performed as described previously (*14*) with modifications. Briefly, cells were fixed either with methanol (for PCNA interactions) for 15 minutes or 4 % buffered formaldehyde (PAF15 interactions) for 12 minutes and permiabilised with 0.2 % triton X-100 in PBS for 5 minutes. Cells were then blocked for 1 h in DMEM media containing 10 % FBS and  incubated with primary antibody in humidity chamber for 1 hr and secondary antibody probes in humidity chamber at 37 °C for 1 h. *In situ* proximity polymerization followed by ligation was performed using a Duolink Detection Kit (Sigma-Aldrich) and nucleus was counter stain with DAPI. Nuclear foci were imaged using a ScanR inverted microscope and processed for QIBC. At least 5000 cells per condition were analyzed in each experiment.

### Chromatin fractionation:

3x $10^6$ cells were harvested from experimental conditions as mentioned in figure legends. The cells were washed with PBS and harvested by scraping. The soluble protein fraction was removed by incubation in 0.5% Triton X-100 in PBS supplemented with 1x protease and phosphatase inhibitors (PPi) cocktail (Roche). The fractions were centrifuged for 5 min at 4 °C 16000 g. The samples were washed with PBS containing 0.5x PPi. Finally, the cell  pellets were lysed in RIPA buffer (50 mM Tris-HCl, pH 7.5, 150 mM NaCl, 0.1% SDS, 1% Triton X-100, 0.5% deoxycholate (Sigma-Aldrich, R0278-500ML), with benzonase (Merk, E1014) and 100 µg/ml RNaseA (Thermo Fisher Scientific, EN0531). The samples was sonicated at low amplitude, on ice for 2 repeats of 20 s pulses following incubation for 1 h on ice. Chromatin bound protein pools were collected by centrifugation at 4 °C, 16000 g for 30 min. Chromatin bound protein pool concentration was quantified using Pierce™ BCA Protein Assay kit (Thermo Fisher Scientific, 23227). 20-50 µg protein from chromatin fraction was used for western blots.

### Western blotting:

Whole-cell extracts (WCE) were obtained by lysis in RIPA buffer (50 mM Tris-HCL, pH 8.0, 150 mM NaCl, 1.0 % IGEPAL CA-630, 0.1% SDS, and 0.1% Na-deoxycholic acid), supplemented with protease and phosphatase inhibitors (ROCHE) containing benzonase (Novagen). Protein extracts from WCE or chromatin fractions were separated by SDS–PAGE after boiling samples in reducing buffer (DTT and beta-mercaptoethanol) as per standard procedures. Separated proteins were transferred from the gel to a nitrocellulose membrane. The membrane was blocked for 1 h in TBS- 0.1 % Tween containing 5% powdered milk (TBS-T) and subsequently incubated with primary antibodies for 2 hours at room temperature or overnight at 4 °C in TBS-T. Phospho-specific primary antibodies were diluted in 3 % BSA TBS-Tween solution. Secondary peroxidase-coupled antibodies (Vector labs) were incubated at room temperature for 1 hour. ECL-based chemi-luminescence was detected with an Amersham Imager 680 system (Software version 2.0).

### Immunoprecipitation:

$2 \times 10^7$ cells from naïve U2OS, stably expressing U2OS, GFP-PCNA U2OS cells with indicated siRNAs (indicated in figure legends) were harvested and chromatin bound proteins were either processed without crosslinking or cross-linked by incubating cells in 0.1% formaldehyde for 15 min at room temperature. The reaction was quenched by the incubation with 0.125 M glycine. Cells were then collected by scraping and washed with PBS and incubated on ice for 15 min with 0.5% Triton X-100 in PBS supplemented with protease and phosphatase inhibitors (PPi) cocktail (Roche). The samples were washed with PBS containing 0.5x PPi and the nuclear pellets were resuspended in RIPA lysis buffer (50 mM Tris-HCl [pH 7.5], 150 mM NaCl, 0.1% SDS, 1% Triton X-100, 0.5% deoxycholate), with benzonase (Merk, E1014) and 100 µg/ml RNaseA (Thermo Fisher Scientific, EN0531). After incubation of 1 h on ice, samples were sonicated at low amplitude on ice for 2 repeats of 20 s pulses. After centrifugation at 16000 g for 30 min, the supernatant was collected and proteins were quantified using Pierce™ BCA Protein Assay kit. 1000 µg of the chromatin was incubated overnight at 4 °C with anti-Myc magnetic beads (Sigma), or Anti-FLAG magnetic agarose (Thermo Fisher Scientific) or GFP-trap magnetic beads (Chromotek). Similar reactions were also performed for naïve U2OS cells with an equivalent amount of beads. 5-10% of the chromatin was used as input control. Bound proteins were eluted from beads by boiling in NuPAGE LDS Sample Buffer (Thermo Fisher Scientific, NP0007) with NuPAGE™ Sample reducing Agent. Total immunoprecipitates were then analyzed with immunoblotting or processed for mass spectrometry analysis as specified in figure legends.

**Chromatin-bound PAF15 immunoprecipitation-Mass spectrometry:**

Chromatin-bound PAF15 immunoprecipitation proteins products were resolved on a NuPAGE Novex® Bis-Tris 4-12% gel (Invitrogen). Lanes for each sample were sliced (~1 $mm^3$) and gel slices were destained further with a buffer containing 50 mM ammonium bicarbonate and 50% acetonitrile. Gel pieces were dehydrated by the addition of 100% acetonitrile. The gel pieces were incubated with 10 mM DTT for 30-45 min and further incubated for 30 min with 55 mM IAA and subsequently dehydrated with 100% acetonitrile. The proteins were digested with trypsin (Sigma) at 37 °C for 16 h after which the samples were acidified with 0.5 % trifluoroacetic acid (TFA). The resulting peptides were desalted on reversed phase C18 StageTips columns, eluted with 40 µl of 50% acetonitrile, 0.1% TFA followed by 10 µl of 70% acetonitrile, 0.1% TFA and subsequently dried by vaccum centrifugation. Samples were redissolved in 0.1% formic acid (FA) of which 5 of 12µl were loaded on an Easy nLC (Thermo Scientific) equipped with a custom made 2 column setup (precolumn; 100 µm ID, 3.5 cm, Reposil-Pur 120 C18-AQ, 5 µm (Dr. Maisch) and analytical column; 75 µm, 18cm, Reposil-Pure 120 C18-AQ, 3 µm (Dr. Maisch)). Peptides were eluted with a gradient of solvent B (95 % acetonitrile, 0.1 % FA) as specified below, with solvent A being 0.1% FA, and sprayed directly into an Exploris 480 (Thermo Scientific) mass spectrometer; . The gradient was constructed as follows; 5% to 25% B in 70 min , 25% to 40% B in 19 min and 40% to 95% B in 1 min.

On the Exploris 480, MS1 data was obtained at resolution 120K, scan range 350-1600, AGC target 2.5e10 and the Max IT set to auto. MS2 data was recorded as top 12 at a resolution of 30K, AGC target 2e5, Max IT 100 ms and 20 sec dynamic exclusion. Data was searched against the SwissProt database of human proteins using Proteome Discoverer 2.5 (Thermo Scientific) with Mascot 2.7 as the search engine. The m/z tolerance was set to 5ppm for precursors and 0.05 Da for fragment ions.Quantification was done as labelfree quantification based on area under the curve of the (up to) top 5 most abundant peptides for each protein, also utilizing match between runs.

### Proximity labeling TurboID PCNA- Mass spectrometry:

Stably expressing PCNA-TurboID U2OS cells treated with control and PAF15 siRNA were grown to 80 % confluency and proximity labelling was carried out with 50 µM biotin for 30 min. Harvested cells were lysed in RIPA buffer (50 mM Tris, 150 mM NaCl, 0.1% (w/v) SDS, 0.5% (w/v) sodium deoxycholate, 1% (v/v) Triton X-10, pH 7.5), sonicated, cleared by centrifugation, and incubated with streptavidin magnetic beads (Pierce) in RIPA-buffer for enrichment of biotinylated proteins. The beads were sequentially washed with RIPA buffer, Tris buffer (50 mM Tris, 150 mM NaCl, pH 7.5), 1 M KCl, 0.1 M Na2CO3, and 2 M urea in 10 mM Tris pH 7.5 using a KingFisher system (ThermoFisher Scientific).

After enrichment of proximity labeled proteins or chromatin, proteins were precipitated on streptavidin magnetic beads (Pierce) or HILIC microparticles (Resyn BioSciences), respectively, in 70% acetonitrile, reduced, alkylated, and washed sequentially in 95% acetonitrile and 70% ethanol to remove detergents (*62*) and digested on beads with trypsin and Lys-C. The resulting peptides were desalted on StageTips and subjected to LCMS analysis on an EASY nano-LC 1200 system (80-120 min gradients) coupled to an Orbitrap Exploris 480 mass spectrometer (ThermoFisher Scientific). MS data were acquired by data-dependent acquisition and subjected to MaxQuant database searches (*63*) using a human Uniprot reviewed sequence database, ‘match between runs’, and ‘label-free quantitation’.

### AlphaFold-based modeling of protein structures:

AlphaFold 3 with its implementation in the AlphaFold Server was used to predict the DNA - protein structures(*31*). PyMOL was used for structure modelling and generation of high-resolution images. The protein sequences were extracted from the Uniprot database. The input DNA sequence was extracted from the experimentally determined structure of human PCNA with DNA (PDB_ID: 7QO1)(*64*). All the five AlphaFold predicted models were aligned or superimposed using the PyMOL software and only models that showed consistency throughout the models are present in the manuscript. PCNA or its yeast homologue POL30 were used a reference for PyMOL alignment or superimposition and only the models showing a root mean standard deviation (RMSD) lower than 1.0 are used throughout the manuscript. The electrostatic surface visualization of PCNA-DNA-PAF15 was performed with a plugin instalment of APBS in PyMOL (*32*). The interpretation of predicted alignment error plots generated by AlphaFold 3 was performed with the PAE viewer(*65*). The PAE plots were further exported as .svg and .png files and only the later were annotated in Adobe Illustrator to obtain the final figures.

**ABC modelling and analysis of E2F occupancy on PCNA and PAF15/PCLAF causal enhancers:**

We downloaded IDR-thresholded peak lists from ENCODE (*66*) and generated a consensus peak list using iterative overlap peak mergin(*67*). We counted reads in peaks using UCSC tools, and Z-transformed the counts. To predict causal enhancers in cell type at the PCNA and PCLAF/PAF15 loci, we used the ABC model (*51*) using the Z-transformed chromatin accessibility as a proxy for enhancer activity and a power transformation of the distance between the enhancer and transcription start site as proxy for contact frequency as suggested in the ABC paper. We selected enhancers with an ABC score above 0.2 as putative causal enhancers. At these enhancers, we calculated the average occupancy of E2F1 and E2F4 and tested for significant differences between the average occupancy across cell types using a paired T test. List of ENCODE DCC Experiment accession numbers: ChIP-seq: E2F1 in K562 (ENCSR153DWR), ChIP-seq: E2F4 in K562 (ENCSR368GJN), ATAC-seq: K562 (ENCSR868FGK), ChIP-seq: E2F1 in HepG2 (ENCSR717ZZW), ChIP-seq: E2F4 in HepG2 (ENCSR924LSO), ATAC-seq: HepG2 (ENCSR291GJU), ChIP-seq: E2F1 in MCF7 (ENCSR000EWX), ChIP-seq: E2F4 in MCF7 (ENCSR505NMN), ATAC-seq: MCF7 (ENCSR422SUG), ChIP-seq: E2F1 in HeLa-S3 (ENCSR000EVJ), ChIP-seq: E2F4 in HeLa-S3 (ENCSR000EVL), DNase-seq: HeLa-S3 (ENCSR959ZXU)

**Public single-cell RNA sequencing data analysis**:

For investigation of single-cell PAF15 expression in human breast and kidney cancer, features, barcodes, and raw read counts from two publicly available datasets were retrieved from the Gene Expression Omnibus repository GSE176078(*68*) and Human Cell Atlas project Haniffa-Human-10x3pv2(*69*). Both datasets were pre-processed using the pipeline suggested by Seurat (v.4.0.3)(*70*). Low quality cells with <200 detected features and mitochondrial gene contributions >20% were removed from the kidney single-cell data. Subsequently, normalization, scaling, and identification of variable features was employed on both datasets using SCTransform(*71*) regressing out mitochondrial-, ribosomal- and hemoglobin gene percentage. Doublets were estimated and removed using scDblFinder (v1.2.0)(*72*). Batch integration was performed with Harmony (v1.2.0)(*73*). Dimensional reduction by UMAP was recalculated using the integrated lower dimensional space using the first 50 PCs and nearest neighbours set to 15. The original Louvain algorithm was employed for community detection. Automated cell annotation of clusters was carried out with CellTypist (v1.6.3)(*74*) using the Adult_Human_Kidney and Cells_Adult_Breast models for kidney and breast tissue cells respectively.

### TCGA data analysis:

For survival analysis, raw read counts were retrieved from all open-access TCGA projects under the data category transcriptome profiling using TCGAbiolinks (v2.32.0)(*75*). Raw counts were batch-corrected using ComBatseq from the sva package (v3.52.0)(*76*). Patients were split based on the best-performing cutoff in gene expression level between the lower and upper quartile using Cox proportional hazards regression modelling. An estimate of a survival curve was computed using Kaplan–Meier with the survival package (v3.7.0). For gene expression analysis in normal and cancer tissue, RSEM-normalised expression data from the TCGA Target GTEx cohort (n = 19,109) was retrieved from the UCSC Xena platform(*77*). All statistics on TCGA-derived data was performed in R and data visualization was carried out with ggplot2 (v3.5.1)(*78*).

### Reproducibility and Data availability statement:

For all QIBC, Proteomics and RNA-seq experiments minimum of three technical repeats and minimum of two biological replicates were performed. Experiments were not randomized and no blinding was used during data analysis. Sample size, statistical tests and number of replicates for all image-based experiments are specified in the figure legends. All the source data, including numerical and statistical source data and codes will be deposited appropriately at the Dryad (https://datadryad.org/about#our-members) or European Bioinformatics Institute (EBI) BioStudies database (https://www.ebi.ac.uk/biostudies/) and will be immediately accessible as an open resource upon acceptance of this manuscript. All proteomics related data will be deposited to the PRIDE database (https://www.ebi.ac.uk/pride/). Any additional data or information in support of this study will be available from corresponding authors upon reasonable request.

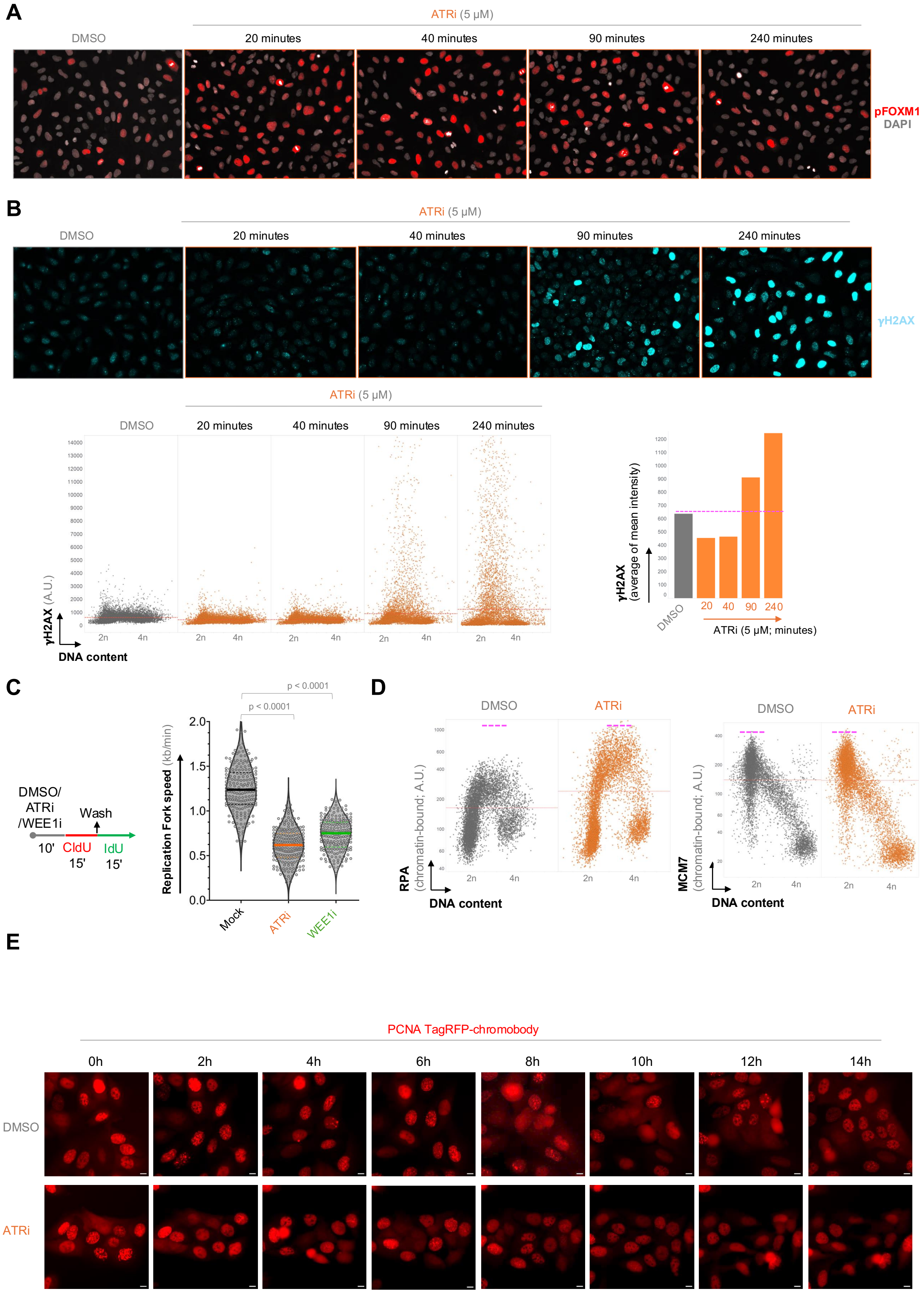
A
ATRi (5 µM)
DMSO
20 minutes
40 minutes
90 minutes
240 minutes
pFOXM1
DAPI
B
ATRi (5 µM)
DMSO
20 minutes
40 minutes
90 minutes
240 minutes
γH2AX
γH2AX (A.U.)
DNA content
2n
4n
γH2AX (average of mean intensity)
DMSO
20 40 90 240
ATRi (5 µM; minutes)
C
DMSO/ ATRi /WEE1i
Wash
10'
CldU 15'
IdU 15'
Replication Fork speed (kb/min)
p < 0.0001
p < 0.0001
Mock
ATRi
WEE1i
D
DMSO
ATRi
RPA (chromatin-bound; A.U.)
MCM7 (chromatin-bound; A.U.)
DNA content
E
PCNA TagRFP-chromobody
0h
2h
4h
6h
8h
10h
12h
14h
DMSO
ATRi

Figure S1

**Fig. S1. Rapid upregulation of CDK1/2 activities and origin firing upon ATR inhibition. (A)** Representative images from high-content microscopy of U2OS cells immunostained for phosphorylated FOXM1 (pFOXM1) following ATR inhibition at the indicated timepoints. **(B)** Top: Representative high-content microscopy images of U2OS cells immunostained for γH2AX following ATR inhibition at the indicated timepoints. Bottom Left: QIBC analysis of γH2AX levels in the cells shown in the top panel. Nuclear DNA was counterstained with 4′,6-diamidino-2-phenylindole (DAPI) (with 2n corresponding to G1 phase and 4n to G2 phase; >8,000 cells per condition). Bottom Right: Quantification of the average mean intensity of γH2AX in cells exposed to the specified timepoints of ATR inhibition. All values are presented in arbitrary units (A.U.). **(C)** (Left) DNA fiber labeling protocol. (Right) Replication speed of forks after mock DMSO or treatments with ATR inhibitor (10 μM) and WEE1 inhibitor Adavosertib (10 μM). n = 200 fibers for each condition. P values were determined by one-way ANOVA with Tukey's test. **(D)** QIBC of cells immunostained RPA2 (Left) and MCM7 (Right) for their chromatin binding. Nuclear DNA was counterstained DAPI (2n corresponds to G1 phase and 4n to G2 phase; >5,000 cells per condition). The horizontal line in QIBC plot depicts the average values. Pink dotted lines indicate the approximate maximum levels of each protein. **(E)** Representative time-lapse snapshots of U2OS cells stably expressing the TagRFP-PCNA chromobody following the indicated treatments and at specified

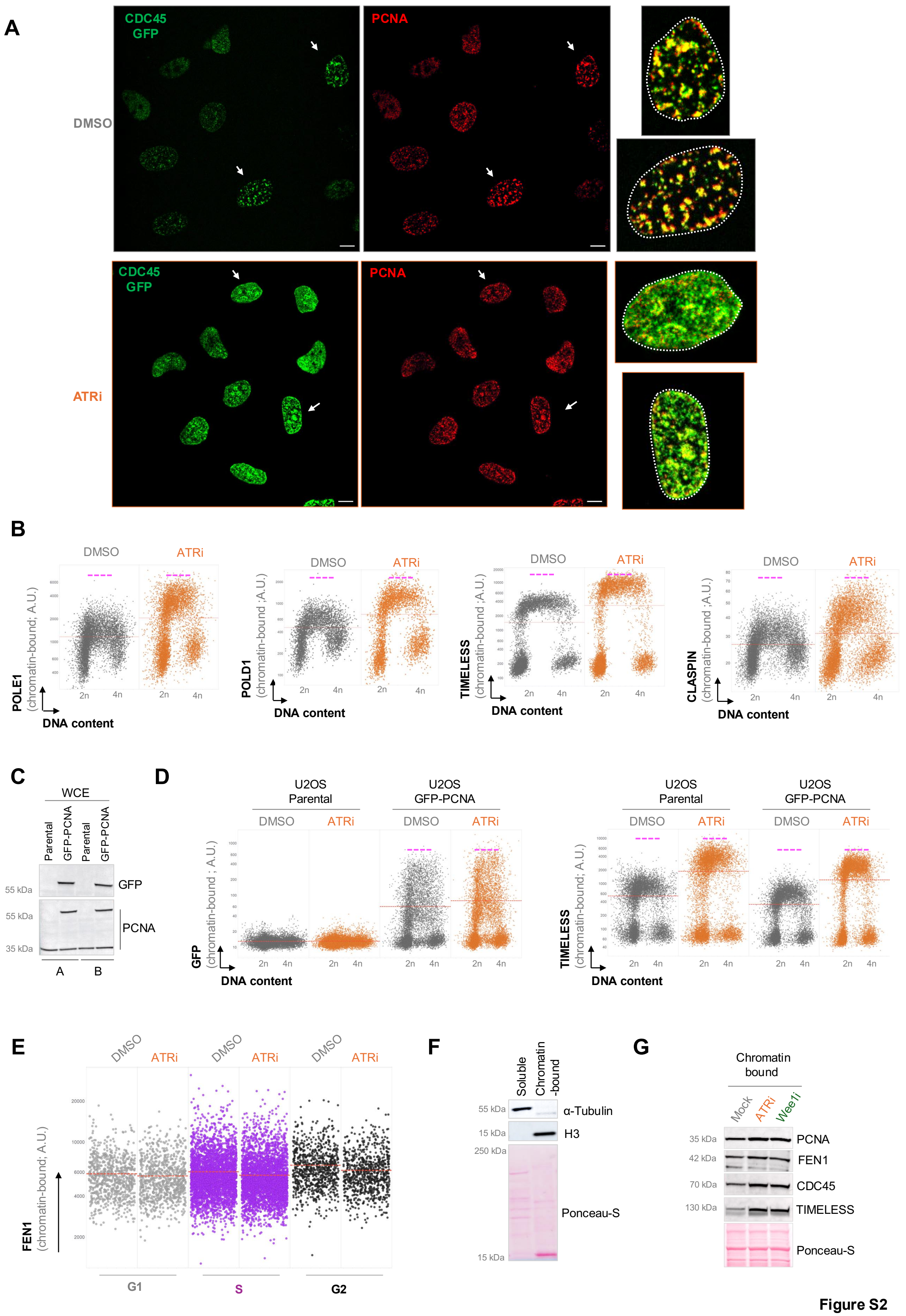
A
CDC45 GFP
PCNA
DMSO
ATRi
B
DMSO
ATRi
POLE1 (chromatin-bound; A.U.)
POLD1 (chromatin-bound ;A.U.)
TIMELESS (chromatin-bound ;A.U.)
CLASPIN (chromatin-bound ;A.U.)
2n
4n
DNA content
C
WCE
Parental
GFP-PCNA
55 kDa
35 kDa
GFP
PCNA
A
B
D
U2OS Parental
U2OS GFP-PCNA
GFP (chromatin-bound ; A.U.)
TIMELESS (chromatin-bound; A.U.)
E
FEN1 (chromatin-bound; A.U.)
G1
S
G2
F
Soluble
Chromatin -bound
15 kDa
250 kDa
α-Tubulin
H3
Ponceau-S
G
Chromatin bound
Mock
Wee1i
42 kDa
70 kDa
130 kDa
FEN1
CDC45
TIMELESS

Figure S2

**Fig. S2 Rapid origin firing upon ATR inhibition and depletion of chromatin-associated PCNA-lagging strand factors. (A)** 3D confocal microscopy image of U2OS cells expressing endogenously tagged CDC45-GFP and immunostained for PCNA following exposure to an ATR inhibitor as described in Fig. 1B. Scale bar, 10 µm. **(B)** QIBC analysis of chromatin-bound POLE1, POLD1, TIMELESS, and CLASPIN in cells treated with ATRi. Nuclear DNA was counterstained with DAPI, where 2n corresponds to the G1 phase and 4n to the G2 phase (with over 5,000 cells analyzed per condition). The horizontal line in QIBC plot depicts the average values. Pink dotted lines indicate the approximate maximum levels of each protein. **(C)** Assessment of GFP tagged PCNA in naïve U2OS and U2OS stably expressing ectopic GFP PCNA. WCE: whole cell extract. **(D)** QIBC analysis of chromatin-bound GFP tagged PCNA (left) and TIMELESS (right) in indicated cells after ATR inhibition. The horizontal line in QIBC plot depicts the average values from over 5,000 cells analyzed per condition. Pink dotted lines indicate the approximate maximum levels of each protein. **(E)** QIBC analysis of chromatin-bound FEN1 across different cell cycle phases upon ATR inhibition. Cell cycle gating was performed using the cell cycle–specific chromatin-binding profile of PCNA. The horizontal line in QIBC plot depicts the average values. **(F)** Subcellular fractionation of U2OS cells followed by immunoblotting for H3 (chromatin-bound fraction) and α-tubulin (soluble fraction). **(G)** Western blot analysis of indicated protein from purified chromatin fractions upon exposure to ATR and WEE inhibitors.

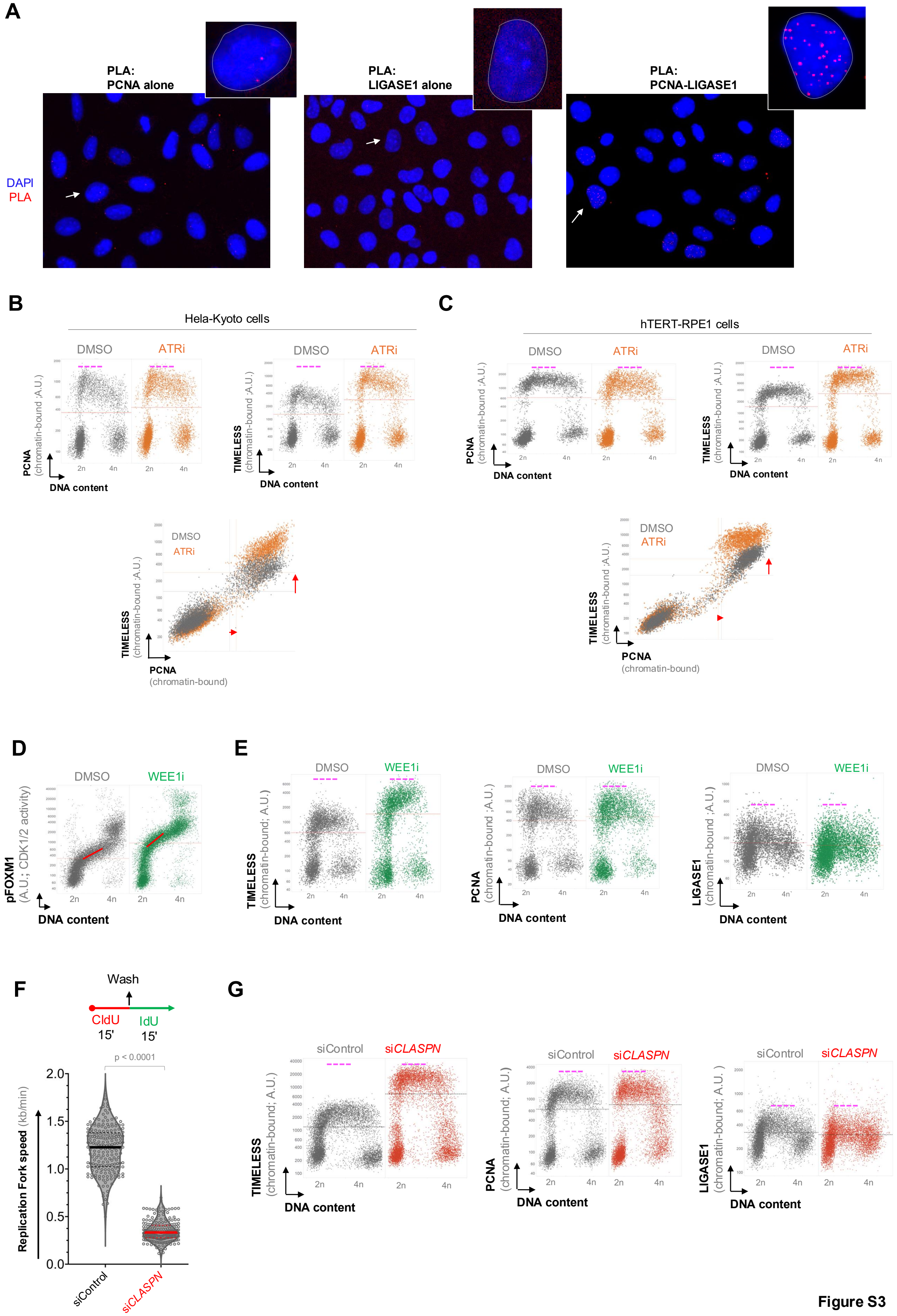
A
PLA:
PCNA alone
PLA:
LIGASE1 alone
PLA:
PCNA-LIGASE1
DAPI
PLA
B
Hela-Kyoto cells
DMSO
ATRi
PCNA
(chromatin-bound ;A.U.)
TIMELESS
DNA content
C
hTERT-RPE1 cells
PCNA
(chromatin-bound)
D
WEE1i
pFOXM1
(A.U.; CDK1/2 activity)
E
LIGASE1
F
Wash
CldU
15'
IdU
15'
p < 0.0001
Replication Fork speed (kb/min)
siControl
siCLASPN
G
(chromatin-bound; A.U.)



Figure S3

**Fig. S3. Natural depletion of chromatin-associated PCNA-lagging strand factors is consistent across cell types and mode of CDK1/2 upregulation. (A)** Representative images of proximity ligation assay (PLA) of PCNA and LIGASE1 on U2OS cells. Arrows indicate cells displayed in more detail. QIBC analysis of chromatin-bound PCNA and TIMELESS in HeLa Kyoto **(B)** and hTERT-RPE1 **(C)** cell lines after ATR inhibition. 2n corresponding to G1 phase and 4n to G2 phase; >5,000 cells per condition. **(D)** QIBC analysis of pFOXM1 in DMSO and WEE1 inhibitor (Adavosertib, 10 µM) treated U2OS cells. n >10,000 cells per condition. (E) QIBC analysis of chromatin-bound fraction of indicated proteins in U2OS cells treated with WEE1 inhibitor. n >5,000 cells per condition (F) (Top) DNA fiber leabeling protocol. (Bottom) Replication fork speed of U2OS cells treated with siCLASPIN. n = 200 fibers for each condition. P values were determined by one-way ANOVA with Tukey's test. **(G)** QIBC analysis of chromatin-bound fraction of indicated protein in U2OS cells treated with siCLASPIN. 2n corresponding to G1 phase and 4n to G2 phase; >5,000 cells per condition. The horizontal lines in QIBC plot depicts the average values. Pink dotted lines indicate the approximate maximum levels of each protein.

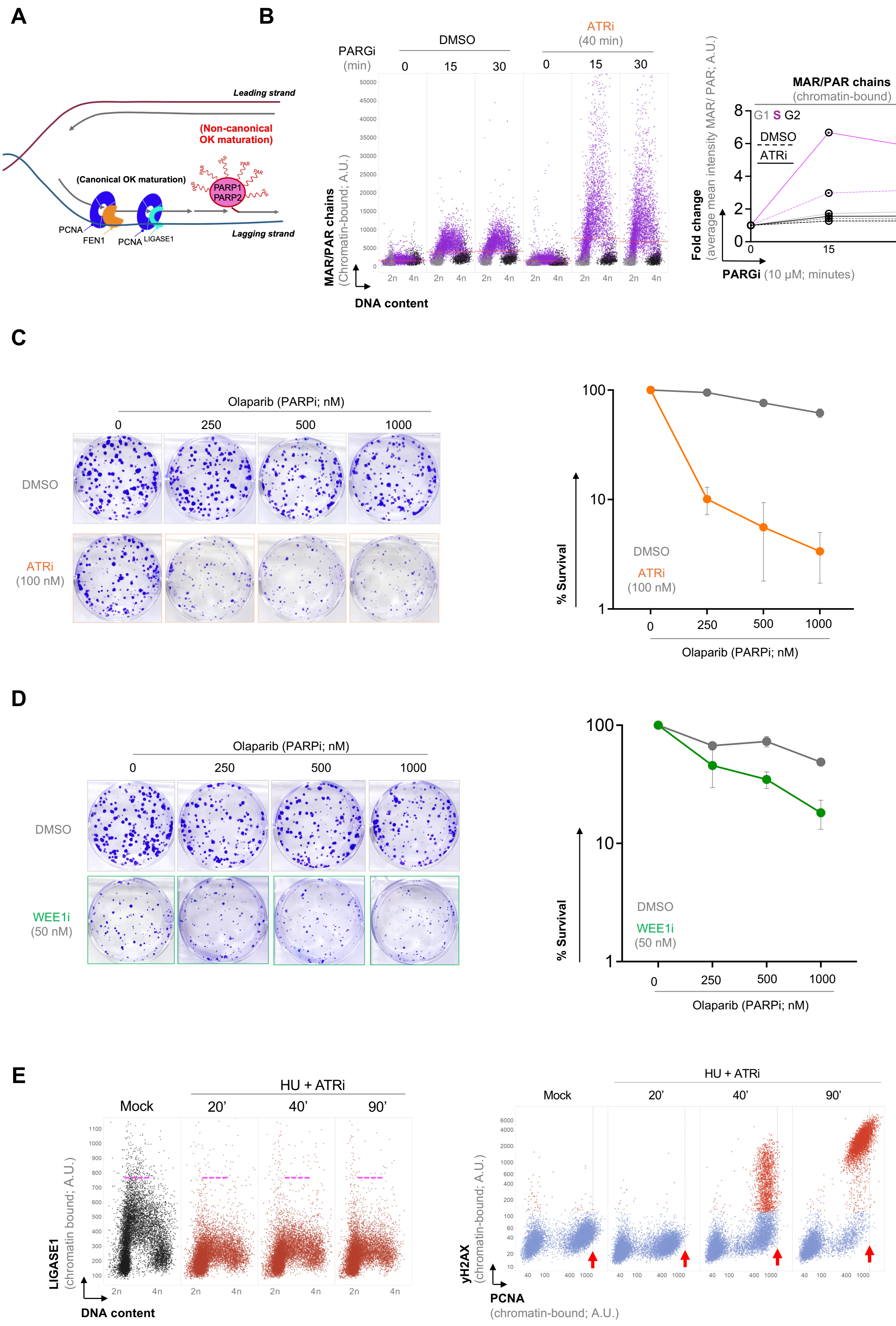
A
B
Leading strand
(Non-canonical OK maturation)
(Canonical OK maturation)
PARP1
PARP2
PAR
PCNA
FEN1
PCNA
LIGASE1
Lagging strand
PARGi (min)
DMSO
ATRi (40 min)
0
15
30
0
15
30
MAR/PAR chains (Chromatin-bound; A.U.)
DNA content
2n
4n
MAR/PAR chains (chromatin-bound)
G1 S G2
DMSO
ATRi
Fold change (average mean intensity MAR/ PAR; A.U.)
PARGi (10 µM; minutes)
C
Olaparib (PARPi; nM)
0
250
500
1000
DMSO
ATRi (100 nM)
% Survival
100
10
1
DMSO
ATRi (100 nM)
Olaparib (PARPi; nM)
D
Olaparib (PARPi; nM)
0
250
500
1000
DMSO
WEE1i (50 nM)
% Survival
DMSO
WEE1i (50 nM)
Olaparib (PARPi; nM)
E
Mock
HU + ATRi
20’
40’
90’
LIGASE1 (chromatin bound; A.U.)
DNA content
HU + ATRi
Mock
20’
40’
90’
γH2AX (chromatin-bound; A.U.)
PCNA (chromatin-bound; A.U.)

Figure S4

**Fig. S4. Non-canonical processing of unligated Okazaki fragments during excessive origin firing.** **(A)** Schematic visualization of non-canonical maturation of Okazaki fragments. PARP1/2 processes Okazaki fragments unligated by FEN1 and LIGASE1 on the lagging strand, indicated by the formation of ADP-ribosylated chains**. (B)** QIBC analysis of the formation of chromatin-bound mono- or poly-ADP ribosylated (MAR/PAR) chains in U2OS cells treated with PARG and ATR inhibitors. 2n corresponding to G1 phase and 4n to G2 phase; >5,000 cells per condition. Fold change in MAR/PAR on chromatin derived from the QIBC data relative to the untreated condition. Survival analysis of cells treated with concentration range of PARP inhibitor Olaparib (250-1000 nM) in combination with ATR (C) or WEE1 (D) inhibition. Values in line plot denote mean ± s.d. **(E)** QIBC analysis of chromatin-bound PCNA and LIGASE1 in cells treated with hydroxyurea (HU) and ATR inhibitor in indicated timepoints. 2n corresponding to G1 phase and 4n to G2 phase; >5,000 cells per condition. Note that LIGASE1 (left) and PCNA (right) do not accumulate on chromatin beyond the level observed in the mock condition, rather show decrease in chromatin loading. Pink dotted lines indicate the approximate maximum levels of each protein.

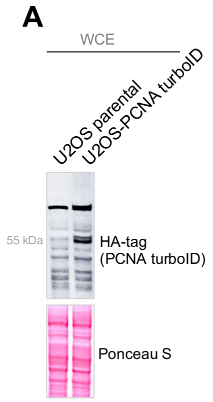


**B**

Log$_{10}$ iBAQ raw values

| | U2OS parental | U2OS-PCNA turbo ID | | | | | |
|---|---|---|---|---|---|---|---|
| | | siControl | | | si*PAF15* | | |
| PCNA | 5.83 | 6.93 | 6.82 | 6.68 | 7.2 | 7.21 | 7.28 |
| CHTF18 | 4.19 | 6.41 | 6.18 | 6.14 | 6.54 | 6.53 | 6.59 |
| RFC1 | 5.16 | 7.02 | 6.89 | 6.80 | 7.23 | 7.40 | 7.40 |
| RFC2 | 0 | 6.26 | 6.14 | 6.06 | 6.45 | 6.39 | 6.45 |
| RFC3 | 0 | 5.50 | 5.44 | 5.37 | 5.71 | 5.69 | 5.65 |
| RFC4 | 0 | 7.72 | 7.58 | 7.5 | 8.03 | 7.99 | 8.03 |
| RFC5 | 6.46 | 6.41 | 6.17 | 6.14 | 6.5 | 6.53 | 6.59 |
| KIAA0101 /PAF15/ PCLAF | 0 | **5.87** | **5.74** | **5.85** | 0 | 0 | 0 |

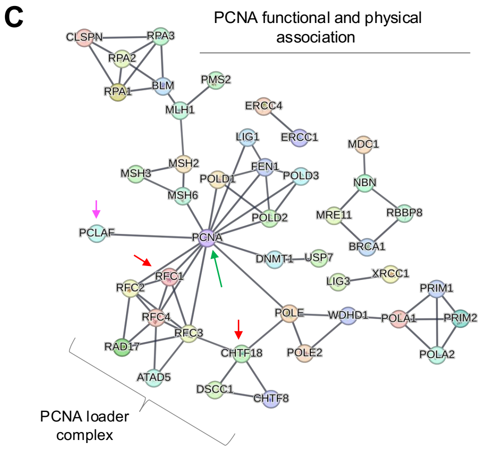


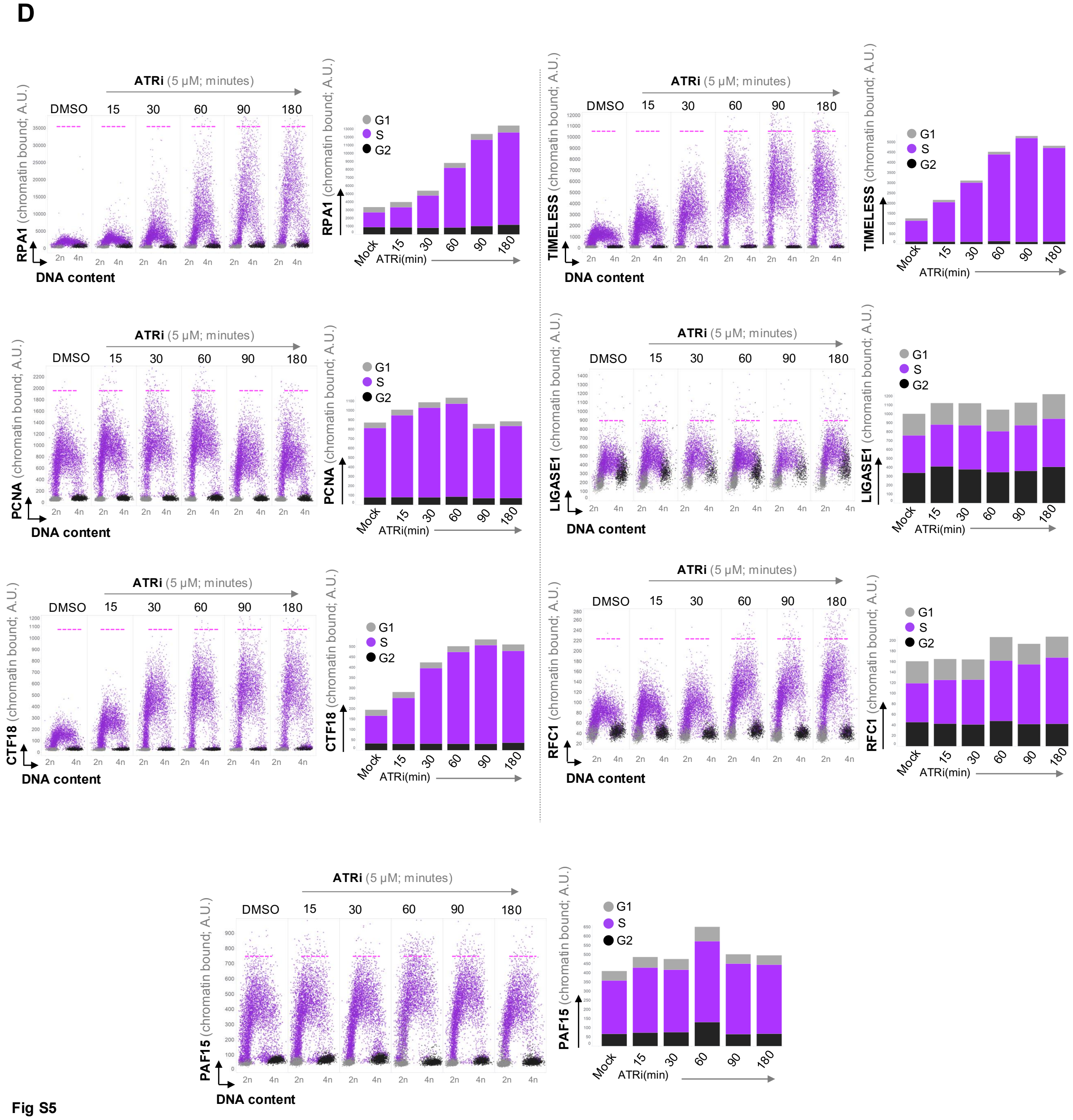


Fig S5

**Fig. S5. Analysis of PCNA-factors during unscheduled origin firing upon ATR inhibition. (A)** Western blot analysis of U2OS cells with turboID-HA-PCNA tag. **(B)** Table showing the results of PCNA proximity-based proteomic screen with TurboID. Raw iBAQ values are shown **(C)**Visualization of PCNA-proximal proteins in String network analysis. The lines depict a direct physical and functional association of PCNA with indicated proteins. **(D)** QIBC analysis and cell cycle specific quantification of indicated chromatin-bound proteins during ATR inhibition at indicated timepoints. 2n corresponding to G1 phase and 4n to G2 phase; >5,000 cells per condition.

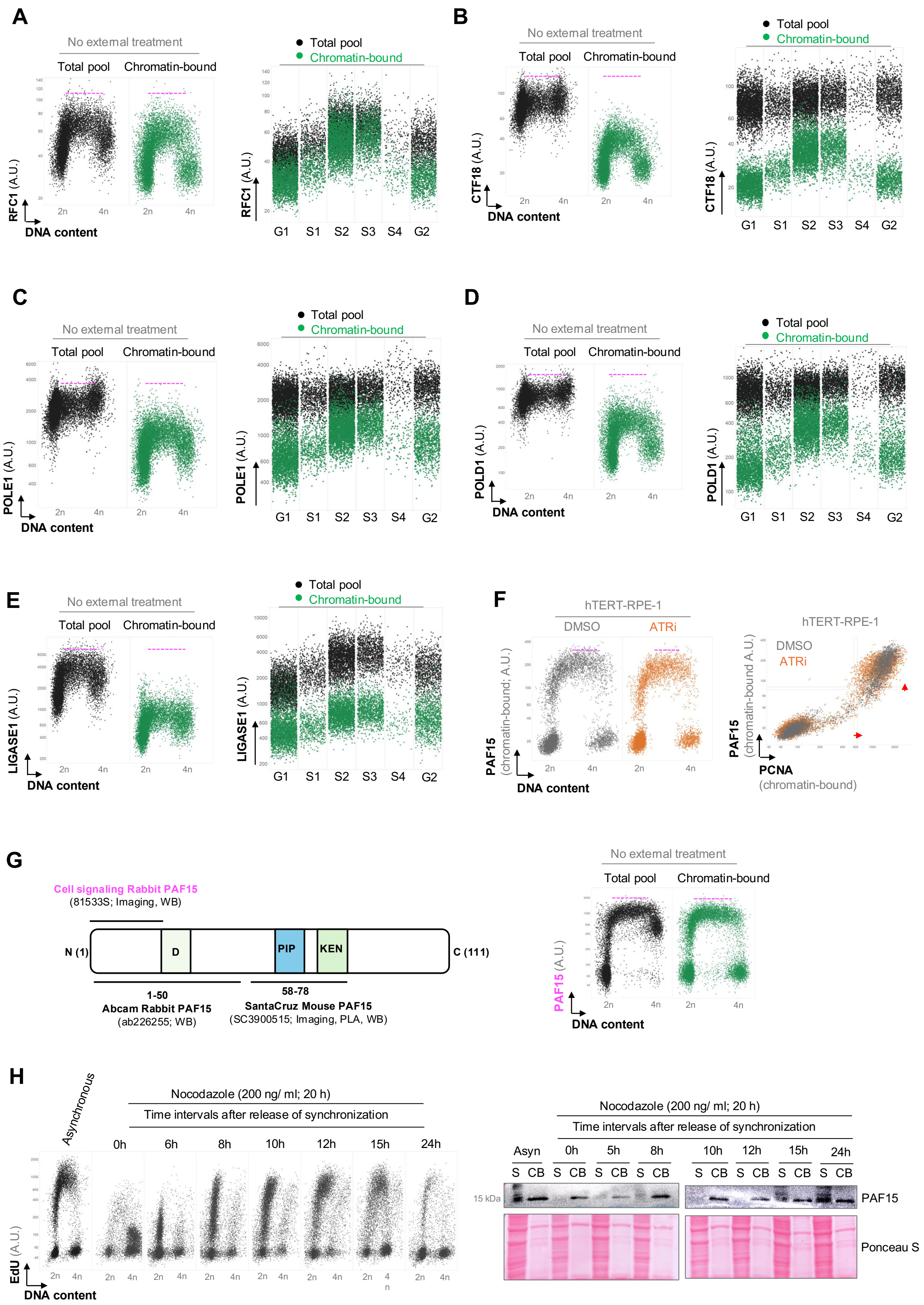
A
No external treatment
Total pool
Chromatin-bound
RFC1 (A.U.)
DNA content
2n
4n
Total pool
Chromatin-bound
G1
S1
S2
S3
S4
G2
B
No external treatment
CTF18 (A.U.)
C
POLE1 (A.U.)
D
POLD1 (A.U.)
E
LIGASE1 (A.U.)
F
hTERT-RPE-1
DMSO
ATRi
PAF15 (chromatin-bound; A.U.)
PCNA (chromatin-bound)
G
Cell signaling Rabbit PAF15
(81533S; Imaging, WB)
N (1)
D
PIP
KEN
C (111)
1-50
Abcam Rabbit PAF15
(ab226255; WB)
58-78
SantaCruz Mouse PAF15
(SC3900515; Imaging, PLA, WB)
PAF15 (A.U.)
H
Asynchronous
Nocodazole (200 ng/ ml; 20 h)
Time intervals after release of synchronization
0h
6h
8h
10h
12h
15h
24h
EdU (A.U.)
Asyn
5h
S
CB
15 kDa
PAF15
Ponceau S

Fig S6

**Fig. S6. A comparison of total and chromatin bound pool of PCNA-associated proteins in S-phase.** QIBC analysis of total and chromatin-bound fractions of RFC1 **(A)**, CTF18 **(B)**, POLE1 **(C)**, POLD1 **(D)** and LIGASE1 **(E)** in naïve U2OS cells. **(F)** QIBC analysis of PAF15 and PCNA in in hTERT-RPE1 cells treated with ATR inhibitor. **(G)** (Left) Binding sites of different PAF15 antibodies. (Right) QIBC analysis of soluble and chromatin-bound fractions of PAF15 in U2OS cells using Cell signaling antibody shown in pink. **(H)** (Left) QIBC analysis of cell cycle with EdU following a 20 h G2M block with microtubule dynamics inhibitor Nocodazole (200 ng/ml) at indicated timepoints post-release. (Right) Western blot analysis of the protein levels of PAF15 soluble (S) and chromatin-bound (CB) fractions. In QIBC plots, 2n corresponding to G1 phase and 4n to G2 phase; >5,000 cells per condition. Pink dotted lines indicate the approximate maximum levels of each protein.

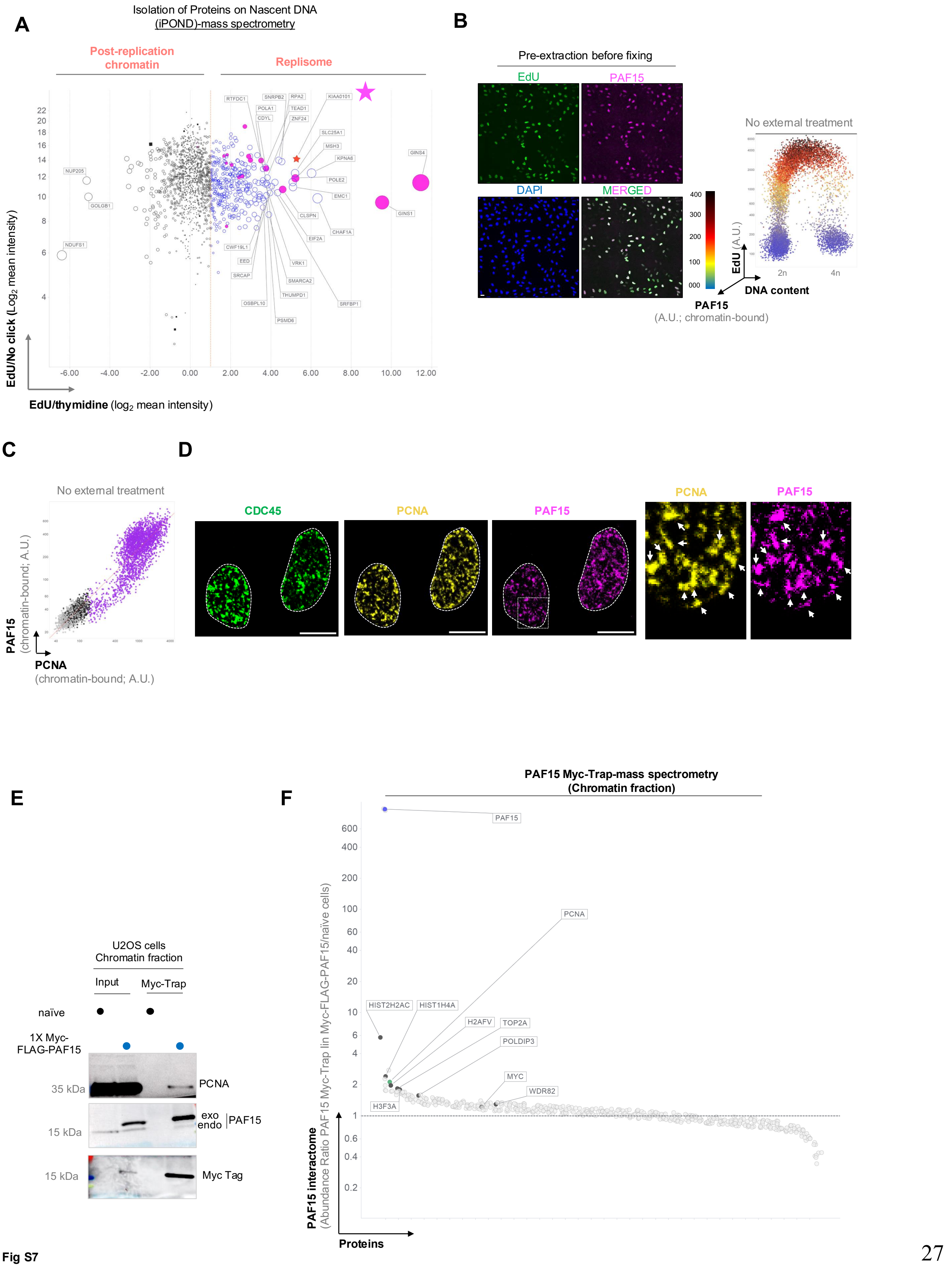
A
Isolation of Proteins on Nascent DNA
(iPOND)-mass spectrometry
Post-replication chromatin
Replisome
EdU/No click (Log2 mean intensity)
EdU/thymidine (log2 mean intensity)
B
Pre-extraction before fixing
EdU
PAF15
DAPI
MERGED
No external treatment
EdU (A.U.)
DNA content
PAF15
(A.U.; chromatin-bound)
C
No external treatment
PAF15
(chromatin-bound; A.U.)
PCNA
(chromatin-bound; A.U.)
D
CDC45
PCNA
PAF15
E
U2OS cells
Chromatin fraction
Input
Myc-Trap
naïve
1X Myc-FLAG-PAF15
35 kDa
PCNA
15 kDa
exo
endo
PAF15
15 kDa
Myc Tag
F
PAF15 Myc-Trap-mass spectrometry
(Chromatin fraction)
PAF15 interactome
(Abundance Ratio PAF15 Myc-Trap lin Myc-FLAG-PAF15/naive cells)
Proteins

Fig S7

**Fig. S7. PAF15 is part of the active replisome a is recruited by PCNA. (A)** Isolation of Proteins on Nascent DNA (iPOND)-mass spectrometry screen. Based on EdU abundancy, the plot is divided into Replisome components (right, in blue) and post-replication chromatin (left, in black). The original data from Somyajit et al. Science 2017 (Reference No: 14) was anlyzed. n = 3 biological replicates. The data show logarithmized ratios of average protein intensities.PAF15 (KIAA0101) as a component of active replisome is marked with a star. **(B)** QIBC analysis of cell cycle distribution of chromatin-bound fraction of PAF15 in U2OS cells. 2n corresponding to G1 phase and 4n to G2 phase; >5,000 cells per condition. **(C)** QIBC analysis of the linear correlation between chromatin-bound fractions of PCNA and PAF15. >5,000 cells. **(D)** Representative 3D confocal microscopy images showing co-localization of PAF15 and PCNA in U2OS cells. Scale bar, 10 µm. **(E)** Immunoprecipitation analysis of PCNA with Myc-trap from FLAG-Myc-tagged PAF15 U2OS cells. **(F)** Mass spectrometry analysis of the enrichment of chromatin-bound proteins trapped by FLAG-Myc-PAF15 in U2OS cells. n = 2 biological replicates. Note that, owing to the missing values of PAF15 peptides in naive U2OS cells (below the detection limit of the MS), the ratio values are very high for PAF15.

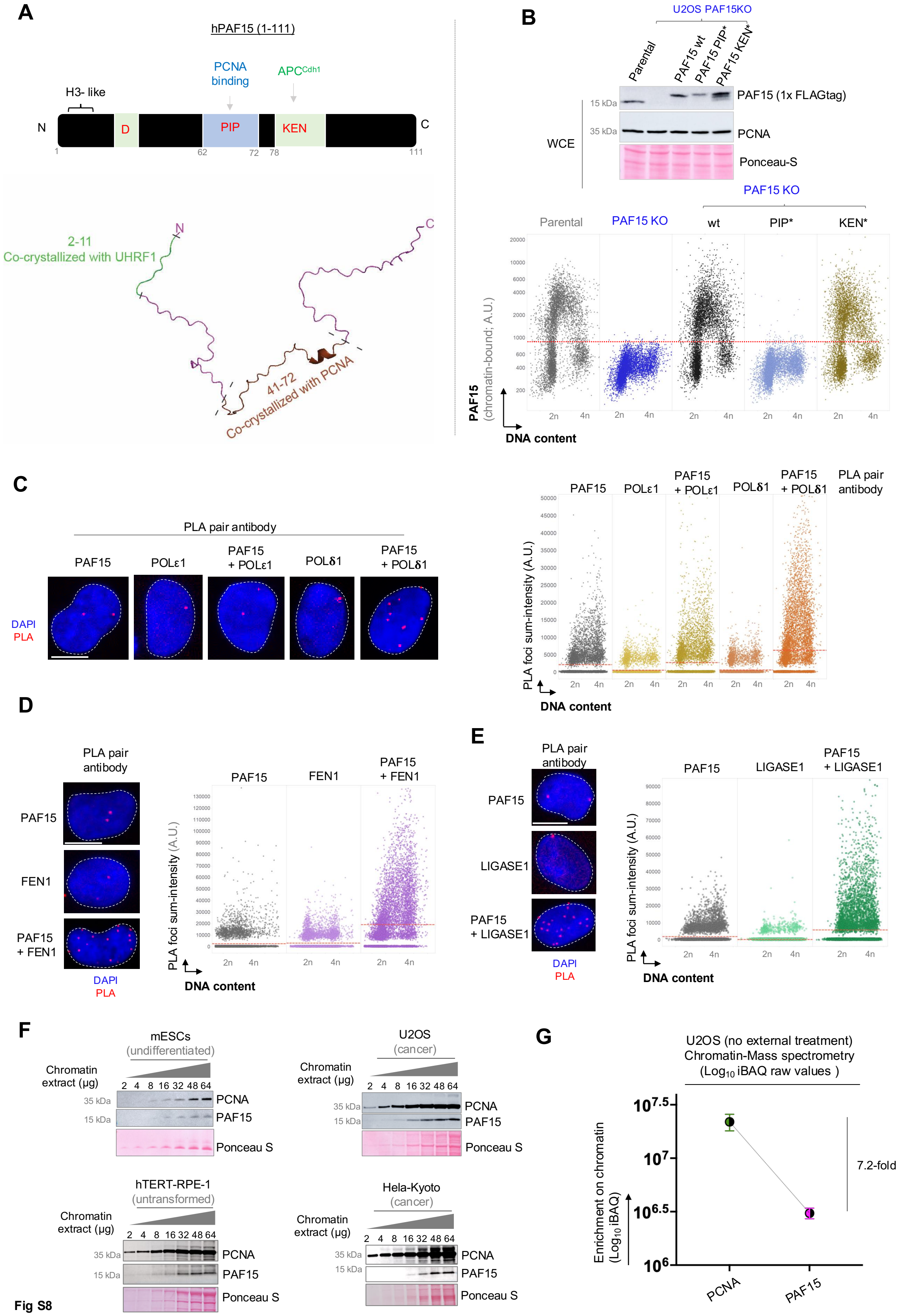

A
hPAF15 (1-111)
PCNA binding
APCCdh1
H3- like
N
D
PIP
KEN
C
1
62
72
78
111
2-11 Co-crystallized with UHRF1
41-72 Co-crystallized with PCNA
B
U2OS PAF15KO
Parental
PAF15 wt
PAF15 PIP*
PAF15 KEN*
15 kDa
35 kDa
PAF15 (1x FLAGtag)
PCNA
Ponceau-S
WCE
PAF15 KO
Parental
PAF15 KO
wt
PIP*
KEN*
PAF15 (chromatin-bound; A.U.)
DNA content
C
PLA pair antibody
PAF15
POLε1
PAF15 + POLε1
POLδ1
PAF15 + POLδ1
DAPI
PLA
PLA foci sum-intensity (A.U.)
DNA content
D
PLA pair antibody
PAF15
FEN1
PAF15 + FEN1
DAPI
PLA
PLA foci sum-intensity (A.U.)
DNA content
E
PLA pair antibody
PAF15
LIGASE1
PAF15 + LIGASE1
DAPI
PLA
PLA foci sum-intensity (A.U.)
DNA content
F
mESCs (undifferentiated)
U2OS (cancer)
hTERT-RPE-1 (untransformed)
Hela-Kyoto (cancer)
Chromatin extract (µg)
2 4 8 16 32 48 64
35 kDa
15 kDa
PCNA
PAF15
Ponceau S
G
U2OS (no external treatment)
Chromatin-Mass spectrometry
(Log10 iBAQ raw values )
Enrichment on chromatin (Log10 iBAQ)
7.2-fold
PCNA
PAF15


Fig S8

**Fig. S8. PAF15 interacts with lagging strand components of the replisome. (A)** (Top) Visualization of the PAF15 molecule with highlighted D, PIP and KEN boxes. (Bottom) Alphafold simulation of the PAF15 structure with highlighted amino-acid sequences important for the binding of UHRF1 and PCNA. **(B)** (Top) Western blot analysis of the PAF15 in whole cell extracts (WCE) of U2OS cells with PAF15 KO and constitutive complentation of mutated PIP/KEN PAF15. (Bottom) QIBC analysis of the chromatin-bound fraction of PAF15 in naïve U2OS and U2OS PAF15 KO stably expressing indicated versions of PAF15**.** 2n corresponding to G1 phase and 4n to G2 phase; >10,000 cells per condition. Red dotted horizon line shows background levels of PAF15 on chromatin. **(C)** (Left) Representative images of PLA assay of PAF15 with indicated proteins in U2OS cells. (Right) Quantification of the PLA foci using QIBC visualized over total DNA content indicating different cell cycle stages. >10,000 cells per condition. **(D and E)** QIBC analysis of PLA of PAF15 with lagging strand components FEN1 and LIGASE1. **(F)** Western blot analysis of chromatin levels of PCNA and PAF15 across different cell types and cell lines. **(G)** Absulute chromatin-bound ratio of PCNA and PAF15 obtained from MS-anlaysis of chromatin fraction from U2OS cells. The ratio is derived from comparing iBAQ values.

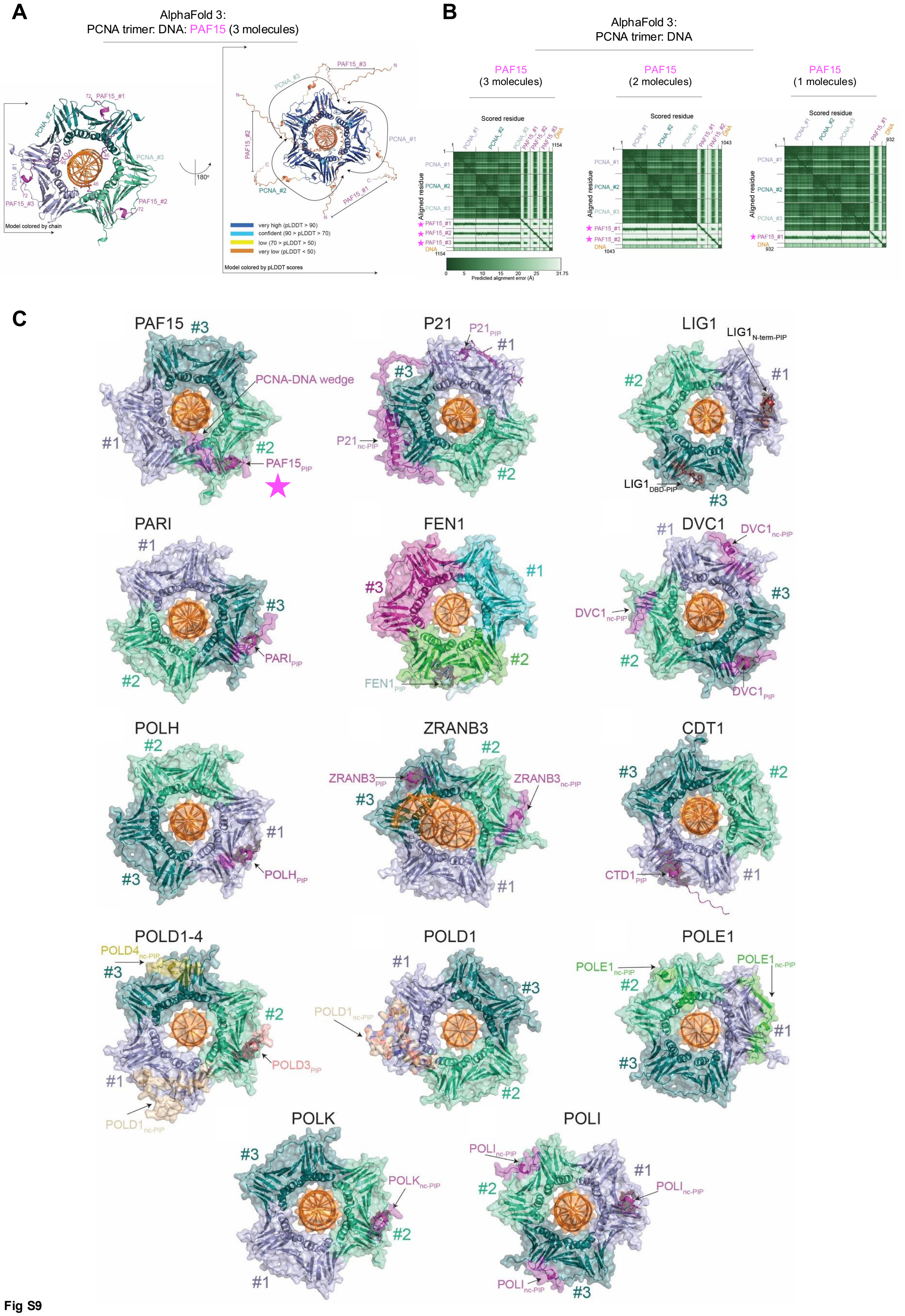
A
AlphaFold 3:
PCNA trimer: DNA: PAF15 (3 molecules)
Model colored by chain
Model colored by pLDDT scores
very high (pLDDT > 90)
confident (90 > pLDDT > 70)
low (70 > pLDDT > 50)
very low (pLDDT < 50)
B
AlphaFold 3:
PCNA trimer: DNA
PAF15
(3 molecules)
PAF15
(2 molecules)
PAF15
(1 molecules)
Scored residue
Aligned residue
Predicted alignment error (Å)
C
PAF15
PCNA-DNA wedge
P21
LIG1
PARI
FEN1
DVC1
POLH
ZRANB3
CDT1
POLD1-4
POLD1
POLE1
POLK
POLI

Fig S9

**Fig. S9. Unique binding mode of PAF15 with PCNA-DNA complex. (A)** AlphaFold model of PAF15-PCNA-DNA complex. Up to 3 molecules of PAF15 can interact with PCNA. PAF15 binds as a molecular wedge between inner PCNA ring and DNA. This structural model is folded with high confidence indicated by high pLDDT scores (right panel). **(B)** The PAE scores of the complex show that all version of one to three molecules of PAF15 are folded with very low predicted alignment errors (<5Å) indicating high confidence at the interaction interfaces between PAF15 and PCNAs highly suggesting a direct interaction. **(C)** AlphaFold simulation of the binding of multiple canonical and non-canonical PCNA-interaction partners. Apart from PAF15 (marked with a star), no other molecules are capable of transversing the inner ring of PCNA together with DNA.

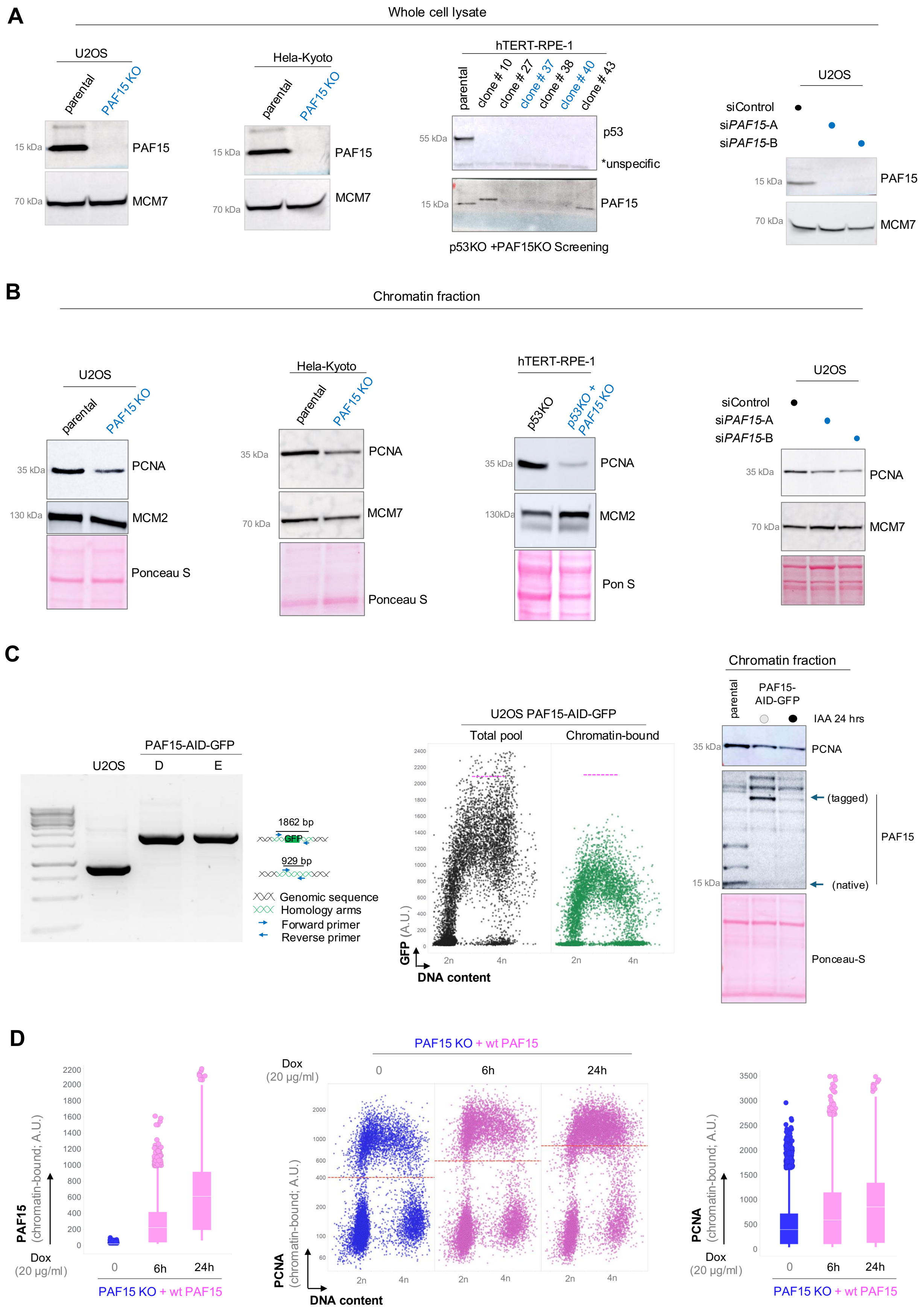
A
Whole cell lysate
U2OS
parental
PAF15 KO
15 kDa
PAF15
70 kDa
MCM7
Hela-Kyoto
parental
PAF15 KO
15 kDa
PAF15
70 kDa
MCM7
hTERT-RPE-1
parental
clone # 10
clone # 27
clone # 37
clone # 38
clone # 40
clone # 43
55 kDa
p53
*unspecific
15 kDa
PAF15
p53KO +PAF15KO Screening
U2OS
siControl
siPAF15-A
siPAF15-B
15 kDa
PAF15
70 kDa
MCM7
B
Chromatin fraction
U2OS
parental
PAF15 KO
35 kDa
PCNA
130 kDa
MCM2
Ponceau S
Hela-Kyoto
parental
PAF15 KO
35 kDa
PCNA
70 kDa
MCM7
Ponceau S
hTERT-RPE-1
p53KO
p53KO + PAF15 KO
35 kDa
PCNA
130kDa
MCM2
Pon S
U2OS
siControl
siPAF15-A
siPAF15-B
35 kDa
PCNA
70 kDa
MCM7
C
PAF15-AID-GFP
U2OS
D
E
1862 bp
GFP
929 bp
Genomic sequence
Homology arms
Forward primer
Reverse primer
U2OS PAF15-AID-GFP
Total pool
Chromatin-bound
GFP (A.U.)
DNA content
2n
4n
Chromatin fraction
parental
PAF15-AID-GFP
IAA 24 hrs
35 kDa
PCNA
(tagged)
PAF15
15 kDa
(native)
Ponceau-S
D
PAF15 KO + wt PAF15
Dox (20 µg/ml)
0
6h
24h
PAF15 (chromatin-bound; A.U.)
PCNA (chromatin-bound; A.U.)
DNA content
2n
4n



Fig S10

**Fig. S10. The loss of PAF15 compromises the stability of PCNA on chromatin.** Western blot validation of the loss of PAF15 **(A)** and destabilization of chromatin-bound PCNA **(B)** in PAF15 KO cell lines (U2OS, HeLa-Kyoto, hTERT-RPE1 p53 KO) and with two PAF15 siRNAs in U2OS cells. (C) (Left) Junction PCR showing homozygous PAF15–AID-GFP tagging; n = 2 biological replicates. (Middle) QIBC analysis of the PAF15-GFP-AID loading during cell cycle. 2n corresponding to G1 phase and 4n to G2 phase; >10,000 cells per condition. Pink dotted lines indicate the approximate maximum levels of PAF15 protein. (Right) Western blot analysis of the stability of chromatin-bound PCNA in U2OS PAF15-GFP-AID and upon inducible degradation of PAF15 with natural degron indole-3-acetic acid (IAA) for 24 h. (D) QIBC analysis of the chromatin-bound PAF15 (Left) and PCNA (Middle and right) in U2OS cells depleted of endogenous PAF15 with Dox-inducible overexpression of PAF15 for indicated timepoints. 2n corresponding to G1 phase and 4n to G2 phase; >10,000 cells per condition. Dotted horizon line shows average levels of PCNA on chromatin.

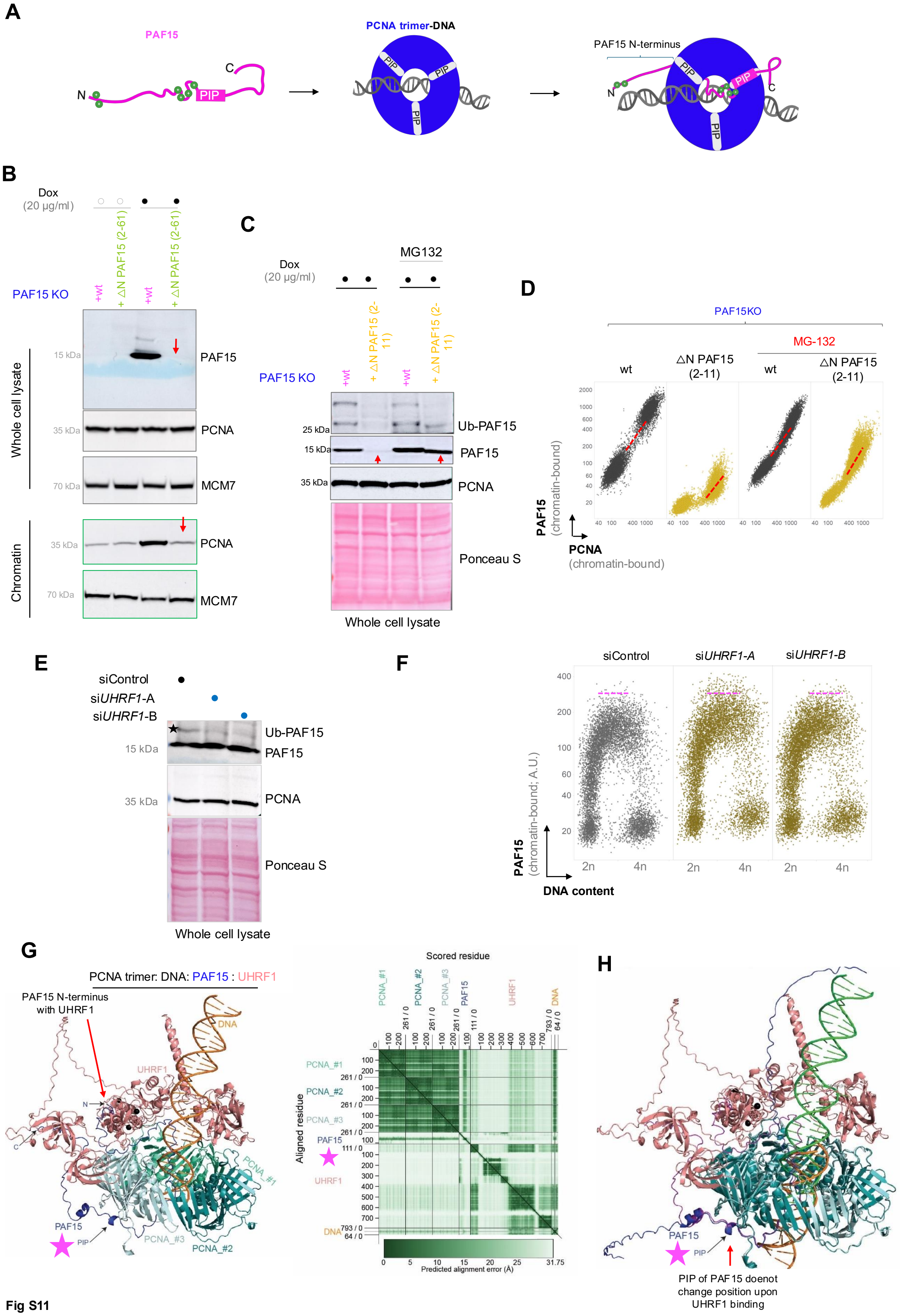
A
PAF15
PCNA trimer-DNA
PAF15 N-terminus
PIP
N
C
B
Dox
(20 µg/ml)
PAF15 KO
+wt
+ ΔN PAF15 (2-61)
PAF15
PCNA
MCM7
Whole cell lysate
Chromatin
C
MG132
+ ΔN PAF15 (2-11)
Ub-PAF15
Ponceau S
D
PAF15KO
MG-132
wt
ΔN PAF15 (2-11)
PAF15 (chromatin-bound)
PCNA (chromatin-bound)
E
siControl
siUHRF1-A
siUHRF1-B
F
PAF15 (chromatin-bound; A.U.)
DNA content
G
PCNA trimer: DNA: PAF15 : UHRF1
PAF15 N-terminus with UHRF1
UHRF1
DNA
PCNA_#1
PCNA_#2
PCNA_#3
Scored residue
Aligned residue
Predicted alignment error (Å)
H
PIP of PAF15 doenot change position upon UHRF1 binding

Fig S11

**Fig. S11. PAF15 directly regulates PCNA dynamics on chromatin. (A)** Cartoon visualization of PAF15 binding to PCNA-DNA complex. Green dots at PIP box and N-terminus indicate positively charged sites. **(B)** Western blot analysis of PAF15 and PCNA in U2OS PAF15 KO with overexpression of either WT or N-terminus truncated versions of PAF15. Red arrows indicate the direct cause of PAF15 loss on stability of chromatin-bound PCNA. Western blot **(C)** and QIBC **(D)** analysis of PAF15 and PCNA in U2OS PAF15 KO with overexpression of either WT or N-terminus truncated versions of PAF15 with the addition of proteasomal inhibitor MG132. $n > 5{,}000$ cells each condition. Red dotted lines in QIBC scatter plots indicate the approximate linear correlation between PAF15 and PCNA. **(E)** Western blot analysis of PAF15 and PCNA in U2OS treated with siRNAs targeting the ubiquitin ligase UHRF1. **(F)** QIBC analysis of PAF15 on chromatin in U2OS treated with siRNAs targeting UHRF1. $n > 5{,}000$ cells for each condition. Pink dotted lines in QIBC scatter plots indicate the approximate maximum levels of each protein. **(G)** AlphaFold prediction of a protein complex formed by three PCNAs, one PAF15, one UHRF1, one double stranded DNA and three $Zn^{2+}$ ions (black spheres) with high confidence (prediction alignment error below 10Å). (H) The alignment of PCNA-PAF15-DNA complexes with or without UHRF1 (root mean square deviation (RMSD) = 0.02, shown in dark blue and purple color) does not change the position of the PAF15PIP with respect to PCNA.

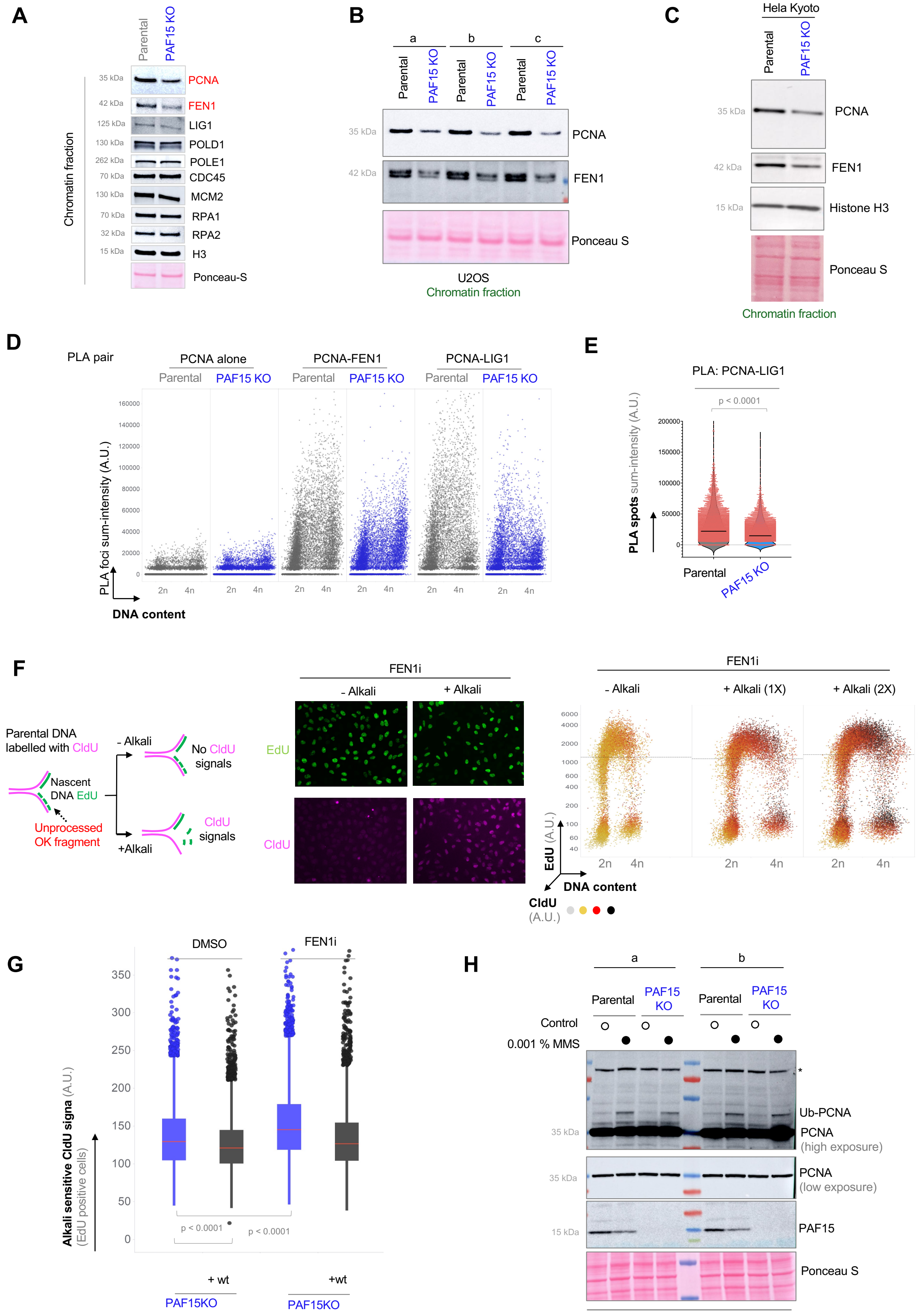
A
Parental
PAF15 KO
Chromatin fraction
PCNA
FEN1
LIG1
POLD1
POLE1
CDC45
MCM2
RPA1
RPA2
H3
Ponceau-S
B
a
b
c
Parental
PAF15 KO
PCNA
FEN1
Ponceau S
U2OS
Chromatin fraction
C
Hela Kyoto
Parental
PAF15 KO
PCNA
FEN1
Histone H3
Ponceau S
Chromatin fraction
D
PLA pair
PCNA alone
PCNA-FEN1
PCNA-LIG1
Parental
PAF15 KO
PLA foci sum-intensity (A.U.)
DNA content
E
PLA: PCNA-LIG1
p < 0.0001
PLA spots sum-intensity (A.U.)
Parental
PAF15 KO
F
Parental DNA labelled with CldU
Nascent DNA EdU
Unprocessed OK fragment
- Alkali
+Alkali
No CldU signals
CldU signals
FEN1i
- Alkali
+ Alkali
EdU
CldU
FEN1i
- Alkali
+ Alkali (1X)
+ Alkali (2X)
EdU (A.U.)
DNA content
CldU (A.U.)
G
DMSO
FEN1i
Alkali sensitive CldU signa (A.U.)
(EdU positive cells)
p < 0.0001
p < 0.0001
+ wt
+wt
PAF15KO
PAF15KO
H
a
b
Parental
PAF15 KO
Control
0.001 % MMS
Ub-PCNA
PCNA
(high exposure)
PCNA
(low exposure)
PAF15
Ponceau S
WCE

**Fig S12**

**Fig. S12. PAF15-mediated stabilization of lagging strand-associated PCNA is crucial for Okazaki fragment maturation. (A)** Selected proteins from the PCNA-turboID proximity proteomics forming core-replisome were analyzed by Western blot of purified chromatin fractions in U2OS parental and PAF15 KO cells. Western blot analysis of chromatin fraction of indicated proteins in different clones of U2OS PAF15 KO **(B)** and HeLa Kyoto PAF15KO cells **(C)**. **(D)** QIBC analysis of PLA foci of PAF15 with lagging strand proteins FEN1 and LIG1 in PAF15KO U2OS cells. n > 10,000 cells **(E)** QIBC quantification of PLA foci of PCNA-LIG1 pair in PAF15KO U2OS cells. n > 5,000 S-phase cells. **(F)** (Left) Schematic visualization of the CldU detection of Alkali sensitive unligated Okazaki fragments. (Middle and right) QIBC analysis of Alkali sensitive unligated Okazaki fragments in U2OS cells treated with FEN1 inhibitor (20 μM). n > 10,000 cells. Horizontal lines in QIBC scatter plots indicate the average values for EdU, which remains unchanged while CldU signals increase upon alkali treatment (shown by color heatmap). **(G)** Comparison of total CldU signal in S-phase (EdU positive) cells between DMSO and FEN1 inhibitor treated U2OS PAF15 KO cells overexpressing PAF15. **(H)** Western blot analysis of PCNA and PAF15 in U2OS parental and PAF15KO cells treated with 0.001 % methyl methane sulfonate (MMS) alkylating agent.

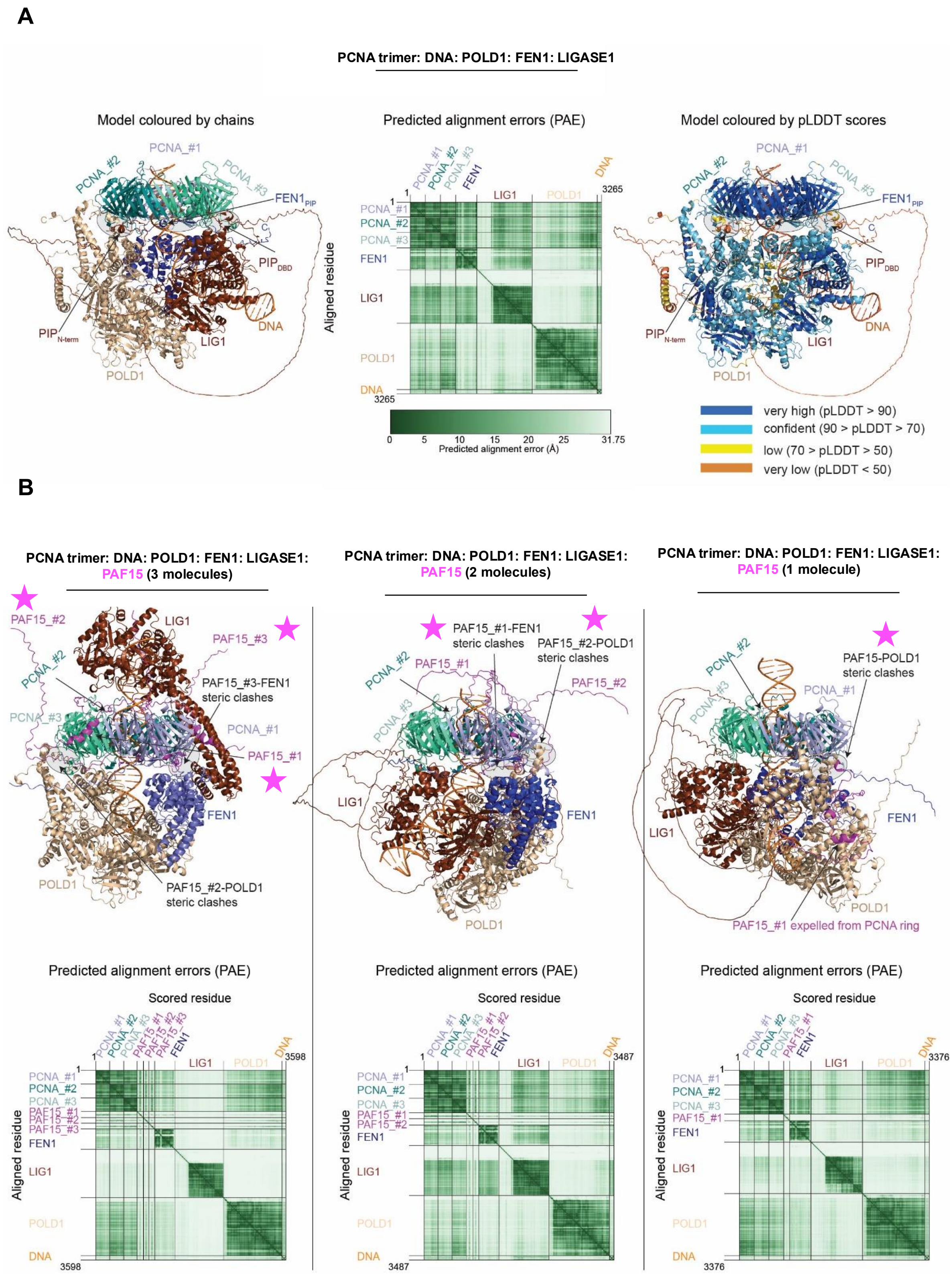

A
PCNA trimer: DNA: POLD1: FEN1: LIGASE1
Model coloured by chains
Predicted alignment errors (PAE)
Model coloured by pLDDT scores
PCNA_#1
PCNA_#2
PCNA_#3
FEN1
FEN1PIP
PIPDBD
PIPN-term
DNA
LIG1
POLD1
Aligned residue
3265
0 5 10 15 20 25 31.75
Predicted alignment error (Å)
very high (pLDDT > 90)
confident (90 > pLDDT > 70)
low (70 > pLDDT > 50)
very low (pLDDT < 50)
B
PCNA trimer: DNA: POLD1: FEN1: LIGASE1: PAF15 (3 molecules)
PCNA trimer: DNA: POLD1: FEN1: LIGASE1: PAF15 (2 molecules)
PCNA trimer: DNA: POLD1: FEN1: LIGASE1: PAF15 (1 molecule)
PAF15_#1
PAF15_#2
PAF15_#3
PAF15_#3-FEN1 steric clashes
PAF15_#2-POLD1 steric clashes
PAF15_#1-FEN1 steric clashes
PAF15-POLD1 steric clashes
PAF15_#1 expelled from PCNA ring
Predicted alignment errors (PAE)
Scored residue
Aligned residue
3598
3487
3376


Fig S13

**Fig. S13. PAF15 controls the access of FEN1, LIG1 and POLD1 to PCNA. (A)** AlphaFold models a heptameric PCNA-FEN1-LIG1-POLD1-DNA complex (Left) where each of the DNA lagging strand factors occupy a vacant PCNA molecule (highlighted by grey ellipses). The N-terminal PIP motif of LIG1 is folded close to PCNA_#3 but kept away from the docking site by POLD1. Thus, LIG1 only uses the $PIP_{DBD}$ motif to access a vacant PCNA. (Middle) The PAE scores of the model shows low distance errors (<10Å) occurring between PCNA_#1 and FEN1, PCNA_#2 and LIG1, PCNA_#3 and POLD1, FEN1 and LIG1. (Right) Overall, the structural model is predicted with good confidence (pLDDT>70) except the intrinsically disordered regions of LIG1 spanning residues 1-260, and POLD1 comprising residues 1-74. **(B)** Addition of three, two or one molecules of PAF15 (indicated by star) to the heptameric PCNA-FEN1-LIG1-POLD1-DNA complex re-organizes the PCNA interactors at the ring. (Right) The predicted model of the heptameric PCNA-FEN1-LIG1-POLD1-DNA in the presence of three PAF15s shows complete displacement of FEN1 and LIG1 from PCNA (PAE scores >31.75Å). AlphaFold preserves the folding of POLD1 and one PAF15 at the same PCNA monomer, reflected by the low PAE scores, but at the cost of creating severe steric clashes. FEN1 is also docked closer to the PCNA_#1 in steric clashes with PAF15_#1. (Middle) Two PAF15 molecules predicted with the heptameric PCNA-FEN1-LIG1-POLD1-DNA are docked onto their cognate PCNAs and displace FEN1 (high PAE scores on the bottom panel). Each PAF15 molecule is in steric clash with FEN1 and POLD1, respectively. (Right) One PAF15 molecule predicted with the heptameric PCNA-FEN1-LIG1-POLD1-DNA complex creates disturbances at the PCNA ring. Both PAF15 and LIG1 are folded away from the PCNA ring as indicated by the high PAE scores on the bottom panel. Moreover, FEN1 loses contact to LIG1 (very high PAE scores).

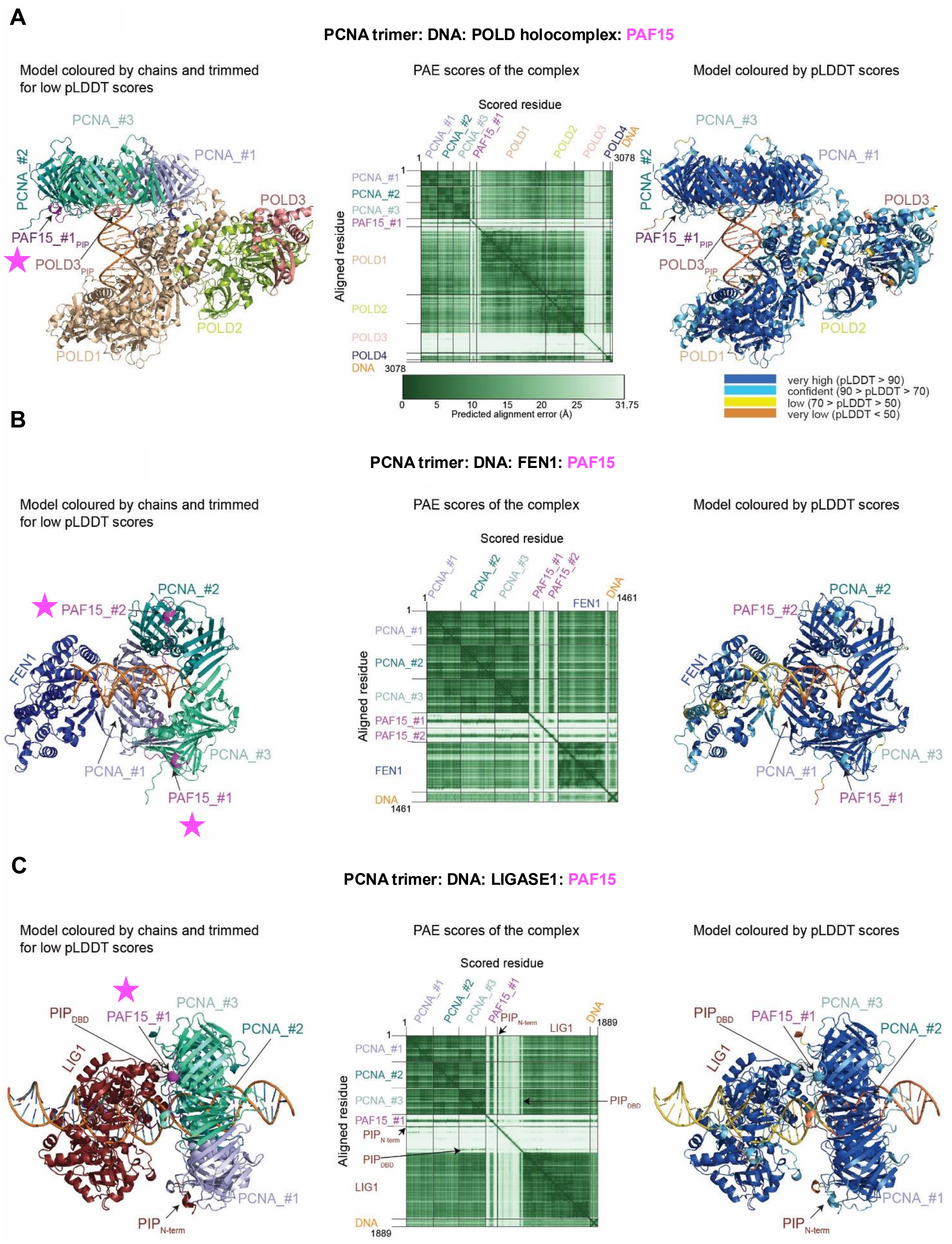
A
PCNA trimer: DNA: POLD holocomplex: PAF15
Model coloured by chains and trimmed for low pLDDT scores
PAE scores of the complex
Model coloured by pLDDT scores
PCNA_#3
PCNA_#2
PCNA_#1
POLD3
PAF15_#1PIP
POLD3PIP
POLD2
POLD1
Scored residue
Aligned residue
PCNA_#1
PCNA_#2
PCNA_#3
PAF15_#1
POLD1
POLD2
POLD3
POLD4
DNA
3078
0 5 10 15 20 25 31.75
Predicted alignment error (Å)
very high (pLDDT > 90)
confident (90 > pLDDT > 70)
low (70 > pLDDT > 50)
very low (pLDDT < 50)
B
PCNA trimer: DNA: FEN1: PAF15
Model coloured by chains and trimmed for low pLDDT scores
PAE scores of the complex
Model coloured by pLDDT scores
PAF15_#2
PCNA_#2
FEN1
PCNA_#1
PCNA_#3
PAF15_#1
Scored residue
Aligned residue
1461
C
PCNA trimer: DNA: LIGASE1: PAF15
Model coloured by chains and trimmed for low pLDDT scores
PAE scores of the complex
Model coloured by pLDDT scores
PIPDBD
PAF15_#1
PCNA_#3
PCNA_#2
LIG1
PCNA_#1
PIPN-term
Scored residue
Aligned residue
1889



Fig S14

**Fig. S14. The occupancy of PCNA by PAF15 controls the access of lagging strand PCNA-interactors.** (A) Alphafold predicts a nonameric PCNA-PAF15-POLD-DNA complex. (Left) PAF15 occupies one PCNA monomer whilst POLD1 and POLD3 saturates the other two. The model is predicted with high confidence as revealed by predicted alignment errors (PAE<10Å, Middle) and high per-residues confidence scores (pLDDT>90, Right). Few intrinsically disordered regions that comprises the residues 1-74 of POLD1, 457-467 of POLD2, 144-455 of POLD3 and 1-40 of POLD4 are predicted with very low pLDDT scores and therefore omitted from the model for clarity. **(B)** The structural model of a heptameric PCNA-PAF15-FEN1-DNA shows that docking of FEN1 to one PCNA monomer allows PAF15 two occupy the other two (Left). The model is predicted with very high confidence as highlighted by low PAEs (Middle) and high pLDDTs (Right). **(C)** AlphaFold predicts a hexameric PCNA-PAF15-LIG-DNA complex where LIG1 binds the PCNA trimer by its PIPN-terminus and PIPDBD domains allowing one PAF15 molecule to access the unoccupied PCNA (Left). The model is predicted with high confidence as revealed by low PAE scores (Middle) and high pLDDT scores (Right). Intrinsically disordered regions present at the N- and C- termini of PAF15 and between the residues 1-260 of LIG1 are predicted with very low pLDDT scores and thus omitted from the model for clarity.

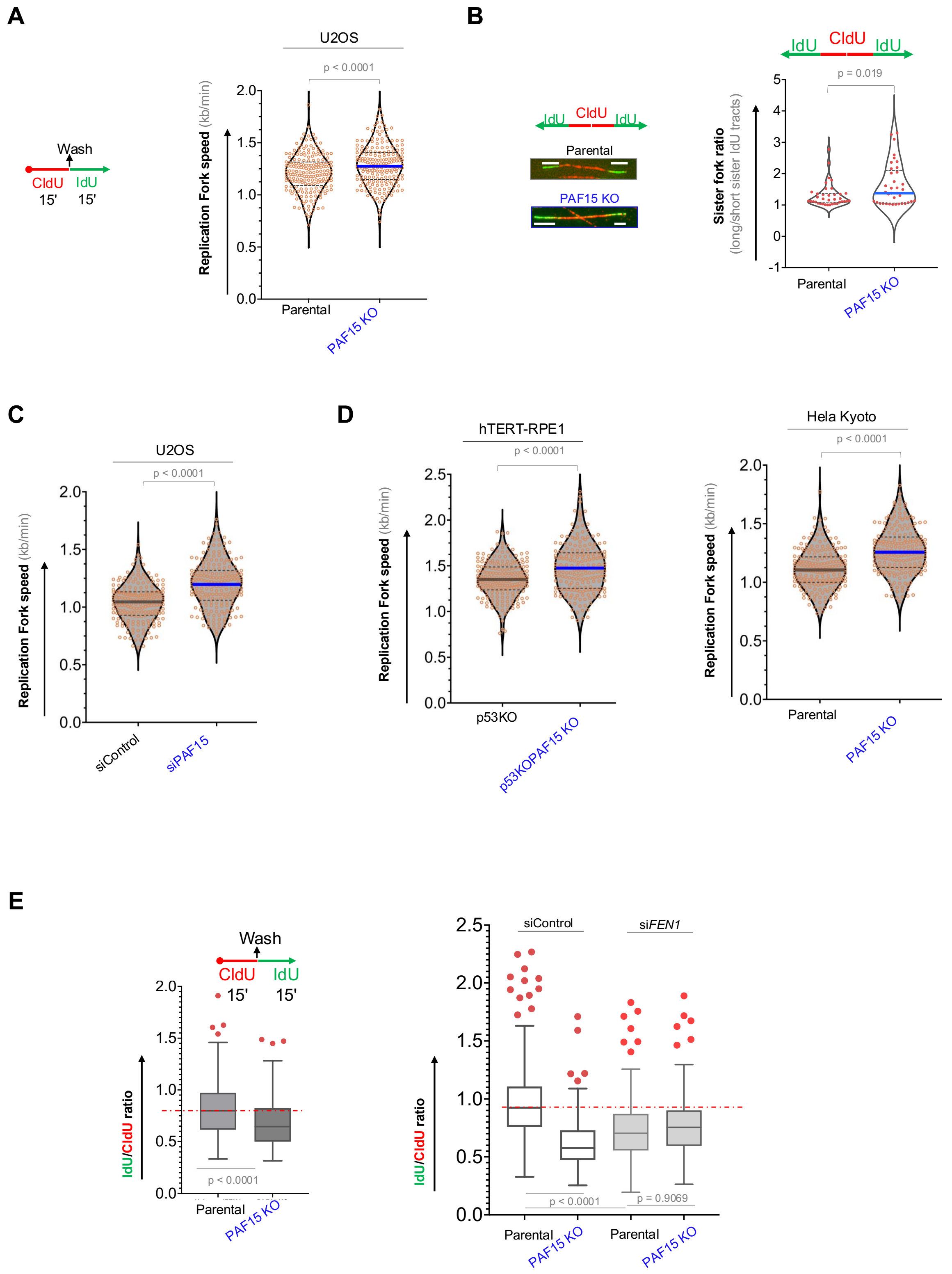
A
Wash
CldU 15'
IdU 15'
U2OS
p < 0.0001
Replication Fork speed (kb/min)
Parental
PAF15 KO
B
IdU CldU IdU
Parental
PAF15 KO
p = 0.019
Sister fork ratio (long/short sister IdU tracts)
C
U2OS
p < 0.0001
Replication Fork speed (kb/min)
siControl
siPAF15
D
hTERT-RPE1
p < 0.0001
p53KO
p53KOPAF15 KO
Hela Kyoto
p < 0.0001
Replication Fork speed (kb/min)
Parental
PAF15 KO
E
Wash
CldU 15'
IdU 15'
IdU/CldU ratio
p < 0.0001
Parental
PAF15 KO
siControl
siFEN1
IdU/CldU ratio
p < 0.0001
p = 0.9069
Parental
PAF15 KO
Parental
PAF15 KO



Fig S15

**Fig. S15. PAF15 loss accelerates fork speed and heightens replication stress in S phase. (A)** Replication fork speed comparison in indicated cell lines between parental and PAF15KO cells. n = 200 fibers per condition. P values were determined by one-way ANOVA with Tukey's test. **(B)** Analysis of sister fork asymmetry in U2OS cells. n = 50 bidirectional fibers per condition. P values were determined by one-way ANOVA with Tukey's test. **(C)** and **(D)** Replication fork speed in indicated cells. n = 200 fibers per condition. P values were determined by one-way ANOVA with Tukey's test. (E) Fork ratio derived by dividing the length of DNA tracts labeled by IdU and CldU in indicated U2OS cells. n = 200 fibers per condition. P values were determined by one-way ANOVA with Tukey's test.

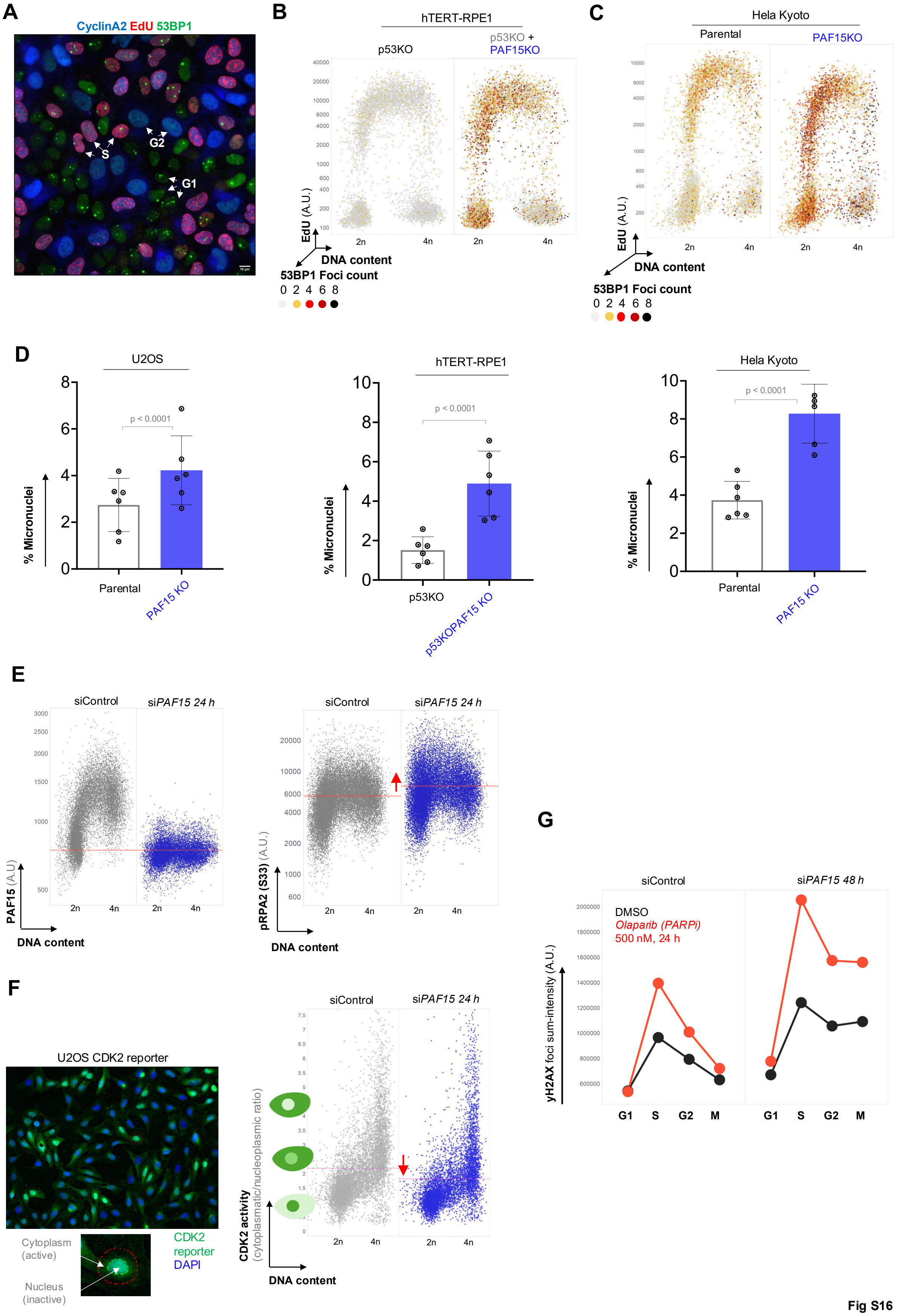
A
CyclinA2 EdU 53BP1
S
G2
G1
B
hTERT-RPE1
p53KO
p53KO + PAF15KO
EdU (A.U.)
DNA content
53BP1 Foci count
0 2 4 6 8
2n
4n
C
Hela Kyoto
Parental
PAF15KO
D
U2OS
hTERT-RPE1
Hela Kyoto
% Micronuclei
p < 0.0001
Parental
PAF15 KO
p53KO
p53KOPAF15 KO
E
siControl
siPAF15 24 h
PAF15 (A.U)
pRPA2 (S33) (A.U.)
DNA content
F
U2OS CDK2 reporter
Cytoplasm (active)
Nucleus (inactive)
CDK2 reporter
DAPI
CDK2 activity (cytoplasmatic/nucleoplasmic ratio)
G
siControl
siPAF15 48 h
DMSO
Olaparib (PARPi)
500 nM, 24 h
γH2AX foci sum-intensity (A.U.)
G1
S
G2
M

Fig S16

**Fig. S16. The loss of PAF15 leads to recruitment of non-canonical pathways of Okazaki fragment maturation. (A)** Representative microscopy image of U2OS cells immunostained for Cyclin A2 and 53BP1 plus EdU staining for S-phase visualization. QIBC of 53BP1 foci/nuclear bodies in p53KO **(B)** and HeLa Kyoto **(C)** parental and PAF15KO cells labeled for EdU and DAPI to stratify cell cycle progression (n > 10,000 cells for each condition; colors indicate the number of 53BP1 nuclear bodies per nucleus. **(D)** Quantification of micronuclei formation in indicated parental and PAF15KO cell lines. n = 500 cells per condition **(E)** QIBC analysis of PAF15 and pRPA2 in U2OS cells with depletion of PAF15 for 24 h. Horizontal lines in PAF15 QIBC blots show effective PAF15 depletion bringing PAF15 to background levels. Horizontal lines in pRPA2 QIBC blots show average value. 2n: G1, 4n: G2; > 10,000 cells per condition. **(F)** Visual representation of CDK2 reporter system in U2OS cells. The cytoplasmic/nuclear ratio gradually increases during cell cycle progression from G1 to G2M. Horizontal lines show average value. 2n: G1, 4n: G2; > 10,000 cells per condition. **(G)** QIBC analysis of yH2AX foci in U2OS PAF15 depleted cells treated with PARP inhibitor Olaparib (500 nM, 24 h). 2n: G1, 4n: G2; > 10,000 cells per condition. Cell cycle gating was done based on DNA content (G1 and G2), EdU

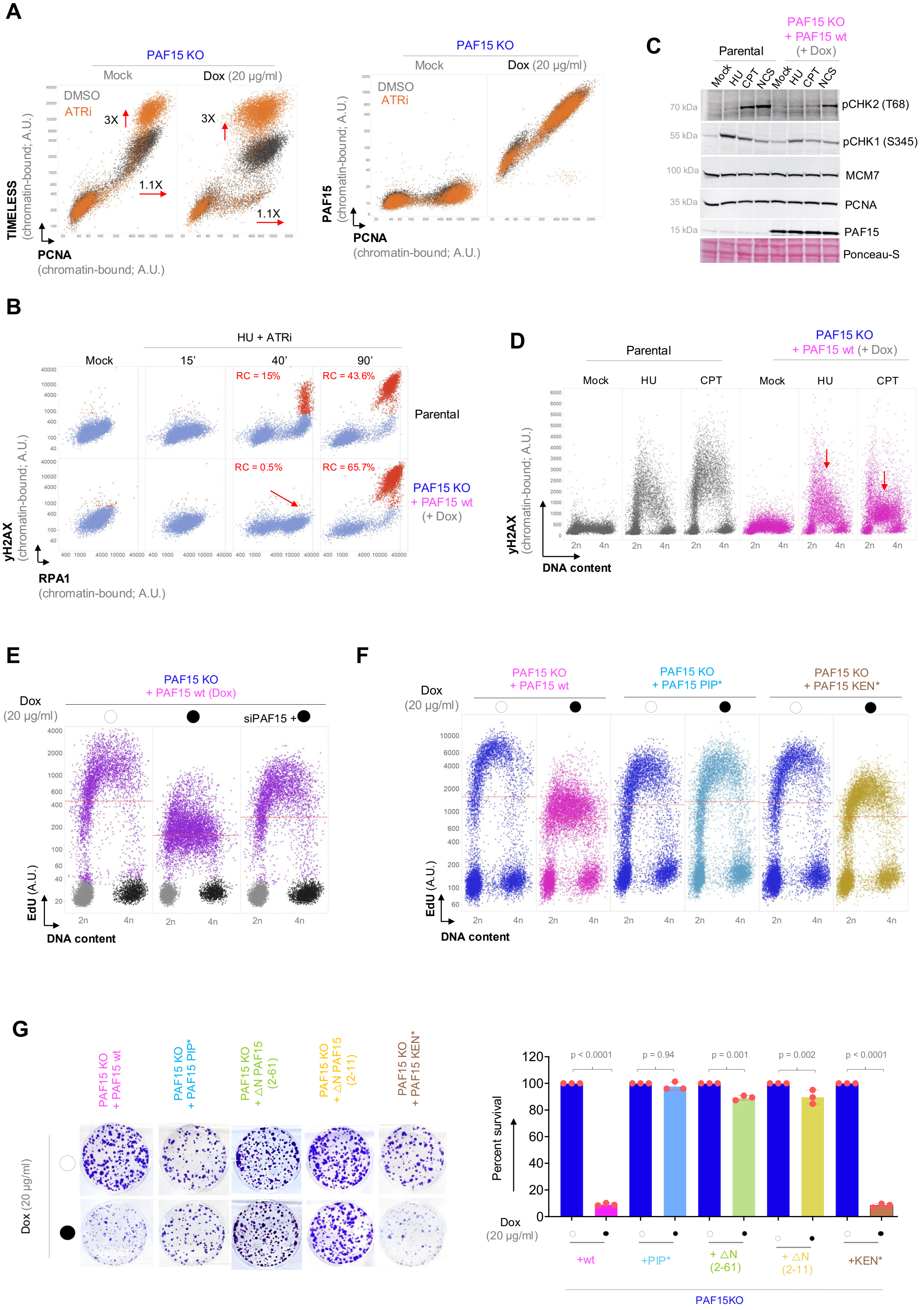
A
PAF15 KO
Mock
Dox (20 µg/ml)
DMSO
ATRi
3X
1.1X
TIMELESS (chromatin-bound; A.U.)
PCNA (chromatin-bound; A.U.)
PAF15 (chromatin-bound; A.U.)
C
Parental
PAF15 KO + PAF15 wt (+ Dox)
Mock HU CPT NCS Mock HU CPT NCS
70 kDa pCHK2 (T68)
55 kDa pCHK1 (S345)
100 kDa MCM7
35 kDa PCNA
15 kDa PAF15
Ponceau-S
B
HU + ATRi
Mock 15' 40' 90'
RC = 15%
RC = 43.6%
Parental
RC = 0.5%
RC = 65.7%
PAF15 KO + PAF15 wt (+ Dox)
γH2AX (chromatin-bound; A.U.)
RPA1 (chromatin-bound; A.U.)
D
Parental
PAF15 KO + PAF15 wt (+ Dox)
Mock HU CPT Mock HU CPT
γH2AX (chromatin-bound; A.U.)
DNA content
E
Dox (20 µg/ml)
PAF15 KO + PAF15 wt (Dox)
siPAF15 +
EdU (A.U.)
DNA content
F
Dox (20 µg/ml)
PAF15 KO + PAF15 wt
PAF15 KO + PAF15 PIP*
PAF15 KO + PAF15 KEN*
EdU (A.U.)
DNA content
G
PAF15 KO + PAF15 wt
PAF15 KO + PAF15 PIP*
PAF15 KO + ΔN PAF15 (2-61)
PAF15 KO + ΔN PAF15 (2-11)
PAF15 KO + PAF15 KEN*
Dox (20 µg/ml)
Percent survival
p < 0.0001
p = 0.94
p = 0.001
p = 0.002
p < 0.0001
+wt
+PIP*
+ ΔN (2-61)
+ ΔN (2-11)
+KEN*
PAF15KO

Figure S17

**Fig. 17. Overexpression of PAF15 severely compromises cell viability.** (A) QIBC analysis of chromatin fractions of TIMELESS (Left) and PAF15 (Right) with PCNA in U2OS cells overexpressing PAF15 treated with ATR inhibitor. n > 10,000 cells per condition. (B) QIBC analysis of chromatin fractions of yH2AX and RPA1 in parental and PAF15 overexpressing U2OS cells treated with combination of hydroxyurea (HU) and ATR inhibitor. n> 10,000 cells per condition. (C) Western blot analysis of indicated protein in parental and PAF15 overexpressing U2OS cells treated with 2mM hydroxyurea (HU, replication stress), 50 nM camptothecin (CPT, Topoisomerase 1 inhibitor) and 50 ng/ml neocarzinostatin (NCS, induction of double-stranded DNA breaks). (D) QIBC analysis of yH2AX in parental and PAF15 overexpressing U2OS cells treated with HU and CPT. n> 10,000 cells per condition. (E) QIBC analysis of EdU in PAF15KO and PAF15 overexpressing U2OS cells with the addition of PAF15 siRNA. n> 10,000 cells per condition. (F) QIBC analysis of EdU in U2OS cells overexpressing WT, PIP-box mutated and KEN-box mutated PAF15. n> 10,000 cells per condition. (G) Survival analysis in U2OS cells overexpressing WT, PIP-box mutated, N-terminus truncated and KEN-box mutated PAF15. P values were determined by one-way ANOVA with Tukey's test.

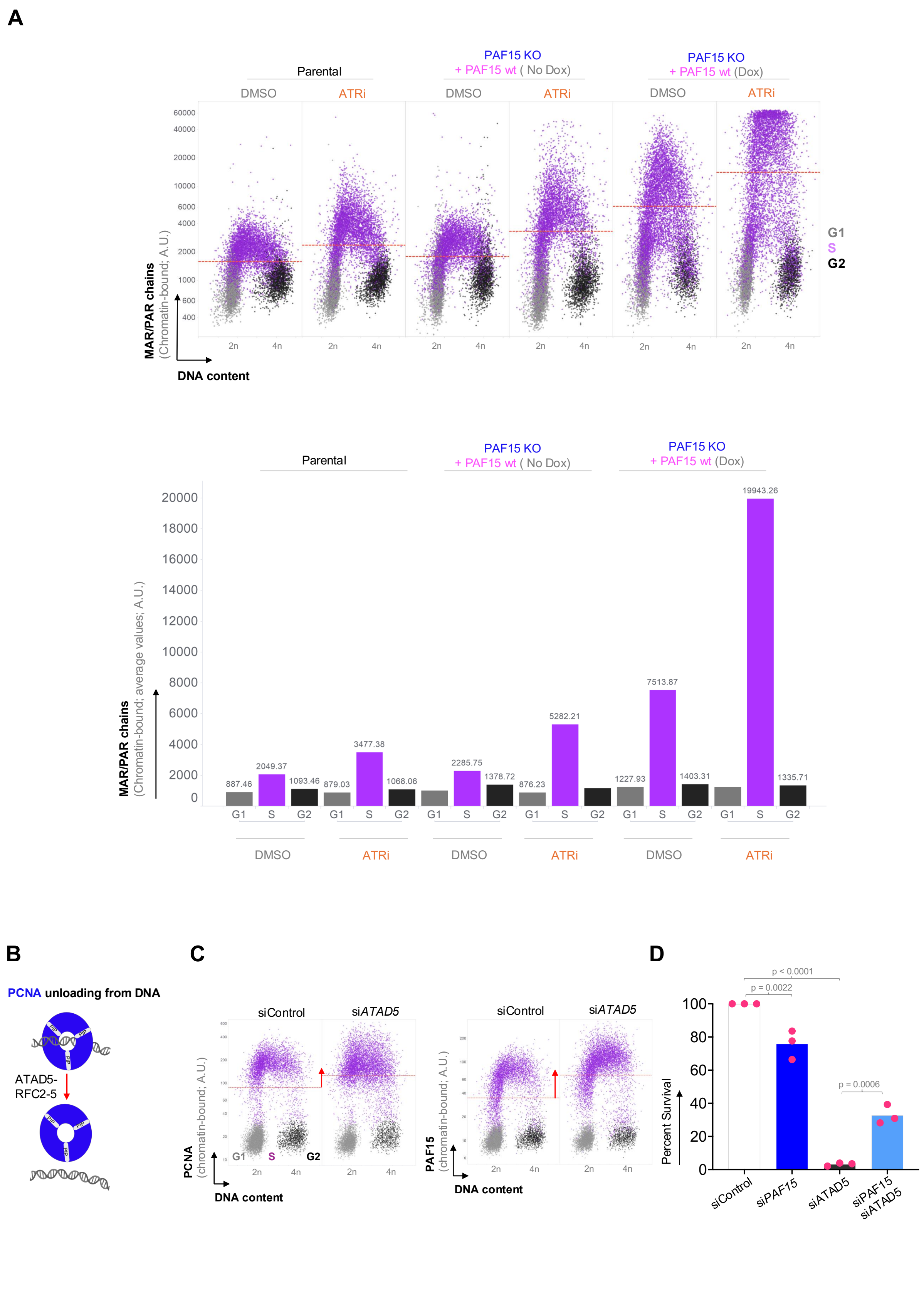
A
Parental
PAF15 KO
+ PAF15 wt ( No Dox)
PAF15 KO
+ PAF15 wt (Dox)
DMSO
ATRi
MAR/PAR chains
(Chromatin-bound; A.U.)
DNA content
2n
4n
G1
S
G2
MAR/PAR chains
(Chromatin-bound; average values; A.U.)
887.46
2049.37
1093.46
879.03
3477.38
1068.06
2285.75
1378.72
876.23
5282.21
1227.93
7513.87
1403.31
19943.26
1335.71
B
PCNA unloading from DNA
ATAD5-
RFC2-5
C
siControl
siATAD5
PCNA
(chromatin-bound; A.U.)
PAF15
(chromatin-bound; A.U.)
DNA content
D
p < 0.0001
p = 0.0022
p = 0.0006
Percent Survival
siControl
siPAF15
siATAD5
siPAF15
siATAD5

**Figure S18**

**Fig. 18. Depletion of PCNA unloader ATAD5 rescues the lethal phenotype of PAF15 overexpression. (A)** (Top) QIBC analysis of chromatin-bound mono- and poly-ADP-ribosylated (MAR/PAR) chains in PAF15 overexpressing U2OS cells treated with PARG and ATR inhibitors. 2n: G1, 4n: G2; > **5**,000 cells per condition**.** (Bottom) Quantification of MAR/PAR chains intensity in different phases of cell cycle gated based on EdU**.** Average values are indicated in different cell cycle phases. **(B)** Schematic visualization of PCNA trimer unloading from DNA. **(C)** QIBC analysis of chromatin fractions of PCNA (Left) and PAF15 (Right) in U2OS cells with ATAD5 depletion. n: G1, 4n: G2; > **5**,000 cells per condition**.** The horizontal line depicts the average values**. (D)** Survival analysis of simultaneous depletion of PAF15 and ATAD5 in U2OS cells. Values denote mean ± s.d. P values were determined by one-way ANOVA with Tukey's test.

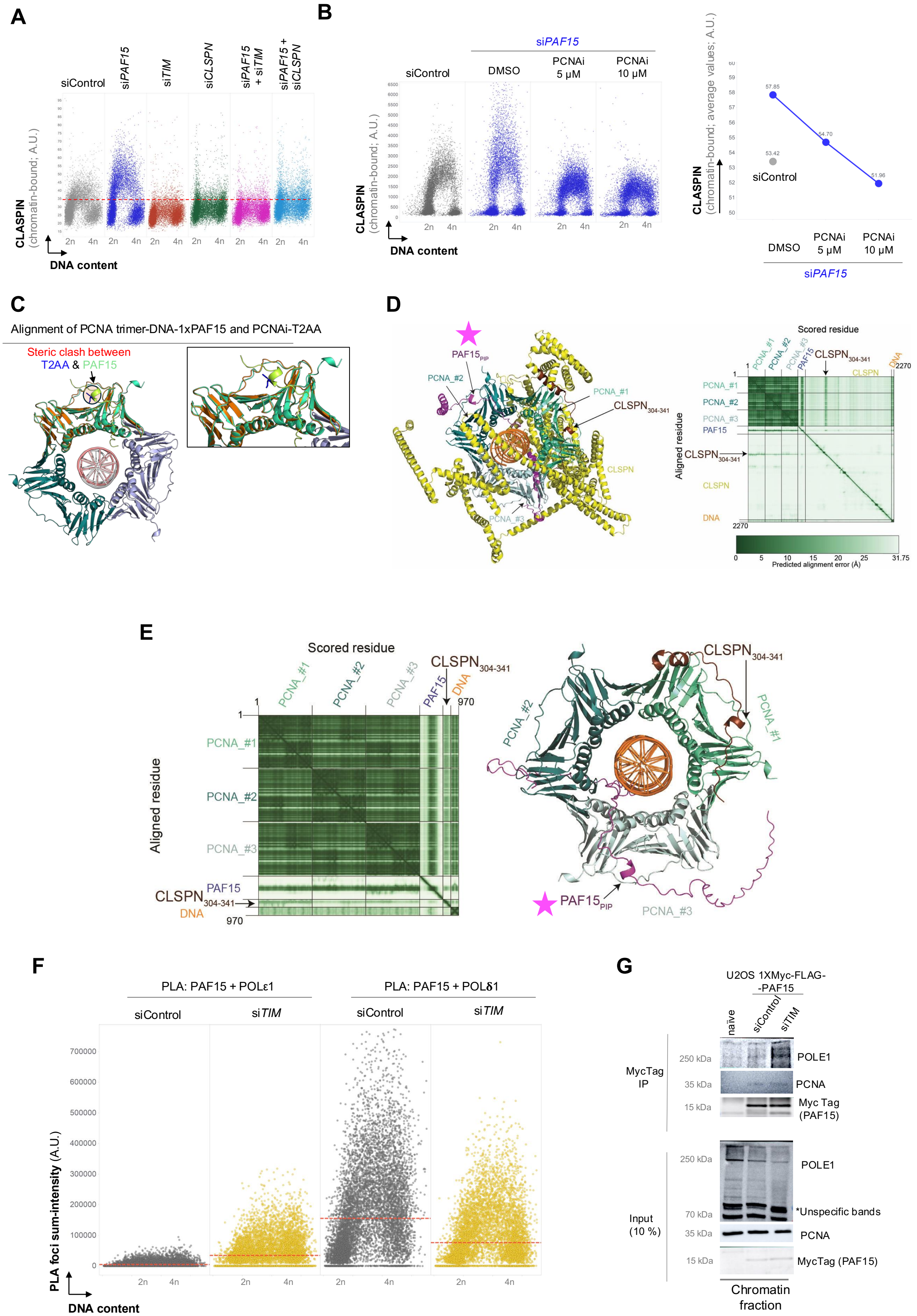
A
siControl
siPAF15
siTIM
siCLSPN
siPAF15 + siTIM
siPAF15 + siCLSPN
CLASPIN (chromatin-bound; A.U.)
DNA content
B
siPAF15
siControl
DMSO
PCNAi 5 µM
PCNAi 10 µM
CLASPIN (chromatin-bound; A.U.)
DNA content
CLASPIN (chromatin-bound; average values; A.U.)
57.85
54.70
51.96
53.42
C
Alignment of PCNA trimer-DNA-1xPAF15 and PCNAi-T2AA
Steric clash between T2AA & PAF15
D
Scored residue
Aligned residue
Predicted alignment error (Å)
E
Scored residue
Aligned residue
F
PLA: PAF15 + POLε1
PLA: PAF15 + POLδ1
siControl
siTIM
PLA foci sum-intensity (A.U.)
DNA content
G
U2OS 1XMyc-FLAG--PAF15
naive
siControl
siTIM
MycTag IP
Input (10 %)
POLE1
PCNA
Myc Tag (PAF15)
*Unspecific bands
Chromatin fraction

Figure S19

**Fig. S19. Inerplay of PAF15 and TIMELESS-CLASPIN for PCNA (A)** QIBC analysis of chromatin-bound CLASPIN in U2OS cells with depletion of indicated proteins. A total of 5,000 cells were analyzed per condition. Red dotted horizontal line in QIBC scatter plots indicates the approximate background signal of CLASPIN on chromatin **(B)** (Left) QIBC analysis of chromatin-bound CLASPIN in U2OS cells with simultaneous depletion of PAF15 and PCNA inhibition (T2AA, 5 and 10 μM). (Right) Average Mean intensity of chromatin-bound CLASPIN staining. **(C)** Modelling of PCNA inhibitor T2AA (PDB: 3WGW) onto the AlphaFold 3 predicted structure of PCNA(trimer)-DNA-PAF15 shows steric clash between PAF15PIP and T2AA **(D)** AlphaFold predicted structural model of three PCNAs in complex with PAF15, CLASPIN (CLSPN) and dsDNA shows high confidence in the folding of PCNA and DNA and very low confidence in the folding of CLSPN. According to the PAE score of the model, the 304-341 residues of claspin that contains the PIP motif (residues 311-318) are in the closest proximity of PCNA, but they are within a predicted alignment error of 20-25Å which can be situated beyond the interaction distance **(E)** CLSPN304-341 is also predicted to form a complex with PCNA (trimer), PAF15 and DNA. The PAE score of the complex indicates that CLSPN304-341 is specifically close to one PCNA molecule (PCNA_#1). **(F)** QIBC analysis of PLA foci of PAF15-POLE1 and PAF15-POLD1 pairs in U2OS cells with depleted TIMELESS (TIM). n > 10,000 cells. The horizontal line depicts the average values for 10,000 cells per condition. **(G)** Immunoprecipitation (IP) analysis of POLE1 in Myc-FLAG-tagged PAF15 from chromatin fraction in U2OS cells under TIMELESS depletion for 30 hours.

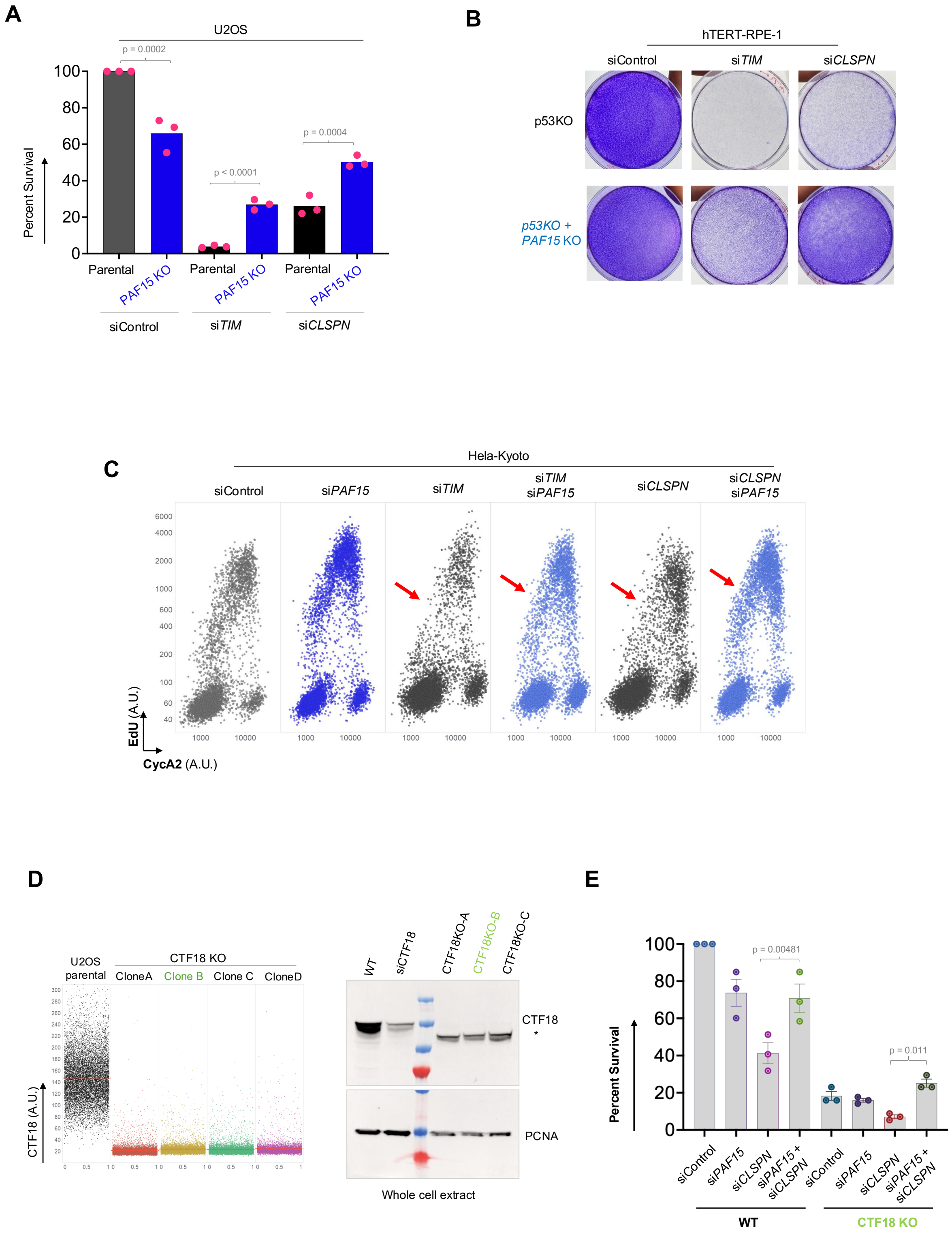
A
U2OS
p = 0.0002
p < 0.0001
p = 0.0004
Percent Survival
Parental
PAF15 KO
siControl
siTIM
siCLSPN
B
hTERT-RPE-1
siControl
siTIM
siCLSPN
p53KO
p53KO + PAF15 KO
C
Hela-Kyoto
siControl
siPAF15
siTIM
siTIM siPAF15
siCLSPN
siCLSPN siPAF15
EdU (A.U.)
CycA2 (A.U.)
D
U2OS parental
CTF18 KO
CloneA
Clone B
Clone C
CloneD
CTF18 (A.U.)
WT
siCTF18
CTF18KO-A
CTF18KO-B
CTF18KO-C
CTF18
PCNA
Whole cell extract
E
p = 0.00481
p = 0.011
Percent Survival
siControl
siPAF15
siCLSPN
siPAF15 + siCLSPN
WT
CTF18 KO

**Figure S20**

**Fig. S20. TIMELESS and CLASPIN shield the CTF18-mediated delivery of PCNA to the leading strand.** (A) Survival analysis of parental and PAF15KO U2OS **(A)** and hTERT-RPE1 **(B)** cells with depletion of TIMELESS and CLASPIN. Values in bar graphs denote mean ± s.d. All the P values were determined by one-way ANOVA with Tukey's test. **(C)** QIBC analysis of Cyclin A2 and EdU in HeLa Kyoto cells with depletion of indicated proteins. A total of 10,000 cells were analyzed per condition. Red arrows indicate S-phase population revival in TIMELESS or CLASPIN depletion together with PAF15. **(D)** QIBC (Left) and Western blot (Right) validation of U2OS CTF18 KO clones and depletion of CTF18 with siRNA. (E) Survival analysis of parental and CTF18KO U2OS cells with depleted PAF15 and CLASPIN. Values in bar graphs denote mean ± s.d. All the P values were determined by one-way ANOVA with Tukey's test.

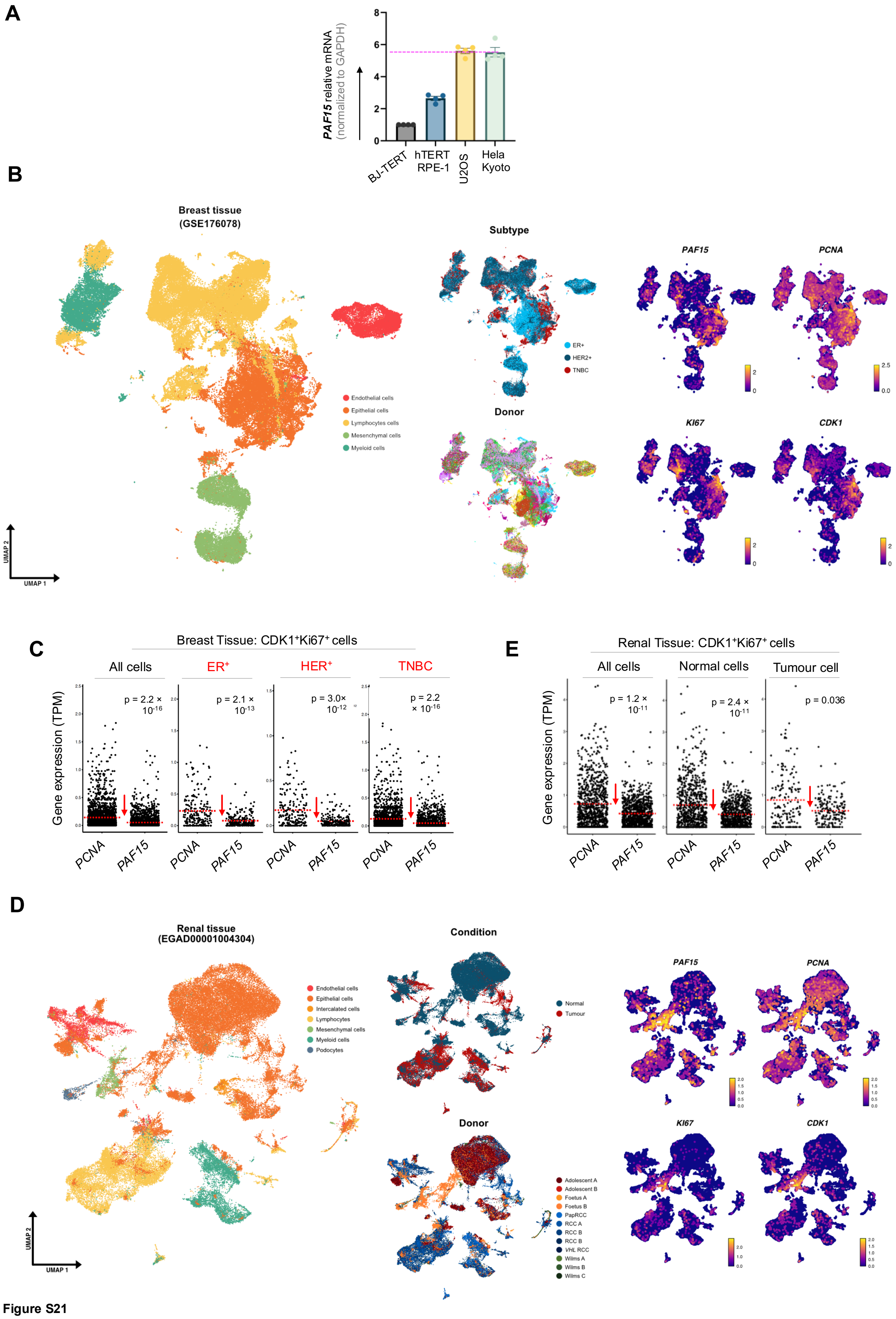
A
PAF15 relative mRNA (normalized to GAPDH)
BJ-TERT
hTERT RPE-1
U2OS
Hela Kyoto
B
Breast tissue (GSE176078)
Endothelial cells
Epithelial cells
Lymphocytes cells
Mesenchymal cells
Myeloid cells
Subtype
ER+
HER2+
TNBC
Donor
PAF15
PCNA
KI67
CDK1
UMAP 1
UMAP 2
C
Breast Tissue: CDK1+Ki67+ cells
All cells
ER+
HER+
TNBC
Gene expression (TPM)
p = 2.2 × 10-16
p = 2.1 × 10-13
p = 3.0× 10-12
p = 2.2 × 10-16
PCNA
PAF15
E
Renal Tissue: CDK1+Ki67+ cells
All cells
Normal cells
Tumour cell
p = 1.2 × 10-11
p = 2.4 × 10-11
p = 0.036
D
Renal tissue (EGAD00001004304)
Endothelial cells
Epithelial cells
Intercalated cells
Lymphocytes
Mesenchymal cells
Myeloid cells
Podocytes
Condition
Normal
Tumour
Donor
Adolescent A
Adolescent B
Foetus A
Foetus B
PapRCC
RCC A
RCC B
RCC B
VHL RCC
Wilms A
Wilms B
Wilms C

**Figure S21**

**Fig. S21. Low PAF15 and PCNA mRNA levels correlate across various cell types in breast and renal tissue. (A)** qPCR analysis of PAF15 mRNA in indicated cell lines. **(B)** Left: UMAP and cell type annotation of cells (n = 100,064) from public repository GSE176078 containing cells from 26 primary tumors from three major clinical subtypes of breast cancer (ER+ = 11, HER2+ = 5, and TNBC = 10). Right: UMAP with cells split by cancer subtype and donor and Log1p gene expression of PAF15, PCNA, KI67, and CDK1. **(C)** Expression of PCNA and PAF15 in CDK1+MKI67+ cells in ER+, HER2+, and TNBC breast cancer subtypes. Red dotted lines denote mean expression values. Expression values are shown as transcripts per million (TPM) with each dot representing a cell. **(D)** Expression of PCNA and PAF15 in CDK1+MKI67+ cells in all cells, normal, and tumor cells from renal cancer. Red dotted lines denote mean expression values. Expression values are shown as transcripts per million (TPM) with each dot representing a cell. **(E)** Left: UMAP and cell type annotation of cells (n = 68,977) from European Genome-Phenome Archive study EGAS00001002325 containing cells from normal renal tissue (fetal = 2, pediatric = 3, adolescent = 2, and adult = 5) and clinical renal cancer types (Wilms tumor = 3, clear cell renal cell carcinoma (ccRCC) = 3, papillary renal cell carcinoma (pRCC) = 1). Right: UMAP with cells split by condition and donor and Log1p gene expression of PAF15, PCNA, KI67, and CDK1. P values were determined by Wilcoxon signed-rank test.

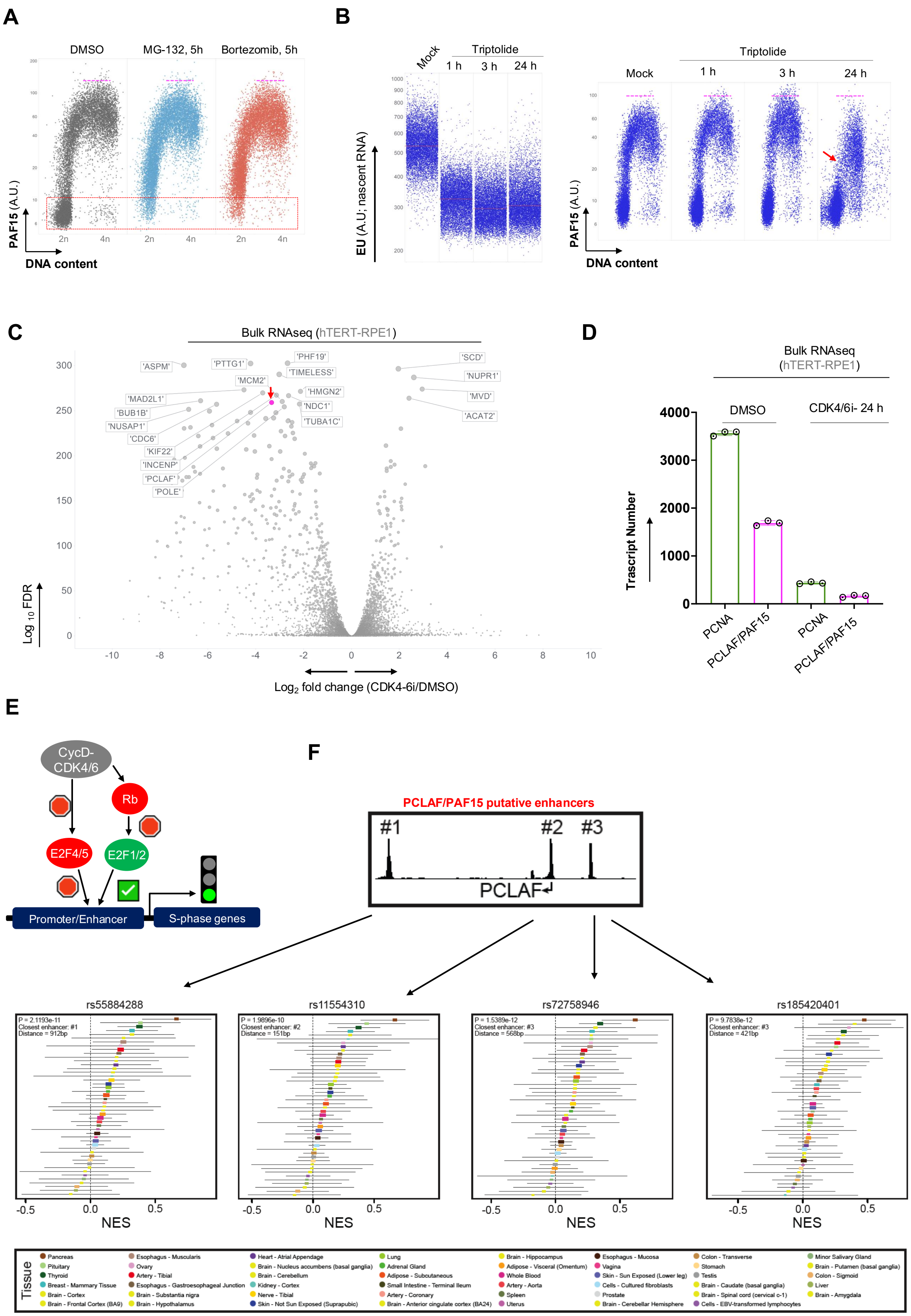

A
DMSO
MG-132, 5h
Bortezomib, 5h
PAF15 (A.U.)
DNA content
2n
4n
B
Mock
Triptolide
1 h
3 h
24 h
EU (A.U; nascent RNA)
DNA content
C
Bulk RNAseq (hTERT-RPE1)
'ASPM'
'PTTG1'
'PHF19'
'TIMELESS'
'MCM2'
'HMGN2'
'NDC1'
'TUBA1C'
'MAD2L1'
'BUB1B'
'NUSAP1'
'CDC6'
'KIF22'
'INCENP'
'PCLAF'
'POLE'
'SCD'
'NUPR1'
'MVD'
'ACAT2'
Log 10 FDR
Log2 fold change (CDK4-6i/DMSO)
D
Bulk RNAseq (hTERT-RPE1)
DMSO
CDK4/6i- 24 h
Trascript Number
PCNA
PCLAF/PAF15
E
CycD-CDK4/6
Rb
E2F4/5
E2F1/2
Promoter/Enhancer
S-phase genes
F
PCLAF/PAF15 putative enhancers
#1
#2
#3
PCLAF
rs55884288
rs11554310
rs72758946
rs185420401
NES
Tissue


Figure S22

**Fig. S22. PAF15 expression is regulated by the E2F family of transcription factors. (A)** QIBC analysis of PAF15 levels in U2OS cells treated with proteasome inhibitors MG-132 and Bortezomib for 5 h. n > 10,000 cells. Rectangular red box depicts that PAF15 is stabilized in G1 cell population upon proteasome inhibition. Pink dotted lines in QIBC scatter plots indicate the approximate maximum levels of each protein. **(B)** QIBC analysis of EU (nascent RNA) and PAF15 levels in U2OS cells treated with Triptolide for 1, 3 and 24 hours. The horizontal line in EU QIBC plot depicts the average values for 10,000 cells per condition. In the QIBC plot of PAF15, pink dotted lines in QIBC scatter plots indicate the approximate maximum levels of each protein and red arrow shows decline of PAF15 upon long-term Triptolide treatment. **(C)** Bulk RNAseq analysis of hTERT-RPE1 treated with CDK4/6 inhibitor (24 h) showing significant downregulation of PAF15 (PCLAF, red arrow). **(D)** Transcript numbers of PAF15 and PCNA in DMSO and CDK4/6 inhibitor treated cells, from data in panel C. **(E)** Schematic visualization of the regulatory pathway of the transcription of S-phase proteins. E2F family of transcription factors is the main coordinator of G1S transition and consists of activator (E2F1/2) as well as repressor (E2F4/5) genes. **(F)** Mapping of expression quantitative trait loci (eQTL) single nucleotide polymorphisms (SNPs) across multiple tissue types in Activity-By-Contact (ABC)-modelled PAF15/ PCLAF causal enhancers (for more information please see Figure 5D).

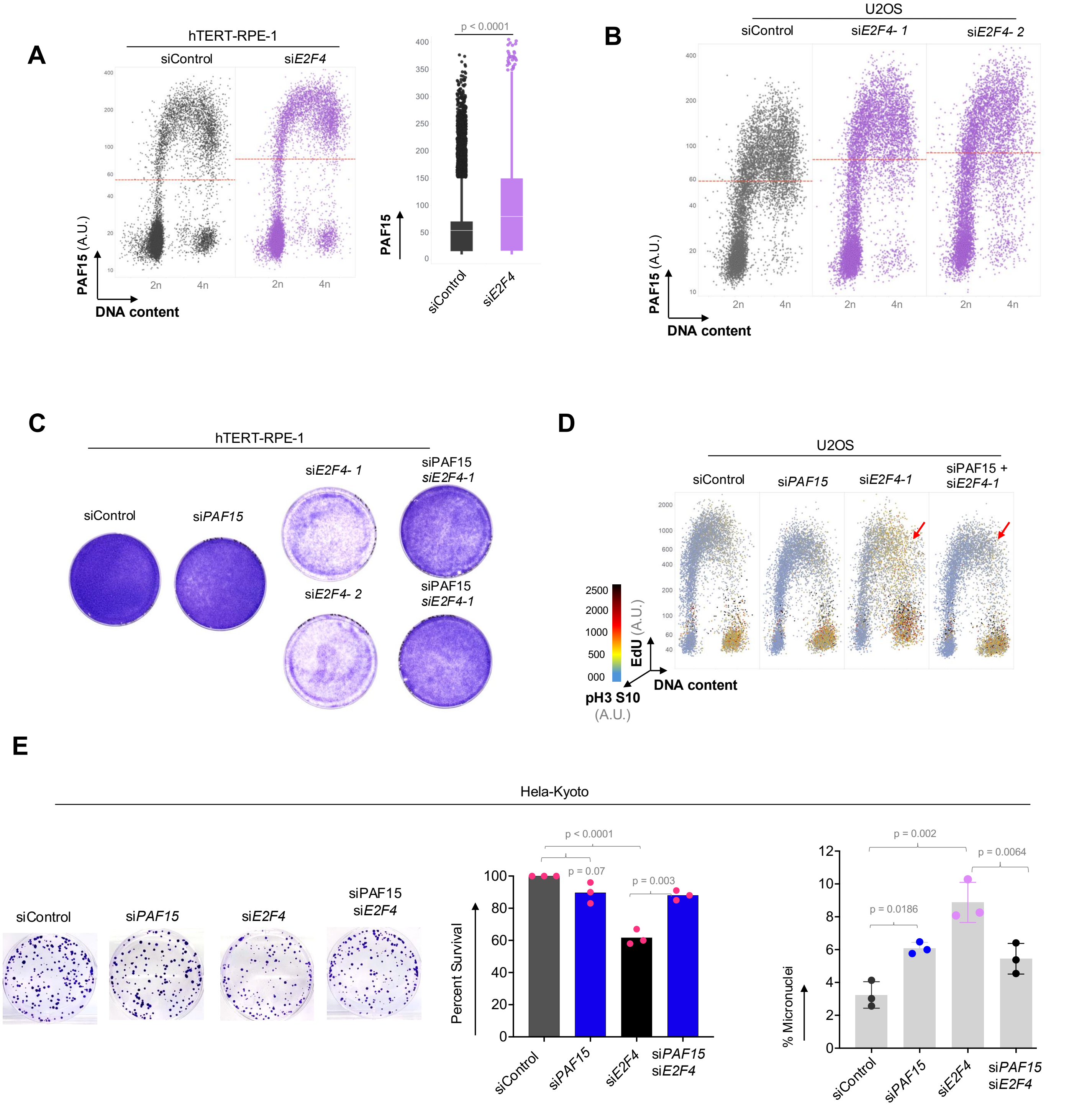
A
hTERT-RPE-1
siControl
siE2F4
PAF15 (A.U.)
DNA content
2n
4n
p < 0.0001
PAF15
siControl
siE2F4
B
U2OS
siControl
siE2F4- 1
siE2F4- 2
PAF15 (A.U.)
DNA content
C
hTERT-RPE-1
siControl
siPAF15
siE2F4- 1
siPAF15 siE2F4-1
siE2F4- 2
siPAF15 siE2F4-1
D
U2OS
siControl
siPAF15
siE2F4-1
siPAF15 + siE2F4-1
EdU (A.U.)
DNA content
pH3 S10 (A.U.)
E
Hela-Kyoto
siControl
siPAF15
siE2F4
siPAF15 siE2F4
Percent Survival
p < 0.0001
p = 0.07
p = 0.003
% Micronuclei
p = 0.002
p = 0.0064
p = 0.0186

Figure S23

**Fig. S23. E2F4 depletion increases PAF15 levels, leading to compromised cell survival rescued by PAF15 depletion.** **(A)** QIBC analysis of PAF15 levels in hTERT-RPE-1 with depleted E2F4 transcription factor. The horizontal line depicts the average values for 10,000 cells per condition. **(B)** QIBC analysis of PAF15 levels in U2OS with depleted E2F4 transcription factor with two distinct siRNAs. The horizontal line depicts the average values for 10,000 cells per condition. **(C)** Survival analysis of simultaneous depletion of PAF15 and E2F4 in hTERT-RPE-1. **(D)** QIBC analysis of S-phase and mitotic fractions of U2OS cells, as indicated by EdU and phosphorylation of Serine 10 on Histone H3 (pH3 S10) respectively, treated by siRNAs targeting PAF15 and E2F4. n > 10,000 cells. **(E)** (Left) Representative images of survival assay on Hela-Kyoto cells depleted for PAF15 and E2F4. (Middle) Quantification of the number of colonies formed during survival assay on U2OS cells depleted for PAF15 and E2F4. (Right) Quantification of micronuclei formation in U2OS cells depleted for PAF15 and E2F4. Values in bar graphs denote mean ± s.d. All the P values were determined by one-way ANOVA with Tukey's test.

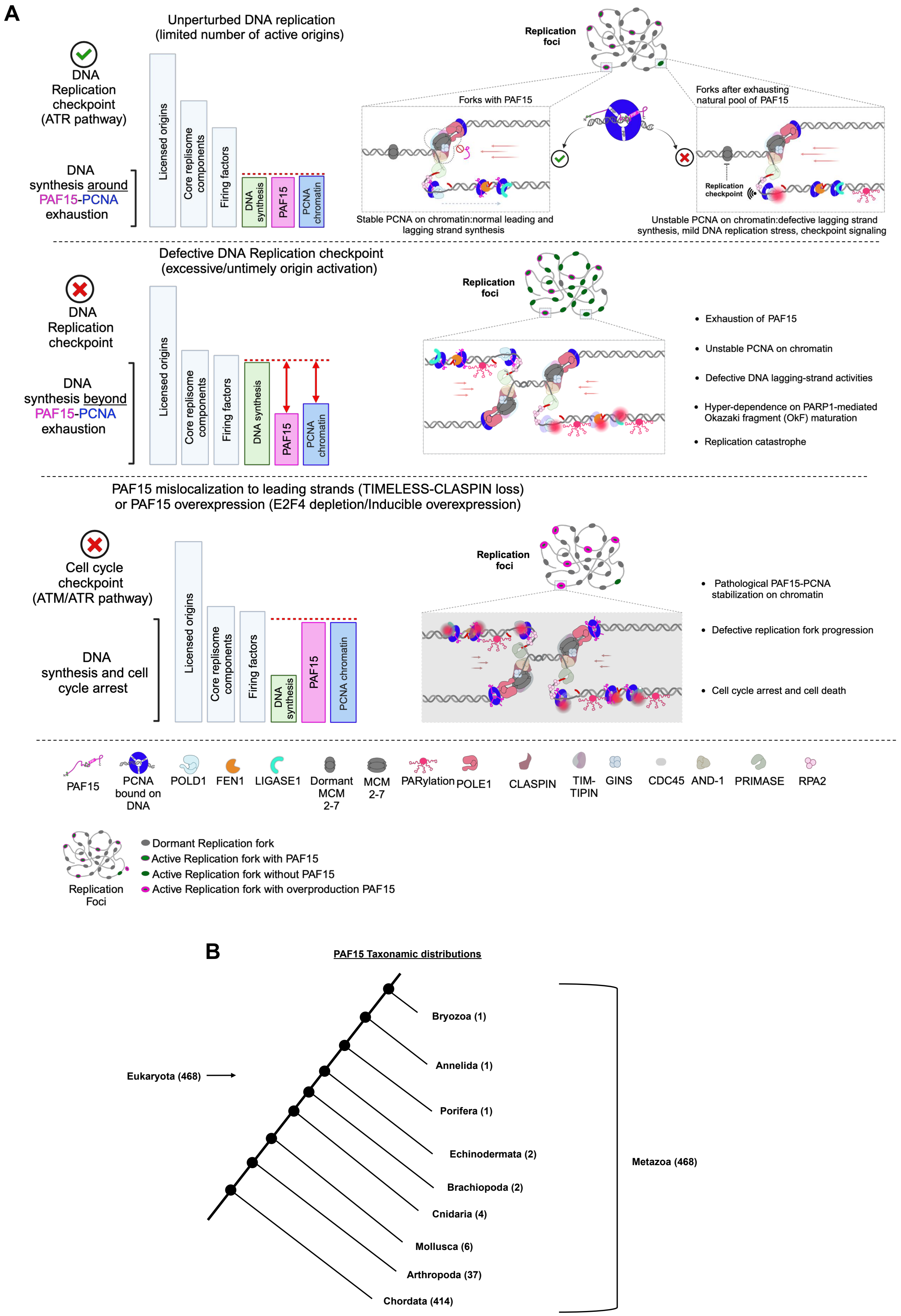
A
Unperturbed DNA replication
(limited number of active origins)
DNA Replication checkpoint (ATR pathway)
DNA synthesis around PAF15-PCNA exhaustion
Licensed origins
Core replisome components
Firing factors
DNA synthesis
PAF15
PCNA chromatin
Replication foci
Forks with PAF15
Forks after exhausting natural pool of PAF15
Replication checkpoint
Stable PCNA on chromatin:normal leading and lagging strand synthesis
Unstable PCNA on chromatin:defective lagging strand synthesis, mild DNA replication stress, checkpoint signaling
Defective DNA Replication checkpoint
(excessive/untimely origin activation)
DNA Replication checkpoint
DNA synthesis beyond PAF15-PCNA exhaustion
Licensed origins
Core replisome components
Firing factors
DNA synthesis
PAF15
PCNA chromatin
Replication foci
Exhaustion of PAF15
Unstable PCNA on chromatin
Defective DNA lagging-strand activities
Hyper-dependence on PARP1-mediated Okazaki fragment (OkF) maturation
Replication catastrophe
PAF15 mislocalization to leading strands (TIMELESS-CLASPIN loss)
or PAF15 overexpression (E2F4 depletion/Inducible overexpression)
Cell cycle checkpoint (ATM/ATR pathway)
DNA synthesis and cell cycle arrest
Licensed origins
Core replisome components
Firing factors
DNA synthesis
PAF15
PCNA chromatin
Replication foci
Pathological PAF15-PCNA stabilization on chromatin
Defective replication fork progression
Cell cycle arrest and cell death
PAF15
PCNA bound on DNA
POLD1
FEN1
LIGASE1
Dormant MCM 2-7
MCM 2-7
PARylation
POLE1
CLASPIN
TIM-TIPIN
GINS
CDC45
AND-1
PRIMASE
RPA2
Replication Foci
Dormant Replication fork
Active Replication fork with PAF15
Active Replication fork without PAF15
Active Replication fork with overproduction PAF15
B
PAF15 Taxonamic distributions
Eukaryota (468)
Bryozoa (1)
Annelida (1)
Porifera (1)
Echinodermata (2)
Brachiopoda (2)
Cnidaria (4)
Mollusca (6)
Arthropoda (37)
Chordata (414)
Metazoa (468)

**Figure S24**

**Fig. S24. A model of PAF15-dependent rate-limiting mechanism for PCNA-lagging strand processing during unperturbed DNA synthesis. (A)** Under normal conditions, stochastic origin firing and the limited availability of the PAF15 pool and chromatin-bound PCNA serve as rate-limiting factors for DNA replication. The dynamic loading of the PAF15–PCNA complex defines the overall range of active DNA synthesis. When the natural PAF15 reservoir is exhausted, PCNA-dependent lagging strand processing is compromised, potentially triggering ATR-mediated checkpoint activation. Middle: With defective DNA replication checkpoint, the rate of DNA synthesis exceeds PAF15 levels, leading to destabilization of lagging strand PCNA on new origins. As a result, the processing of lagging strand becomes unstable, leading to dependence on non-canonical PARP1-mediated pathway of Okazaki fragment maturation, eventually resulting in replication catastrophe. Bottom: PAF15 should be strictly confined to the lagging strands and rate-limited during unperturbed DNA replication. This conclusion stems from observations that misregulation of PAF15 severely impacts DNA replication, potentially leading to lethal consequences. In cells with TIMELESS and Claspin depletion, PAF15 localizes to the leading strands, where it obstructs PCNA activities. Similarly, when PAF15 is overexpressed—either via ectopic expression or loss of E2F4 control—it leads to excessive chromatin loading of PAF15, which in turn completely disrupts PCNA functions. Ultimately, these disruptions cause significant replication fork slowing, blockade of checkpoint signaling, and cell death. See text for more details. **(B)** A simplified version of the PAF15 taxonomic/phylogenetic tree within Eukaryota was generated by comparing a total of 468 Metazoan species. In this tree, Metazoa serve as the root, and each node represents a branching point within the eukaryotic lineage. Branches terminate at nodes that are organized based on the number of species they contain. Within the Metazoa group, Bryozoa, Annelida, and Porifera have the fewest species (one each), whereas Chordata boasts the highest number (414 species). Protein sequence alignment analysis was performed using EMBL-EBI software packages to compare sequence similarities across various species. Notably, protein sequence similarities were observed only in higher eukaryotes, while lower species in the evolutionary tree did not exhibit such similarities. Expasy and Phylo.io were employed to generate the phylogenetic tree and produce its simplified version based on the output data.
